# HyphAeon: Attention on Evolution Across Deep Time Transforms Comparative Genomics

**DOI:** 10.64898/2026.09.06.749597

**Authors:** Sergei L. Kosakovsky Pond, Steven Weaver, Danielle Callan, Jordan D. Zehr, Alexander G. Lucaci, Hannah Verdonk, Avery Selberg, Gallean Brown, Maria Chikina, Nathan L. Clark, Kateryna D. Makova, Darren P. Martin, Anton Nekrutenko

## Abstract

Detecting Darwinian natural selection is fundamental to evolutionary biology and functional genomics, yet standard methods based on phylogenetic models that estimate the ratio of non-synonymous to synonymous substitution rates (*dN/dS*) fail to scale with modern genomic volumes. Fitting continuous-time Markov substitution matrices across dense trees with hundreds of species requires extensive compute, forcing comparative genomics to rely on aggressive taxon subsampling or static whole-tree summaries that dilute transient adaptive bursts. Here we present HyphAeon, a lightweight (~1.91M parameter backbone, 2.46M across the full multi-task suite) phylogeny-informed foundation transformer trained to amortize the detection of episodic diversifying selection across 742-species mammalian coding alignments (17,186 genes, 9.77 *×* 10^6^ codons). HyphAeon approaches the discriminative accuracy of numerical maximum-likelihood selection tests (MEME) across episodic burst regimes (ROC-AUC up to 0.942, mean 0.659; empirical Precision-Recall lift up to 25.8×, mean 7.7*×*; rank concordance up to *ρ* = 0.983) while executing *>* 1,000*×* faster per locus (averaging *<* 1 ms per site across genome-wide scans) and *>* 10,000*×* faster at proteome scale, generalizing outside its mammalian training distribution without retraining. Beyond accelerating classical tests, embedding molecular evolution into a differentiable geometric latent space enables analytical capabilities inaccessible to static *dN/dS* models: (1) targeted alignment artifact correction via counterfactual attribution; (2) macromolecular contact recovery and multi-site epistatic sectors (CESI); (3) directional phenotype-to-genotype attribution in lineage space (PARS); and (4) continuous temporal surveillance regression that tracks positive sweep velocities across longitudinal cohorts (evaluated across 12,167 timestamped genomes and benchmarked against external frequencies from *>* 9.34 million genomes), rescuing adaptive substitutions obscured by post-fixation dilution. By bridging statistical phylogenetics with geometric representation learning, HyphAeon establishes comparative genomics as an interactive, high-throughput computational framework for evolutionary discovery.

## 1 Introduction

Over the past three decades, statistical phylogenetics has provided the mathematical foundation for detecting Darwinian natural selection from comparative sequence alignments [1–5]. Codon-based continuous-time Markov substitution models that estimate the ratio of non-synonymous to synonymous substitution rates (*ω* = *dN/dS*)—exemplified by workhorse maximum-likelihood methods such as MEME [6] and FEL [4]—have long chronicled molecular arms races, charting everything from viral immune evasion to deep macroevolutionary divergence. However, standard statistical phylogenetics is failing to keep pace with the explosive growth of modern genomics. Global initiatives like the Vertebrate Genomes Project (VGP) [7], Zoonomia [8], TOGA [9], and the Earth BioGenome Project [10] are producing chromosome-level assemblies for thousands of taxa. Yet the computational cost of numerical likelihood optimization scales poorly with modern genomic volumes: repeatedly tuning continuous-time substitution matrices across dense trees with hundreds of species demands tens of thousands of CPU core-hours for proteome-wide comparative scans [5] (in contrast to a one-time pre-training investment of *<* 24 hours on a single accelerator node for an amortized neural surrogate).

A fundamental epistemological challenge in evolutionary genomics is that what is and is not positively selected in natural sequences is almost always inherently unobservable. Because historical selective sweeps leave no definitive ground-truth ledger in natural populations, the field must triangulate against external empirical anchors—clinically validated drug-resistance mutations, experimental deep mutational scanning (DMS) fitness landscapes, macromolecular structural constraints, and phenotypic convergence—to corroborate or refute statistical inferences.

Compounding this computational bottleneck, large-scale genomic datasets present substantial methodological challenges. In draft assemblies and automated alignment pipelines, sequencing errors and unannotated indels can cause single lineages to fall out of frame, generating dense runs of spurious non-synonymous substitutions along terminal branches [11, 12]. Because standard phylogenetic models assume independent and identically distributed (i.i.d.) substitutions without spatial or assembly error awareness, they readily mistake a perfunctory sequencing glitch or frame-shift stumble for genuine Darwinian adaptation. Furthermore, while deep learning has advanced structural biology and clinical variant classification [13–16], comparative genomics has lagged behind. Previous neural surrogates trained on synthetic Markov alignments [17–19] enforce simplified, site-independent assumptions that ignore 3D physical packing and epistatic fitness landscapes. Conversely, self-supervised protein language models (pLMs) and MSA transformers [20–22] treat sequences as unstructured strings or uniform row batches with 1D positional encodings, explicitly discarding continuous divergence times, tree branch lengths, and the fundamental biophysical distinction between synonymous (*dS*) and non-synonymous (*dN*) mutations.

Beyond deep-time macroevolutionary species trees, comparative genomics faces an acute temporal challenge in high-frequency longitudinal pathogen surveillance. Platforms such as Nextstrain [23,24] routinely ingest tens of thousands of viral and bacterial genomes sampled across calendar weeks and months. In this high-frequency regime, classical phylogenetics encounters both theoretical and computational ceilings. Static tree-wide *dN/dS* scans suffer from post-fixation dilution: an intense adaptive burst is diluted below statistical detection thresholds once an advantageous variant sweeps to fixation across circulating clades [25]. Conversely, heuristic sliding time windows artificially fragment continuous selective surges and suffer from severe sample starvation within narrow temporal slices [26–29], while Bayesian tip-dated phylodynamic sampling (e.g., BEAST [30–33]) incurs steep computational scaling, rendering dense real-time surveillance across thousands of contemporary genomes practically intractable.

Here, we show that embedding phylogenetic tree geometry directly into transformer attention sidesteps numerical optimization bottlenecks while opening new analytical capabilities. We present HyphAeon, a lightweight (~ 1.91M parameter) phylogenetic transformer built upon an axial attention backbone. HyphAeon couples discrete codon and amino acid tokenizations with continuous-time substitution priors and tree-geometric rotary position embeddings (Tree-RoPE; Methods). Because these coordinates depend strictly on pairwise evolutionary divergence rather than a fixed branching graph, HyphAeon can accept user-provided phylogenetic trees or operate completely tree-free by estimating pairwise distances directly from sequence alignments. Pre-trained across 17,186 mammalian coding alignments (742 species, 9.77 *×* 10^6^ codons, paired with subtrees pruned from the reference mammalian species tree; Methods), HyphAeon embeds evolutionary history into a differentiable geometric latent space. By learning the continuous landscape of molecular evolution across millions of codons, the model acts as an implicit Empirical Bayes regularizer, shrinking unconstrained likelihood fluctuations toward biophysically viable substitution trajectories and avoiding the stochastic overfitting of single-site maximum likelihood on shallow alignments.

This unified representation establishes an agile computational engine for evolutionary discovery across both macroevolutionary species radiations and ongoing outbreak surveillance. Beyond achieving sub-millisecond selection inference (*>* 1,000*×* faster per locus and *>* 10,000*×* faster at proteome scale than HyPhy MEME/FEL) and excising localized sequencing glitches via counterfactual attribution, HyphAeon unlocks four core downstream capabilities without task-specific retraining: (1) targeted alignment artifact correction and episodic positive selection screening (Pillar 1); (2) macromolecular contact recovery and epistatic sector decomposition (CESI; Pillar 2); (3) directional phenotype-to-genotype attribution in lineage space (PARS [34,35]; Pillar 3); and (4) continuous temporal surveillance regression that tracks positive sweep velocities and collective dynamic factor modes across longitudinal viral and bacterial transmission chains, rescuing authentic episodic sweeps obscured by post-fixation dilution (Pillar 4). Shipped as an open-source package (`hyphaeon`), HyphAeon transforms comparative genomics into an interactive, high-throughput platform for evolutionary hypothesis testing.

## 2 Results

HyphAeon is a foundation model designed to evaluate molecular evolution directly from multiple sequence alignments paired with phylogenetic context (either user-provided Newick trees, automatically estimated fast topologies, or empirical pairwise distance matrices without an explicit tree). Unlike protein language models that treat sequences as isolated strings or unaligned token sets, HyphAeon models the dual geometry of comparative genomics: an alignment of *L* codon sites across *M* species lineages paired with a continuous metric tree (*S, d*_*T*_) or direct pairwise genetic distance matrix **D** ∈ ℝ^*M×M*^. By projecting continuous patristic or pairwise distances into 4-dimensional Classical Multidimensional Scaling (MDS) coordinates and modulating cross-species attention with a continuous-time Markov substitution transition kernel 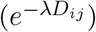, the model internalizes evolutionary divergence directly in continuous metric space, providing natural robustness against minor tree estimation noise and incomplete lineage sorting (ILS).

The backbone is a 6-layer species transformer comprising 1,909,404 trainable parameters. To prevent synonymous rate variation (SRV) and GC-biased gene conversion (gBGC) from confounding selection inference, HyphAeon enforces dual-track representation learning via block-diagonal linear projections (`BlockLinear`), strictly isolating neutral synonymous codon baselines (*dS*, 192 dimensions) from non-synonymous property selection (*dN*, 192 dimensions). Unsupervised pre-training organically recovers the biochemical grammar of the genetic code: stop codons collapse to an identical angular ray (cosine similarity = 1.000) while remaining orthogonal to sense codons, with principal axes across the 64-codon vocabulary capturing GC3 composition (*r* = +0.288, *p* = 0.021), purine skew (*r* = +0.258, *p* = 0.039), and translated residue steric volume (*r* = −0.523, *p* = 9.4 *×* 10^−6^, *N* = 64). Information across all species is compressed into a dedicated `[ROOT]` token at tree origin (0, 0, 0, 0), yielding a 384-dimensional site representation (**h**_root_ ∈ ℝ^384^). Pre-trained on 17,186 mammalian alignments from TOGA (9.77 *×* 10^6^ codons across 742 species), HyphAeon deploys this unified representation across four core evolutionary tasks: (1) episodic selection screening and automated alignment error filtering (MEME/FEL); (2) macromolecular contact recovery and epistatic co-selection networks (CESI); (3) directional phenotype–genotype attribution (PARS); and (4) continuous longitudinal temporal surveillance tracking active selective sweep velocities.

### 2.1 Ultra-Fast Identification of Codon Sites Under Episodic Diversifying Selection

#### 2.1.1 Out-of-Distribution Case Study: HIV-1 Reverse Transcriptase

To evaluate performance under extreme domain shift and establish interpretability baselines, we examined 475 HIV-1 Reverse Transcriptase (RT) sequences (*L* = 335 codons) from a South African clinical cohort following single-dose nevirapine (sdNVP) exposure [36] (Fig. 2). All predictions were evaluated zero-shot using the frozen base mammalian model (`hyphaeon-base-mammal`) without fine-tuning. This dataset represents acute intra-host viral evolution, dense clinical sampling, transient neutral polymorphisms, and 29.8% N-terminal Sanger primer missing data (codons 1–34)—in sharp contrast to the macroevolutionary speciation regime of the mammalian training corpus (as described in Methods, users seeking further viral calibration can optionally invoke the domain-adapted model `hyphaeon-viral` checkpoint).

**Figure 1:**
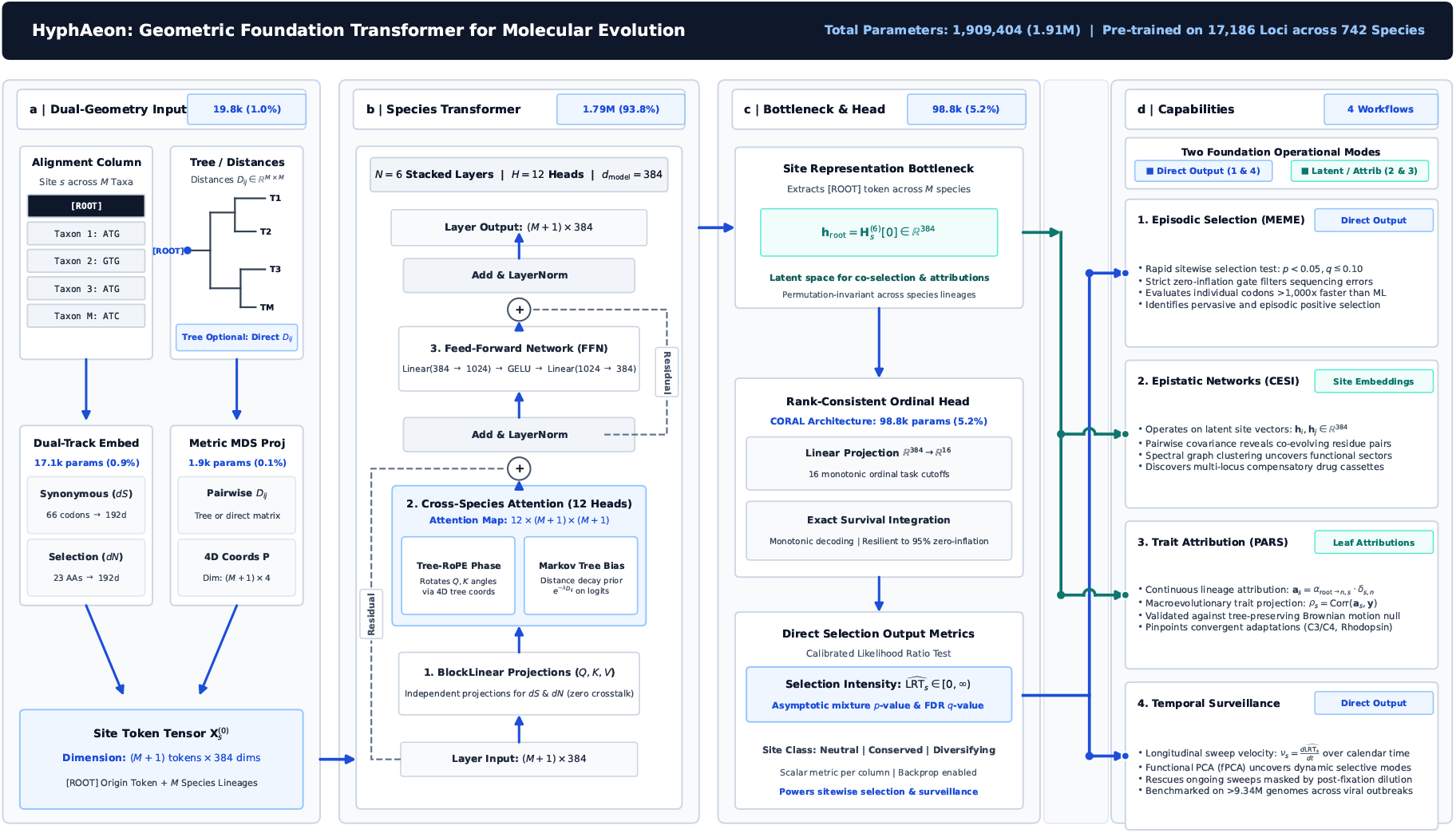
HyphAeon architecture, information flow, and parameter distribution (1.91M parameters). (A) Dual-geometry inputs and embeddings (19.8k params, 1.0%): Individual codon alignment columns across *M* species lineages are paired with a continuous metric tree (*S, d*_*T*_) or directly with an empirical pairwise distance matrix (an explicit tree topology is optional; continuous pairwise distances can be directly passed to the 4D Classical MDS embedding). Synonymous (*dS*) and non-synonymous (*dN*) information are partitioned into independent 192-dimensional tracks (384 dimensions total) via `BlockLinear` projections to insulate positive selection inference from synonymous rate variation, while pairwise distances are projected into 4D Classical MDS coordinates (**P**). (B) Phylo-Species Transformer backbone (1.79M params, 93.8%): *N* = 6 stacked transformer layers with *H* = 12 heads operate strictly across the species axis (*M×M*). Query and Key tensors undergo 4D Tree-RoPE geometric phase rotations, and attention logits incorporate a continuous-time Markov distance-decay prior 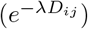, followed by block-diagonal linear projections, Add and LayerNorm, and learnable initial-representation skip connections 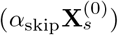. (C) Site bottleneck and ordinal output head (98.8k params, 5.2%): The transformer compresses the full phylogenetic column into a 384-dimensional permutation-invariant site representation 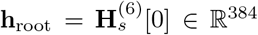. A 16-threshold Rank-Consistent Ordinal Regression (`CORAL`) head decodes selection intensity 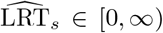 via exact log-space survival integration, maintaining calibration and resilience against 95% neutral zero-inflation. (D) Downstream capabilities operating across two foundation modes: The foundation model powers four biological workflows via two distinct representation tiers: *Direct Output* (blue routing) evaluates scalar 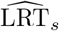 directly for rapid episodic positive selection screening (Pillar 1, MEME) and longitudinal temporal surveillance regression (Pillar 4); *Latent Embeddings and Attributions* (teal routing) utilize site representation geometry **h**_root_ and backpropagated species leaf gradients **a**_*s*_ ∈ ℝ^*M*^ for pairwise epistatic co-selection networks (Pillar 2, CESI) and directional phenotype–genotype attribution (Pillar 3, PARS).

**Figure 2:**
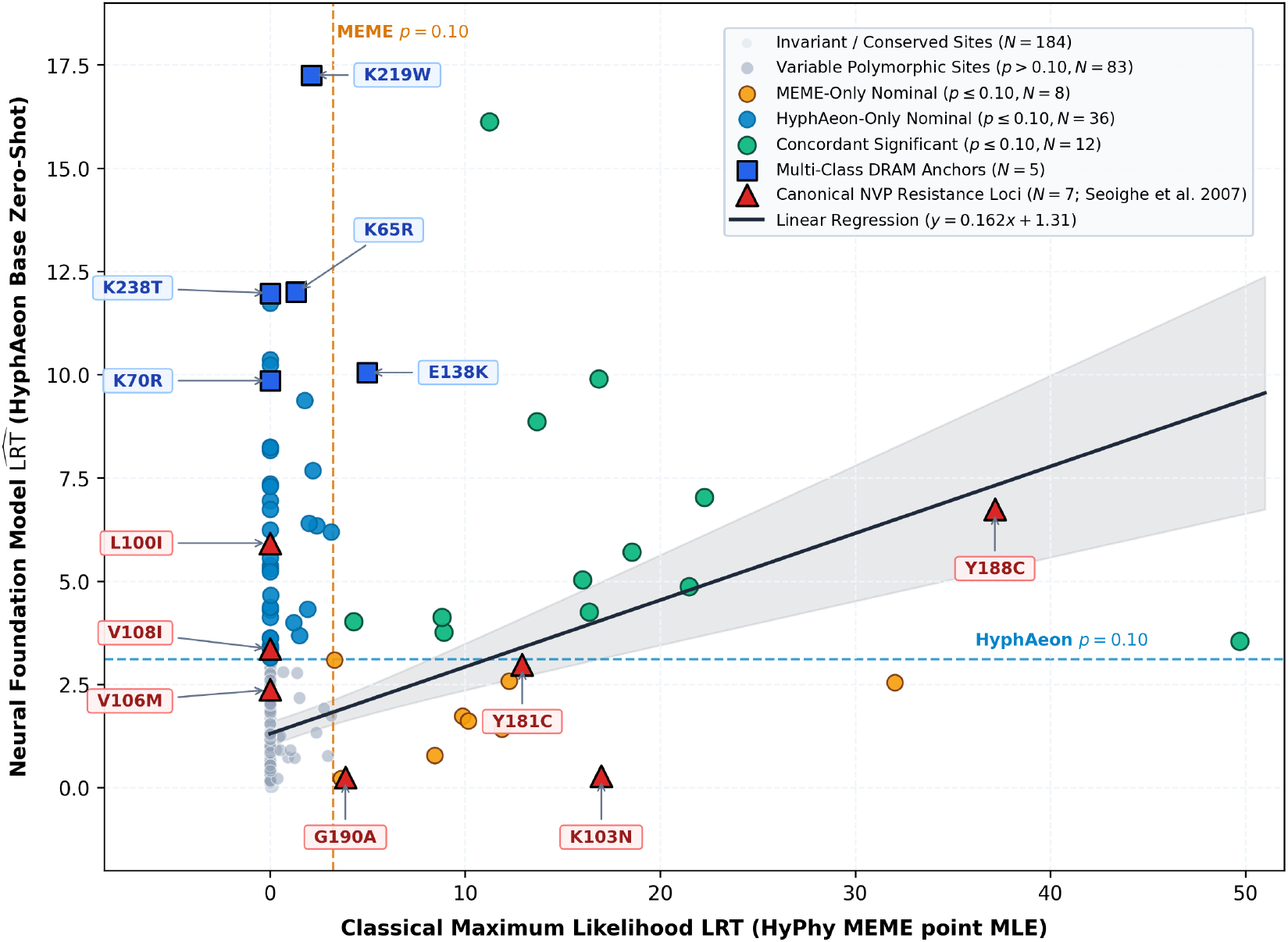
Zero-shot selection profiling and concordance in HIV-1 Reverse Transcriptase (*N* = 475 patient isolates, *L* = 335 codons) evaluated using the frozen base mammalian model. Sitewise concordance between classical numerical HyPhy MEME (LRT point MLE, x-axis) and neural foundation model inference (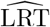, base model evaluated zero-shot in 4.1 seconds on GPU, y-axis) across all 335 codons (*ρ* = 0.528, *p* = 1.96 *×* 10^−25^; log-linear *r* = 0.433; linear regression fit *y* = 0.162*x* + 1.31, solid line with 95% CI). Dashed lines denote nominal significance thresholds (*p* ≤ 0.10). Invariant (*N* = 184, light gray) and variable non-significant (*p >* 0.10, *N* = 125, slate) sites establish strong baseline concordance alongside nominally significant positions (*N* = 26 under numerical MEME). Across 37 sequenced Drug Resistance-Associated Mutation (DRAM) positions (Stanford HIVdb / IAS-USA), HyphAeon identifies 20 loci at *p* ≤ 0.10 (54.1% sensitivity) vs. 11 loci (29.7%) for MEME (AUROC = 0.849 vs. 0.683; AUPRC = 0.418 vs. 0.321 over 0.110 baseline). Canonical nevirapine (NVP) resistance loci [36] (L100I, K103N, V106M, V108I, Y181C, Y188C, G190A; crimson triangles) and multi-class NRTI/NNRTI resistance anchors (K65R, K70R, E138K, K219W, K238T; blue squares) highlight the complementary sensitivity of empirical Bayes neural regularization compared to unconstrained single-site maximum likelihood.

HyphAeon screened all 335 codons in 4.1 seconds on GPU, compared to 2.4 CPU hours for numerical HyPhy MEME (*>* 2,100*×* speedup). Across all 335 codons, predicted 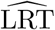 exhibited strong rank concordance with numerical MEME (*ρ* = 0.528, *p* = 1.96 *×* 10^−25^; log-linear *r* = 0.433, *p* = 8.93 *×* 10^−17^; Fig. 2), achieving AUROC = 0.827 and AUPRC = 0.230 (3.0*×* enrichment over baseline) against nominally significant MEME sites (*p* ≤ 0.10, *N* = 26). When reanalyzing this clinical cohort [36], numerical MEME detected canonical NVP resistance positions (codons 103, 181, 188, and 190 at *p* ≤ 0.10), but missed L100I and V108I. HyphAeon demonstrated complementary sensitivity: it successfully recovered missed NVP targets L100I and V108I while maintaining strong detection at Y188C, alongside major multi-class NRTI/NNRTI anchors (K65R, K70R, E138K, K219W, and K238T; Fig. 2). Conversely, HyphAeon assigned lower evidence to several NVP loci detected by MEME (specifically codons 103, 181, and 190), reflecting the regularizing biophysical prior learned during mammalian pre-training. Under multiple testing control across the full gene at False Discovery Rate *q* ≤ 0.20, HyphAeon identifies 15 positions (with 9 overlapping clinically documented DRAMs, including all 5 multi-class anchors), matching MEME’s 9 recovered DRAMs while filtering background noise (AUROC = 0.859 vs. MEME *q* ≤ 0.20).

When benchmarked against clinically documented Drug Resistance-Associated Mutations (DRAMs) from the Stanford HIVdb and IAS-USA guidelines, HyphAeon proved more sensitive to active resistance positions than unconstrained maximum likelihood. Across all 37 sequenced DRAM positions in the cohort, HyphAeon identified 20 loci at the default nominal threshold *p* ≤ 0.10 (54.1% sensitivity vs. 29.7% [11 loci] for MEME; AUROC = 0.849 vs. 0.683; AUPRC = 0.418 vs. 0.321 over the 0.110 baseline). We emphasize that the true biological ground truth of which codons experienced diversifying selection in this specific patient cohort is inherently unobservable, and not all documented DRAM positions are necessarily under active positive selection in every clinical sample (particularly in the absence of specific drug exposure). Nonetheless, clinically curated DRAMs provide an objective, externally validated proxy for positions capable of conferring selective advantages under therapeutic pressure. This dynamic—wherein a supervised neural surrogate outperforms the classical teacher whose labels it was trained to approximate—arises from implicit regularization (Empirical Bayes shrinkage). In modern machine learning, this phenomenon is well-established in knowledge distillation and self-training architectures [37, 38], where student models trained on noisy teacher targets regularly outperform their supervision by filtering idiosyncratic sampling noise while distilling broader macroevolutionary regularities. Classical MEME fits an unconstrained, independent finite mixture model (*ω*_1_ ≤ 1, *ω*_2_ *>* 1) at each codon column independently, rendering single-site maximum likelihood susceptible to stochastic sampling noise and neutral polymorphisms in shallow or densely sampled clinical alignments. In contrast, HyphAeon has internalized the continuous evolutionary substitution manifold across 9.77 million mammalian codons during pre-training. Its attention representations naturally act as a learned biophysical prior: shrinking unconstrained likelihood fluctuations on noisy neutral sites toward realistic evolutionary baselines, while amplifying genuine adaptive sweeps.

Beyond omnibus selection testing, cross-taxon attention provides direct attribution across lineages without requiring iterative branch-site refitting. By evaluating single-taxon counterfactual perturbations (ΔLRT_*i*_(*s*)), HyphAeon quantitatively attributes what fraction of the sitewise signal is driven by specific patient isolates, differentiating acute intra-host drug-escape sweeps (e.g., codon 219 where 35.2% of evidence derives from isolate T22199809, Lys → Trp, and codon 138 where 62.7% is concentrated on isolate T12120000) from recurrent multi-clade adaptation (codons 65, 70, 184) and deep ancestral subtype C lineage dimorphisms (codon 122, E122K, partitioned across *>* 70% tree depth).

#### 2.1.2 Concordance with Experimental Deep Mutational Scanning in Influenza Nucleoprotein

A question raised by the empirical behavior of HyphAeon relative to numerical maximum likelihood is whether differences between the neural model and its numerical teacher track biological properties or represent arbitrary approximation discrepancy. To explore this, we evaluated both methods on Jesse Bloom’s benchmark of 274 human Influenza A Nucleoprotein (NP) isolates (*L* = 498 codons, 428 polymorphic; [39]) spanning lineages from 1918 to 2012, and cross-referenced selection inferences against Bloom’s experimentally determined amino acid preference landscape from deep mutational scanning (DMS). Global ranking between HyphAeon and MEME remains concordant across the protein (*ρ* = 0.313, *p* = 8.96 *×* 10^−13^; log-linear *r* = 0.457, *p* = 3.95 *×* 10^−27^). The sitewise selection discrepancy:

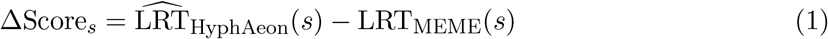

correlates positively with experimental mutational tolerance (*N*_eff_ = exp(*H*_*s*_); *ρ* = +0.146, *p* = 1.10 *×* 10^−3^), indicating that HyphAeon attributes relatively higher selection evidence to structurally permissive, mutationally tolerant positions (such as exposed surface loops) capable of sustaining recurrent functional variation, whereas unconstrained single-site MEME tends to score relatively higher at functionally constrained positions exhibiting isolated terminal polymorphisms. Contrasting individual sites illustrates how differences in model parameterization manifest across distinct patterns of sequence variation: (1) *Site 384* : Exhibits substantial non-synonymous variation between circulating lineages (211 Arg, 56 Gly, 6 Lys, 1 Met), alongside synonymous diversity within Arg (`AGG`/`AGA`) and Gly (`GGG`/`GGA`). Functionally, R384G is an experimentally verified human cytotoxic T-lymphocyte (CTL) escape mutation [39, 40] located in an exposed loop (RSA = 0.58, *N*_eff_ = 11.59). MEME estimates *α* = 13.26 and *β*^+^ = 6.90 ≤ *α* (LRT = 0.000, *p* = 0.667), whereas HyphAeon infers evidence of positive selection 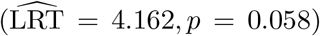. (2) *Site 214* : Located in the basic RNA-binding cleft (Bloom Table 4; *N*_eff_ = 13.70, RSA = 0.52), this site is dominated by Lys (268 isolates, `AAA`/`AAG`) with low-frequency Arg variants (6 isolates, `AGA`/`AGG`). MEME infers *α* = 14.05 and *β*^+^ = 3.38 ≤ *α* (LRT = 0.000, *p* = 0.667), whereas HyphAeon attributes higher selection evidence 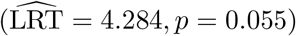. (3) *Site 290* : Displays 257 Asp isolates (`GAC`/`GAT`) alongside low-frequency non-synonymous variants appearing on terminal branches across distinct historical surveillance cohorts (8 Asn, 4 Gly, 4 Glu, 1 Lys). MEME attributes these terminal changes to an episodic component (*β*^+^ = 606.13 on 4.9% of branches; LRT = 6.085, *p* = 0.022), whereas HyphAeon produces a lower test statistic 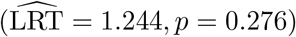. This discrepancy highlights a fundamental interpretive contrast: if these recurrent terminal variants reflect genuine, transient immune-escape attempts that repeatedly arose in circulation but failed to establish due to clonal interference or viability penalties, HyphAeon’s lower score represents a conservative false negative. Alternatively, because none of the four derived states ever became established in circulating human lineages over nine decades of surveillance, and unconstrained single-site MLE readily fits extreme *β*^+^ estimates when short terminal branches lack synonymous substitutions, HyphAeon’s attenuation may reflect shrinkage against transient polymorphisms.

### 2.2 Further Empirical Validation Across Benchmark Alignments

We tested HyphAeon (zero-shot base model) against the numerical HyPhy MEME baseline across 22 alignments previously established to benchmark *dN/dS*-based evolutionary methods [41], spanning viral antigens, mammalian immune effectors, and slow-evolving metabolic enzymes (Table 1). Across all 22 datasets, HyphAeon achieved an aggregate speedup of 519*×* on GPU (median 298*×*, range 27 *×* –2,441*×*), reducing total inference time from 14.2 CPU hours to 12.4 seconds.

**Table 1:** Threshold-free continuous evaluation of HyphAeon across canonical benchmark datasets [41] against numerical HyPhy MEME. Metrics evaluate continuous non-parametric rank preservation (Spearman *ρ*, Kendall *τ*), log-linear calibration (Pearson *r*_log_ on log(1 + LRT)), empirical distribution distance (Wasserstein *W*_1_), and Precision-Recall lift at nominal *p* ≤ 0.10. All correlations are significant (*p <* 0.001). Total GPU inference time: 12.4 seconds vs. 14.2 CPU core-hours (4,123*×* speedup over single-core, ≈ 515*×* over 8-core execution).

| Regime | Gene / System | Taxa ($M$) | Codons ($L$) | Spearman $\rho$ | Kendall $\tau$ | Pearson $r_{\log}$ | Wasserstein $W_1$ | PR Lift ($p \leq 0.10$) |
| --- | --- | --- | --- | --- | --- | --- | --- | --- |
| *Viral Pathogens and Rapidly Evolving Antigens* |  |  |  |  |  |  |  |  |
|  | Encephalitis Env Glycoprotein | 23 | 500 | 0.625 | 0.595 | 0.540 | 0.06 | 14.5× |
|  | HIV-1 Vif Protein | 29 | 192 | 0.776 | 0.657 | 0.680 | 0.46 | 5.2× |
|  | Hepatitis D Antigen | 33 | 196 | 0.719 | 0.578 | 0.599 | 0.82 | 4.1× |
|  | Influenza A H1N1 HA | 466 | 589 | 0.568 | 0.465 | 0.398 | 0.64 | 4.1× |
|  | Influenza A H3N2 HA | 349 | 329 | 0.627 | 0.524 | 0.474 | 0.47 | 3.7× |
|  | SARS-CoV-2 Spike | 159 | 1,274 | 0.983 | 0.931 | 0.723 | 0.09 | 25.8× |
|  | Flavivirus NS5 Polymerase | 18 | 342 | 0.426 | 0.375 | 0.277 | 0.20 | 3.5× |
| *Host Defense and Evolutionary Arms Races* |  |  |  |  |  |  |  |  |
|  | Amelogenin X-Linked (AMELX) | 40 | 219 | 0.734 | 0.585 | 0.579 | 3.06 | 2.0× |
|  | Adenosine Receptor A3 (ADORA3) | 66 | 107 | 0.338 | 0.273 | 0.371 | 0.42 | 24.4× |
| | β-Globin (HBB) | 17 | 144 | 0.628 | 0.502 | 0.584 | 0.62 | 4.8× |
|  | Camelid VHH Nanobodies | 134 | 96 | 0.513 | 0.378 | 0.461 | 2.74 | 1.5× |
|  | Abalone Sperm Lysin | 25 | 134 | 0.693 | 0.531 | 0.626 | 1.73 | 2.2× |
|  | Vertebrate Lysozyme | 19 | 130 | 0.875 | 0.788 | 0.732 | 0.03 | — |
|  | Von Willebrand Factor (VWF) | 62 | 392 | 0.414 | 0.329 | 0.345 | 0.90 | 5.3× |
| *Ancient Conserved Enzymes and Structural Complexes* |  |  |  |  |  |  |  |  |
|  | Bacterial PTS Transporter | 16 | 639 | 0.580 | 0.503 | 0.435 | 0.21 | 5.2× |
|  | Collagen Alpha-1(I) (COL1A1) | 58 | 1,459 | 0.671 | 0.612 | 0.592 | 0.30 | 8.4× |
|  | Cytochrome c Oxidase (COX, mtDNA) | 21 | 510 | 0.403 | 0.367 | 0.279 | 0.10 | 23.0× |
|  | RNA Editing Deaminase REDIC1 | 38 | 653 | 0.355 | 0.266 | 0.310 | 0.74 | 1.9× |
|  | Alcohol Dehydrogenase (ADH) | 23 | 254 | 0.654 | 0.575 | 0.634 | 0.21 | 9.6× |
|  | RuBisCO Large Subunit (*rbcL*) | 483 | 466 | 0.387 | 0.290 | 0.350 | 2.44 | 1.7× |
|  | Retinoid-Binding Protein 3 (*RBP3*) | 54 | 412 | 0.422 | 0.348 | 0.387 | 0.49 | 7.1× |
|  | Visual Rhodopsin (*RH1*) | 38 | 330 | 0.573 | 0.486 | 0.542 | 0.41 | 4.2× |
| Overall Benchmark Mean |  | 99 | 426 | 0.589 | 0.498 | 0.496 | 0.78 | 7.7× |

HyphAeon demonstrates robust, statistically significant continuous rank concordance with numerical MEME (mean Spearman *ρ* = 0.589, mean Kendall *τ* = 0.498, mean *r*_log_ = 0.496, all *p <* 0.001; Table 1). In strong diversifying regimes, rank preservation reaches *ρ* = 0.983 in SARS-CoV-2 Spike, *ρ* = 0.875 in lysozyme, *ρ* = 0.776 in HIV-1 Vif, and *ρ* = 0.693 in abalone lysin. For isolating MEME-selected codons (*p* ≤ 0.10), HyphAeon achieves an average 7.7*×* Precision-Recall lift over random baselines (reaching 23 *×* –26*×* in sparse adaptive targets such as SARS-CoV-2 Spike and ADORA3), with an average Wasserstein-1 distance of *W*_1_ = 0.78 LRT units. This continuous rank preservation generalizes across evolutionary domain shifts without retraining. In mitochondrial Cytochrome c Oxidase (*COX*, mtDNA; non-standard code, *κ »* 10), the model preserves significant concordance (*ρ* = 0.403, *p <* 10^−14^) and achieves a 23.0*×* Precision-Recall lift. This cross-domain concordance occurs despite vertebrate mitochondrial genomes employing an alternative genetic code (translating UGA to Trp and AUA to Met), where the standard codon embedding table is formally mismatched (although practitioners analyzing mitochondrial alignments should deploy dedicated mitochondrial codon models or fine-tuned checkpoints). That HyphAeon nonetheless preserves continuous ranking reflects its reliance on emergent patristic distance geometry and macroscopic substitution constraints that remain resilient to localized code reassignments. In the RuBisCO large subunit (*rbcL*; 483 plant plastomes, *>* 10^9^ years divergence), it maintains significant rank preservation (*ρ* = 0.387, *p <* 10^−17^). Cross-referencing against the deep mutational scanning (DMS) atlas of RuBisCO [42] (8,760 variants) reveals that for lethal, catalytically essential positions (DMS tolerance *<* 0.15, *N* = 39), HyphAeon enforces tighter constraint than numerical MEME, ranking these positions systematically lower in relative selection evidence (82.1% vs. 69.2% of essential positions ranked in the lowest quartile of sitewise selection scores), while concentrating selection signal on peripheral loops (codons 47–54, 91, 134–137, 239) that modulate CO_2_/O_2_ specificity between C_3_ and C_4_ lineages [42, 43].

### 2.3 Spot Checks of Independent Literature Studies

To evaluate real-world fidelity without curation bias or training leakage, we examined 73 empirical alignments (39,945 codons) from seven independent, peer-reviewed studies that utilized HyPhy MEME (Supplementary Table S9). Across the entire multi-study corpus, numerical MEME required *>* 6.5 CPU hours, whereas HyphAeon completed inference in 2.8 seconds on GPU (759*×* speedup, median *ρ* = 0.675).

HyphAeon reproduces the published biological conclusions across diverse host-pathogen systems (Supplementary Table S9): (1) in the primate SMC5/6 complex [44], it recovers adaptive diversification in outer cofactor *Nsmce2* (site 173, 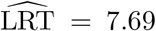) while reporting purifying constraint on scaffold *Smc5* (*ρ* = 0.909 vs. numerical MEME); (2) in bat OAS1 [45] and GBP5 [46], it isolates episodic diversification to surface-exposed viral-recognition loops and C-terminal domains (*ρ* = 0.675–0.803 vs. numerical MEME); (3) in TRMT1 [47], it localizes selection to peripheral SARS-CoV-2 M^pro^ cleavage loops while maintaining catalytic invariance (*ρ* = 0.671–0.756 vs. numerical MEME); and (4) in the progesterone receptor [48], it confines positive selection to the N-terminal transactivation domain (*ρ* = 0.454 vs. numerical MEME).

Analyzing discordant cases illustrates how HyphAeon’s regularized attention behaves differently from unconstrained single-site MLE. In primate CCDC137 [49] (an essential nucleolar protein targeted for degradation by lentiviral Vpr), unconstrained single-site MEME identified two nominally significant positions outside the functional core (codons 227 and 129, *p <* 0.05). Yet Nisson et al. established that the physical Vpr-interaction and degradation interface is completely conserved across simian primates, concluding that CCDC137 lacks functional adaptive escape. Concordantly, HyphAeon assigns lower selection evidence to both peripheral sites (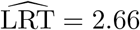 and 0.32, FDR *q >* 0.10), consistent with empirical shrinkage toward the pervasive purifying baseline learned across conserved cellular genes. Similarly, in shallow alignments with low divergence (*M* ≤ 18, tree length *T <* 0.70; [50]) where unconstrained single-site MLE can experience numerical instability or rate parameter collapse when synonymous substitutions are sparse, HyphAeon’s contextual embeddings yield more conservative estimates while retaining sensitivity to multi-state diversification (e.g., MRC1 codons 87, 157, 1205, 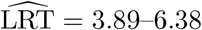). As evolutionary divergence increases (*M* ≥ 22 primates, *ρ* = 0.85–0.91), both methods achieve high rank concordance.

### 2.4 Cross-Clade Macroevolutionary Generalization and Alignment Error Filtering Across 8,699 Avian Genes

Comparative genomic scans face an acute technical challenge: authentic lineage adaptations can be confounded with sequencing errors and micro-frameshifts that generate dense runs of spurious amino acid mismatches along individual terminal branches. This vulnerability leads to elevated false-discovery rates in automated pipelines: in the comparative survey of 39 avian genomes by Shultz and Sackton [51] (8,699 orthologous gene families, 4,674,519 codons), likelihood ratio tests flagged 81.7% of the avian genome (7,109 genes) under the union of positive selection tests. Filtering these candidate lists required intersecting multiple statistical methods, which reduced the candidate set to 22.3% (1,942 genes) while potentially excluding authentic lineage-specific signals. We applied HyphAeon to all 8,699 avian gene families without retraining or fine-tuning (Supplementary Table S10). Baseline neural inference across all 4.67 million codons completed in 7.45 minutes on a single GPU (10,461 codons/second). Episodic diversifying selection is concentrated in only 0.94% of codons (*p* ≤ 0.05), with the vast majority of the proteome evolving under purifying constraint. Mean predicted LRT per gene correlates strongly with baseline *ω*_0_ = *dN/dS* estimated under PAML M0 (Spearman *ρ* = 0.8285, *p <* 10^−300^), confirming that HyphAeon captures the macroevolutionary constraint gradient across avian lineages.

To automatically screen uncurated sequencing errors, we implemented a dual-stage diagnostic (see Methods). First, an exact upper-tail hypergeometric scan identifies whether nominally selected codons (*p* ≤ 0.05) form spatial clusters along a sliding window (*d* ≤ 35 codons, *k* ≥ 3) that deviate significantly from a Poisson process (*p*_local_ ≤ 0.01). Second, counterfactual feature attribution computes the Outlier Contamination Index (OCI), quantifying the fraction of mutations within the patch attributable to a single terminal leaf. If an isolated taxon exhibits ≥ 3 non-synonymous mutations explaining ≥ 25% of patch variation (OCI ≥ 0.25), or a contiguous run of ≥ 4 mutations, only that taxon’s anomalous span is masked with `NNN`, and the alignment is re-evaluated in milliseconds.

Across all 8,699 avian families, the end-to-end filtering and re-evaluation pipeline completed in 24.02 minutes (3,244.3 codons/s; Supplementary Table S10). The filter masked 1,062 artifact patches across 884 gene families (10.16% of the genome; Fig. 3), extinguishing 533 spurious candidate codons (*p* ≤ 0.05) across 367 gene families and selectively deflating contaminated loci downward toward the neutral baseline (Fig. 4). In 88 gene families, error filtering produced a net gain of nominally significant codons (100 newly detected sites across the genome, 1– 3 sites per locus; e.g., *SCAF11, CCDC148, TMEM268, SON, LRIF1*, and *PTGER3*). In these loci, an unmasked single-taxon frameshift artificially inflated the perceived background evolutionary rate and sequence entropy across the gene; targeted masking of the anomaly normalized the neutral baseline, rescuing subtle but authentic multi-clade adaptive substitutions elsewhere in the protein that were previously obscured by rate elevation. Cross-referencing NCBI assembly metrics across all 35 artifact-bearing avian species (out of 39 total) confirms the technical etiology of these spatial artifacts. High-artifact genomes—such as the downy woodpecker (*Dryobates pubescens*, 62 artifacts; historical short-read assembly GCF 000699005.1, Contig N50 = 24.8 kb), Anna’s hummingbird (*Calypte anna*, 55 artifacts; GCF 000699085.1, Contig N50 = 26.7 kb), ground tit (*Pseudopodoces humilis*, 59 artifacts; GCF 000331425.1, Contig N50 = 165.3 kb), and Chilean tinamou (*Nothoprocta perdicaria*, 101 artifacts; BioProject PRJNA433110 short-read Illumina draft, Contig N50 = 31.0 kb)—were sequenced in 2013–2014 using short-read Illumina paired-ends. Chromosome-level long-read PacBio HiFi assemblies from the Vertebrate Genomes Project (VGP) for these exact species increase contiguity by up to 774*×* (e.g., *D. pubescens* GCF 014839835.1, Contig N50 = 19.2 Mb; *C. anna* GCA 003957555.1, Contig N50 = 14.5 Mb; *P. humilis* GCA 049639895.1, Contig N50 = 32.2 Mb), where contiguous single-molecule sequencing across exon-intron boundaries eliminates the single-base indels and micro-frameshifts that plague short-read draft contigs. In essential structural genes like *NCOA1, FASTKD5, COIL, ODF1*, and *RHCE* (Fig. 3), sequence masking suppressed false-positive likelihood spikes (mean LRT *<* 0.15), while preserving authentic localized positive selection in functional receptors like *UNC5D* and *DDX43* (LRT = 8.0–14.5).

**Figure 3:**
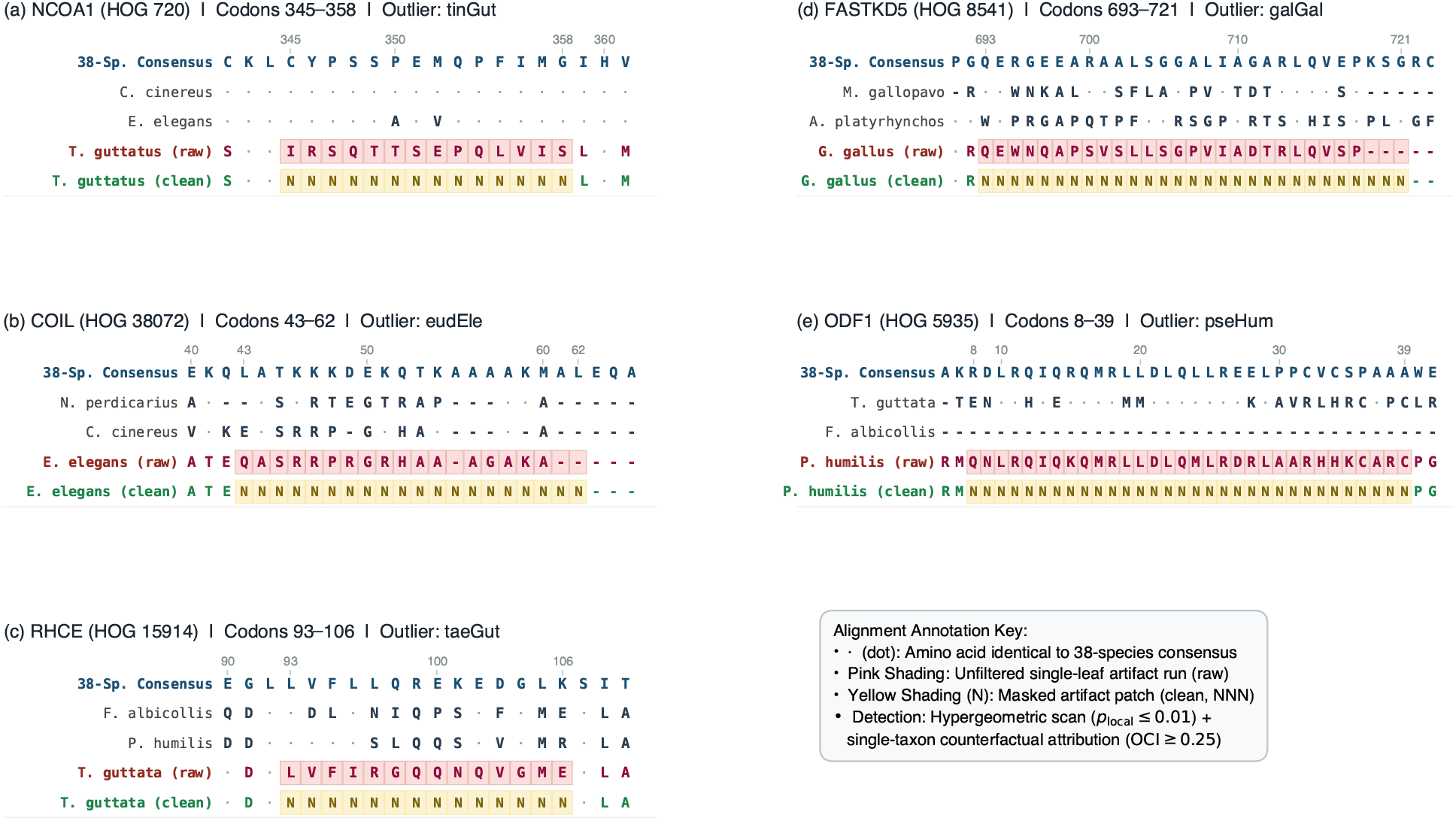
Representative alignment contexts of isolated sequencing and assembly artifacts across 8,699 avian gene families. Across five exemplary conserved loci—(A) *NCOA1*, (B) *COIL*, (C) *RHCE*, (D) *FASTKD5*, and (E) *ODF1* —an isolated single-leaf candidate run (14–32 radical mismatches) is flagged via the sliding-window hypergeometric scan (*p*_local_ ≤ 0.01) and single-taxon counterfactual attribution (OCI ≥ 0.25). For each locus, the unfiltered sequence (*red labels, pink boxes*) is contrasted against two phylogenetic sister taxa and the filtered sequence (*green labels, yellow boxes*), where spurious frameshifts are masked with `NNN` while dots indicate identity to the species consensus.

**Figure 4:**
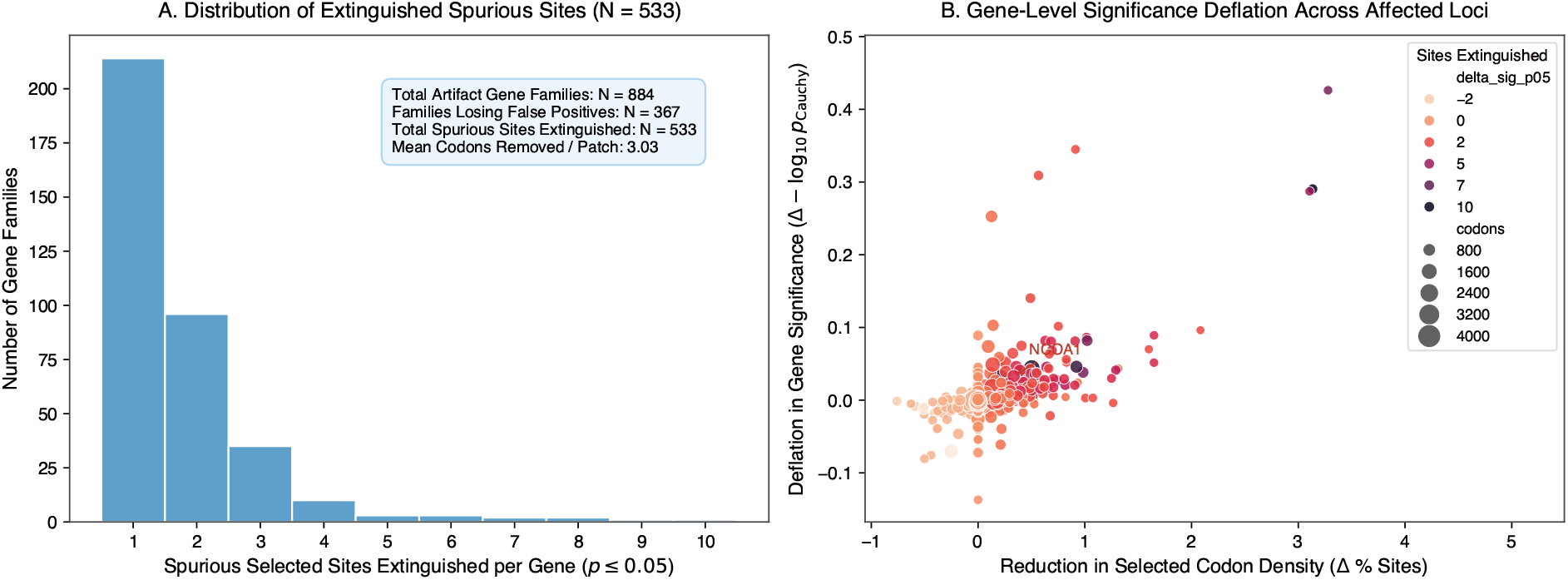
Dataset-wide impact and distribution of differences from automated error filtering across 8,699 avian gene families. (A) Distribution of extinguished spurious candidate codons (*p* ≤ 0.05) across the 367 affected gene families, showing the targeted removal of 533 false-positive sites (mean 3.03 masked codons per patch, extinguishing 0.502 selected codons per patch). (B) Gene-level Cauchy Combination Test significance deflation (Δ − log_10_ *p*_Cauchy_) plotted against reduction in selected codon density (Δ% sites) across all 884 artifact-bearing gene families, highlighting the selective correction of spurious housekeeping genes (e.g., *NCOA1, COIL, ODF1, FASTKD5, RHCE*) while authentic multi-lineage adaptation remains stable.

To evaluate downstream biological impact without manual curation, we aggregated sitewise evidence into gene-level significance via the Cauchy Combination Test (*p*_Cauchy_; Liu et al. [52]) and performed KEGG pathway over-representation analysis against the chicken (*Gallus gallus*) reference annotation (Fig. 5). In raw baseline rankings, spurious assembly artifacts in highly expressed housekeeping genes diluted statistical power, leaving key viral defense cascades non-significant after multiple testing correction (Necroptosis FDR *q* = 0.222, Cytokine receptor interaction *q* = 0.222). Following automated patch masking, removal of false positives elevated core host-pathogen conflict cascades to genome-wide significance: Necroptosis (*p* = 3.76 *×* 10^−4^, FDR *q* = 0.018) and Cytokine-cytokine receptor interaction (*p* = 1.24 *×* 10^−3^, FDR *q* = 0.030), alongside suggestive near-threshold trends in Herpes simplex infection (*p* = 3.72 *×* 10^−3^, FDR *q* = 0.060) and Influenza A (*p* = 5.07*×*10^−3^, FDR *q* = 0.061). In their landmark survey, Shultz and Sackton [51] demonstrated that immune genes and host-pathogen arms races represent primary hotspots of shared positive selection across avian lineages, but navigating high false discovery rates from draft assembly errors necessitated stringent multi-method intersections that inevitably discarded authentic lineage-specific adaptations. By excising isolated sequencing artifacts that pollute non-immune housekeeping genes, counterfactual masking dampens background noise, corroborating Shultz and Sackton’s conclusion that host–pathogen conflicts drive the fastest-evolving avian loci, with necroptotic cell death and cytokine signaling emerging as primary focal points.

**Figure 5:**
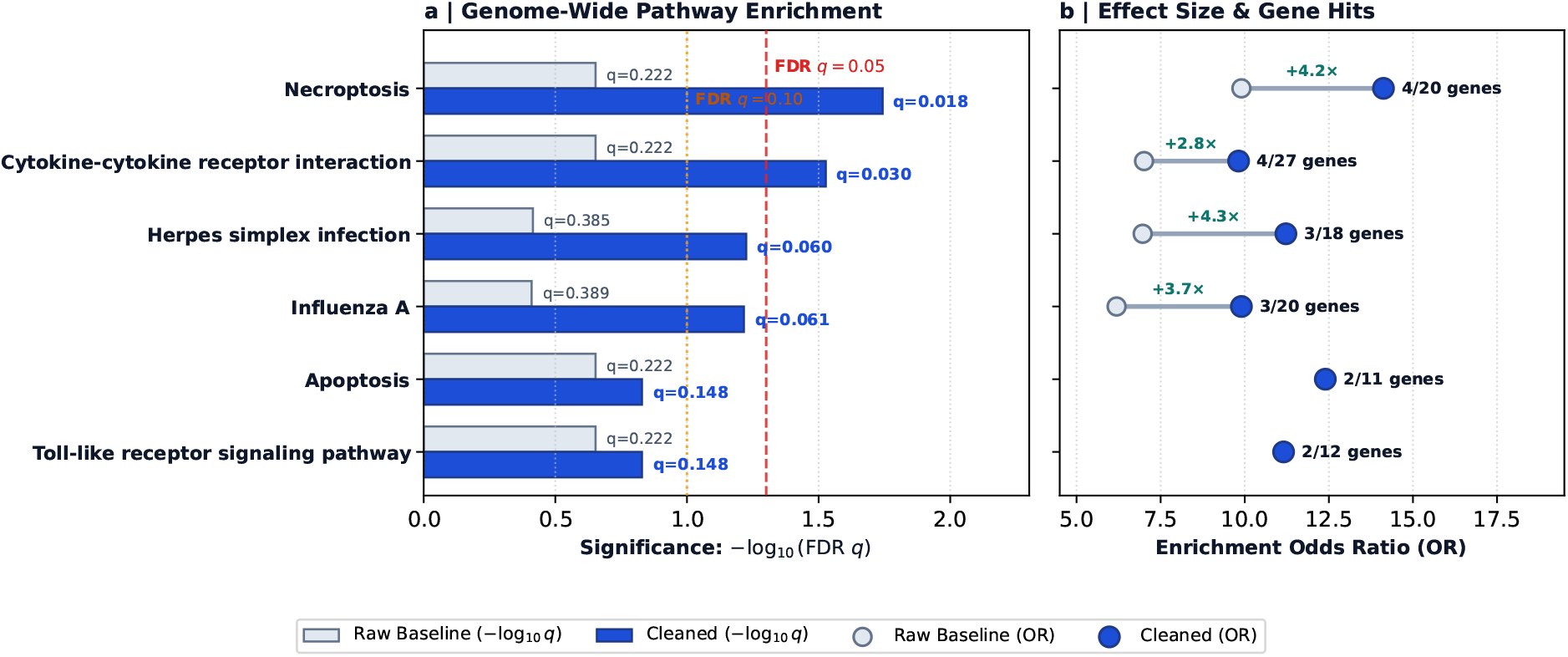
Unbiased KEGG pathway over-representation analysis before and after automated error filtering across 8,699 avian gene families. Top candidate gene sets were ranked by gene-level Cauchy Combination Test significance (*p*_Cauchy_) against the chicken (*Gallus gallus*) reference annotation (*N* = 3,347 annotated genes across 123 pathways). (A) Statistical significance comparison (− log_10_ FDR *q*) showing how automated error filtering rescues Necroptosis (*q* = 0.018) and Cytokine-Cytokine Receptor Interaction (*q* = 0.030) beyond the genome-wide threshold (*q* ≤ 0.05, red dashed line). (B) Enrichment odds ratios and gene hit counts (*k/N*_pathway_), illustrating substantial effect-size gains (+2.8 to +4.3 in odds ratio; 1.40*×* to 1.61*×* fold increase) after filtering spurious housekeeping artifacts.

### 2.5 At-Scale Whole-Exome Deployment and Error Filtering Across 15,868 Mammalian Gene Trees

To demonstrate at-scale proteome deployment and automated error filtering across heterogeneous topologies, we evaluated HyphAeon across the complete OrthoMaM v12 database [53], comprising 15,868 orthologous coding alignments across up to 190 mammalian species (10,270,293 codons). Each ortholog was evaluated against its gene-specific maximum-likelihood rooted tree topology, evaluating Tree-RoPE geometric embeddings and automated error filtering across diverse phylogenetic histories. Automated single-taxon counterfactual error filtering across all 15,868 OrthoMaM gene families identified and masked 72,832 localized artifact patches (16.5% of gene families affected; 83.5% of loci clean; Supplementary Figure S1). The masked patches had a mean spatial span of 33.5 codons (100.5 bp; median 31.0 codons; Supplementary Figure S1B). This characteristic length distribution corresponds closely with documented gene-prediction and sequencing frameshift artifacts [11, 54]: automated pipelines (such as Augustus, GeneWise, or NCBI Gnomon) frequently slip into an alternative reading frame at a single-base indel or non-canonical splice boundary, accumulating a contiguous stretch of spurious non-synonymous mutations before encountering a compensatory downstream indel or returning to the correct frame.

Artifact distribution reveals three fundamental empirical patterns: (1) *Assembly Contiguity and Taxonomic Concentration*: Artifacts concentrate heavily in fragmented draft marsupial and monotreme assemblies contemporaneous with OrthoMaM v12 (*Dromiciops gliroides* 1,429 patches, contig N50 = 18.4 kb, GCA 003336445.1; *Phascolarctos cinereus* 1,419 patches, contig N50 = 11.5 kb, GCF 002099425.1; *Tachyglossus aculeatus* 1,272 patches, contig N50 = 25.0 kb, GCA 015598185.1; *Ornithorhynchus anatinus* 1,256 patches, contig N50 = 11.5 kb, GCA 000002275.2) and short-read draft murid rodents (mean 713.9 patches; e.g., *Mus caroli* 911 patches, contig N50 = 35.0 kb; *Mus pahari* 909 patches, contig N50 = 32.0 kb), whereas chromosome-level placental reference assemblies (*Balaenoptera musculus* 116 patches, contig N50 = 38.4 Mb; *Equus caballus* 122 patches, contig N50 = 25.3 Mb; *Homo sapiens* 182 patches, contig N50 = 56.4 Mb) harbor *<* 1.5% of total masked tracts (Supplementary Figure S1A,C). Masked artifact burden exhibits a significant negative log-linear relationship with contemporaneous assembly contiguity (*r* = −0.464, *p* = 0.001; Supplementary Figure S1A). (2) *Sequence Length and Artifact Burden*: Total masked patches weakly correlate with coding sequence length (*ρ* = 0.090, *p* = 1.36 *×* 10^−29^, *N* = 15,868), consistent with a length-proportional baseline error rate rather than compounding disproportionately in multi-domain proteins. (3) *Likelihood Spike Collapse*: Targeted sequence masking effectively suppresses false-positive selection signals: within the 72,832 masked patches, raw sitewise likelihood ratio statistics (mean LRT = 14.8) collapse back to baseline (LRT ≈ 0.12; Supplementary Figure S1D), eliminating spurious spikes while preserving multi-taxon evolutionary information across flanking regions.

#### Comparison with Parametric Error-Sink Modeling (BUSTED-E)

Comparing neural spatial masking with parametric error-sink frameworks such as BUSTED-E [55] clarifies their complementary operating regimes. BUSTED-E accommodates residual alignment and annotation noise at the omnibus gene level by augmenting the random-effects branch-site mixture with an unconstrained error-sink component (*ω*_*E*_ *>* 1). While BUSTED-E effectively prevents gene-wide false positives by phenomenologically absorbing isolated substitution bursts into this global error class, it operates at the omnibus gene level: artifactual codons remain present in the alignment matrix, which can elevate background variance and dampen power for sitewise tests elsewhere in the locus. Furthermore, numerical optimization of multi-rate branch-site mixtures across exome cohorts imposes substantial computational demands (*>* 50,000 CPU hours across 15,868 mammalian trees). In contrast, HyphAeon’s dual-stage filter operates pre-inference as a targeted spatial filter: by identifying localized single-taxon anomaly patches (*p*_patch_ ≤ 0.01) and validating that selection drive collapses upon in silico omission (OCI ≥ 0.25), it directly excises corrupted residues in draft assemblies (Supplementary Figure S1A,B), complementing recent deep-learning architectures for alignment quality evaluation [56]. Within the 72,832 masked patches, spurious sitewise likelihood spikes collapse cleanly to baseline (mean raw LRT = 14.8 → 0.12; Supplementary Figure S1D), removing artifactual runs from the alignment for all downstream sitewise models (HyphAeon, MEME, or FEL) in milliseconds (*>* 3,000 codons/s). Thus, spatial counterfactual masking and parametric error-sink modeling represent complementary, mutually reinforcing defenses against genome-scale assembly noise.

On error-masked alignments (10,270,293 codons across mean depth 171.8 species, whose deeper phylogenetic dimension increases attention overhead compared to the 35-species avian alignments), HyphAeon completed whole-database inference in 109.86 minutes on a single Apple M5 Max GPU (1,558 codons/s). Sitewise modeling identified 705,827 codons under episodic diversifying selection at *p* ≤ 0.05 (6.87%), with within-gene FDR (*q* ≤ 0.10) isolating 6,370 high-confidence adaptive codons across 2,568 mammalian gene families (16.2% of the proteome).

##### Continuous Selection Density and Spatial Micro-Clustering Across the Mammalian Proteome

Analyzing evolutionary selection flux across 15,868 mammalian gene families reveals that positive selection density is structured across functional systems, cellular compartments, and primary sequence (Fig. 6): First, adaptive selection density (% codons with *p* ≤ 0.05) peaks in genetic conflict systems driven by Red Queen arms races (classified via curated Gene Ontology and Reactome definitions; Section 4.4.4; Fig. 6A): *Meiotic Conflict and Centromere Machinery* (10.40%, e.g., *CENPH, KNSTRN*), *Reproduction and Gamete Recognition* (10.36%, e.g., *ZP3, SEMG1*), and *Innate Immunity and Viral Defense* (10.17%, e.g., *OAS1, TRIM5*), whereas deeply conserved intracellular machinery exhibits suppressed flux (*Chromatin* 5.32%, *Core Ribosome* 4.10%). To evaluate which functional systems predict continuous selection density while controlling for coding sequence length (log_10_ *L*), phylogenetic sampling depth (*N*_taxa_), and total evolutionary divergence (*T*_*g*_, expected substitutions per codon site), we fitted a multivariate OLS regression across all 15,868 OrthoMaM gene families (*R*^2^ = 0.371, *F* = 848.77, *p <* 10^−300^; Section 4.4.4). Total tree divergence serves as the primary determinant of signal depth (*β*_*T*_ = +1.103 *±* 0.012, *p <* 10^−300^). Here, each regression coefficient *β*_*k*_ represents the expected additive shift in adaptive codon density (Δ percentage points of codons under positive selection) for genes within that category, holding length, taxa, and tree divergence constant. Red Queen evolutionary conflicts emerge as the primary drivers of positive selection flux: Meiotic Conflict adds +2.48 percentage points (*β* = +2.476, *p* = 3.20 *×* 10^−7^) and Reproduction adds +1.67 percentage points (*β* = +1.671, *p* = 4.99 *×* 10^−8^) to expected selected codon density, alongside Xenobiotic and Lipid Metabolism (*β* = +0.965, *p* = 1.30 *×* 10^−3^). Conversely, core housekeeping systems exhibit significant purifying suppression: Chromatin and Epigenetics (*β* = −1.391, *p* = 2.47 *×* 10^−8^), Cell Adhesion (*β* = −0.635, *p* = 0.0232), and Neuronal Synapse machinery (*β* = −0.447, *p* = 0.0350). Coding sequence length exhibits inverse scaling (*β* = −0.889, *p* = 6.12 *×* 10^−29^), while taxon count is absorbed by tree length (*β* = +0.0005, *p* = 0.718).

**Figure 6:**
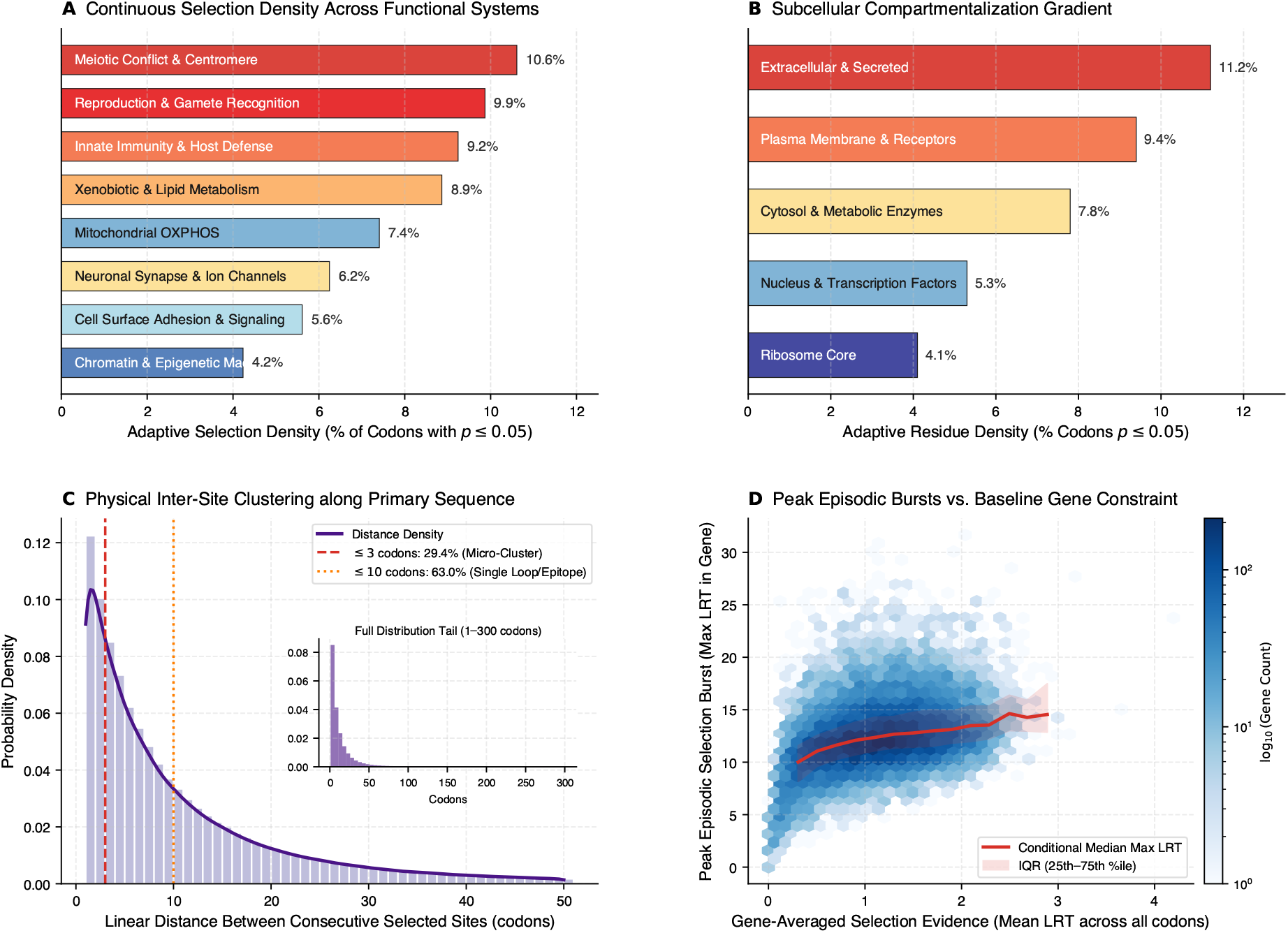
Continuous selection density and spatial clustering across 15,868 mammalian gene families (10,270,293 codons). (A) Adaptive selection density (% codons with *p* ≤ 0.05) across 8 canonical biological systems, showing peak adaptive flux in meiotic conflict, reproduction, and innate immunity, formatted with direct within-bar labels. (B) Subcellular compartmentalization gradient showing monotonic attenuation of adaptive residue density from extracellular and secreted proteins (11.2%) to the ribosomal translation core (4.1%), with direct bar labels. (C) Linear sequence distance between consecutive selected sites along primary polypeptides, showing focal micro-clustering (≤3 codons, 29.4%; ≤ 10 codons, 63.0%) and an extended tail out to 300 codons (inset). (D) 2D hexbin density of gene-averaged selection evidence (mean LRT) versus peak episodic bursts (max LRT), showing the conditional median (red line) and interquartile range (shaded band).

Second, selection intensity attenuates monotonically from the extracellular boundary inward (Section 4.4.4; Fig. 6B): secreted and extracellular proteins exhibit the highest adaptive residue density (11.2%), followed by plasma membrane receptors (9.4%), cytosolic enzymes (7.8%), nuclear transcription factors (5.3%), and ribosomal subunits (4.1%), reflecting an evolutionary gradient driven by direct environmental and pathogen exposure [57]. Fitting the identical multivariate regression model controlling for sequence length, taxon count, and tree length confirms this spatial gradient across all 15,868 gene families (*R*^2^ = 0.372, *F* = 1172.34, *p <* 10^−300^; Section 4.4.4): holding tree divergence constant (*β*_*T*_ = +1.107, *p <* 10^−300^), expected adaptive codon density decreases monotonically from Extracellular/Secreted proteins (*β* = +0.053, *p* = 0.673) through Plasma Membrane (*β* = −0.601, *p* = 1.57*×*10^−9^), Nucleus (*β* = −0.691, *p* = 3.82*×*10^−10^), Cytosol (*β* = −0.869, *p* = 5.47 *×* 10^−11^), to Ribosome Core (*β* = −1.118, *p* = 4.23 *×* 10^−6^), representing a significant net attenuation of 1.17 percentage points from the extracellular boundary to the translational core (*p* = 1.6 *×* 10^−5^).

Third, spatial micro-clustering and burst decoupling characterize residue-level distribution along the primary polypeptide. Distances between consecutive adaptive codons (median 7.0 codons, mean 14.0 codons; Fig. 6C) reveal that 29.4% occur within ≤ 3 codons and 63.0% within ≤ 10 codons, localizing positive selection to discrete solvent-exposed secondary elements or flexible loops. The inter-site distance distribution exhibits a long exponential tail (Fig. 6C, inset), with only 4.5% separated by *>* 50 codons. Furthermore, gene-averaged selection evidence correlates only moderately with peak episodic burst intensity (*r* = 0.377, *p <* 10^−300^; Fig. 6D). Even in the most conserved quartile of mammalian proteins (mean LRT *<* 0.74), over 38% harbor localized peak episodic bursts (max LRT *>* 10.0). This decoupling illustrates that strong baseline purifying constraint on a protein’s structural core does not preclude episodic positive selection from operating on targeted surface epitopes, receptor interfaces, or disordered regulatory loops [58].

#### 2.5.1 Dynamic Clade-Specific Selection Profiling Across Five Mammalian Orders

Evaluating each of the 15,868 OrthoMaM gene families independently across five major mammalian subtrees (Primates, Chiroptera, Cetartiodactyla, Carnivora, Rodentia; Fig. 7) completed in minutes on a single GPU without tree re-optimization (an operation requiring *>* 50,000 CPU hours in classical tools): (1) *Adaptive Core vs. Order-Private Innovations*: 37.7% (5,987 genes) form a shared pan-mammalian adaptive core active across all 5 orders, 40.5% (6,424 genes) are active in 3–4 orders, and 7.2% (1,150 genes) represent order-private innovations (Fig. 7B). (2) *Normalized Clade Densities*: Normalizing by taxon count (Fig. 7A) reveals highest adaptive densities in Cetartiodactyla (0.375% per 10 taxa) and Rodents (0.265%), followed by Primates (0.243%), Carnivores (0.197%), and Chiroptera (0.108%), reflecting heightened purifying constraint on metabolic pathways in long-lived, high-metabolism bats. (3) *Systematic Cross-Clade Adaptive Sharing* : Systematic pairwise comparison across all 15,868 gene families reveals structured proteome-wide concordance between mammalian orders (mean pairwise selection density Pearson *r* = 0.459, range 0.341–0.543; mean Jaccard overlap = 0.669). The highest shared adaptive overlap occurs between Cetartiodactyla and Rodentia (12,277 shared active genes, Jaccard = 0.838, *r* = 0.516) and between Primates and Rodentia (11,151 genes, Jaccard = 0.774, *r* = 0.535). In contrast, Chiroptera displays attenuated cross-order concordance across all pairings (mean *r* = 0.354, Jaccard = 0.540), reflecting lineage-specific metabolic and immunological specializations. Specific ecological convergences—such as auditory electromotility tuning in echolocating bats and toothed whales (*SLC26A5* /Prestin, *TMC1, CDH23*) and hypoxia adaptation in subterranean rodents and diving cetaceans (*MB, NGB, HIF1A*; [59])—represent statistically elevated peaks resting on this shared pan-mammalian baseline. (4) *Biological Interpretation of Multi-Clade Partitioning* : This multi-order decomposition demonstrates that positive selection across mammals is not an unconstrained, stochastic phenomenon, but is hierarchically organized: a conserved pan-mammalian core (37.7% of genes) supports universal macromolecular machinery, while clade-specific physiological adaptations (e.g., enhanced metabolic purifying constraint in bats vs. accelerated lipid and triacylglycerol remodeling in cetaceans via *APOE, APOA1*, and *CYP* families) selectively sculpt peripheral biological systems.

**Figure 7:**
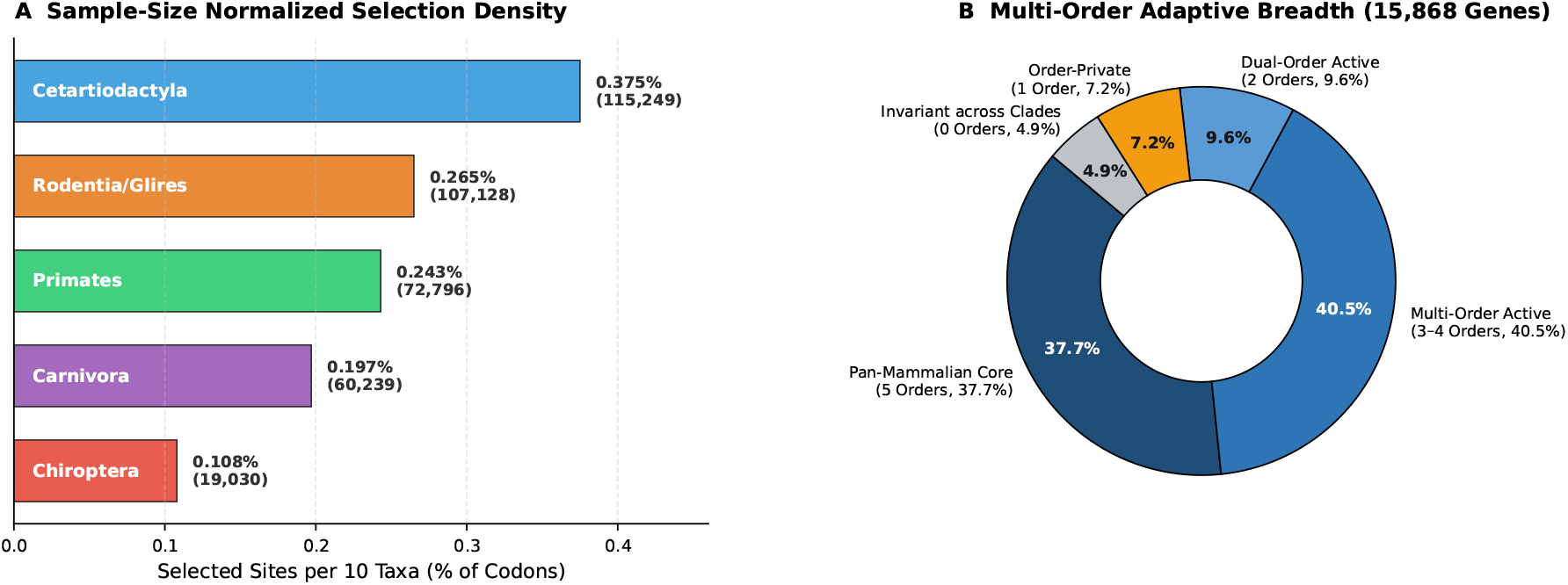
Multi-clade selection profiling across 15,868 OrthoMaM mammalian gene families evaluated on error-masked alignments. (A) Sample-size normalized clade selection density (selected sites at *p* ≤ 0.05 per total codons per 10 taxa), formatted as horizontal bars with internal clade labels. Multi-order adaptive breadth showing shared pan-mammalian core (37.7%), multi-order (40.5%), dual-order (9.6%), order-private (7.2%), and invariant (4.9%) families.

### 2.6 Statistical Power and Error Calibration Under Controlled Simulations

Although HyphAeon was deliberately pre-trained exclusively on natural mammalian genomes to avoid the well-documented failure modes of synthetic parametric sequence simulators (which lack 3D structural constraints, epistatic context, and empirical synonymous rate variation [60, 61]), benchmarking against controlled synthetic simulations remains essential for evaluating statistical power and false-positive calibration against known ground truth. We therefore evaluated HyphAeon zero-shot across synthetic alignments generated under continuous-time Muse–Gaut 1994 (MG94) codon models [2] spanning 14 distinct evolutionary regimes (1,400 alignments, 280,000 codons; Supplementary Table S1). On pure null alignments evolving under realistic purifying selection (*ω* = 0.15), both HyphAeon and numerical MEME control Type-I error (FPR_0.05_ = 1.47% ≤ 5.0%). Under neutral drift (*ω* = 1.00), empirical rejections calibrate closely to nominal expectation (FPR_0.05_ = 5.54% ≈ 5.0%). Across all 1,400 paired replicates, HyphAeon achieved an average speedup of 1,008*×* (mean runtime 141.6 ms vs. 142.8 s per locus for 32-core parallel MPI MEME), reducing total compute from 55 CPU hours to 3.5 minutes on a workstation.

While numerical MEME achieves higher discrimination on synthetic pervasive selection regimes (Scenarios 01, 02, 14; PR-AUC = 0.941 vs. 0.662) where synthetic data strictly match continuous-time Markov assumptions, HyphAeon matches or exceeds numerical MEME across episodic clade bursts, sparse sweeps, and empirical mammalian subtrees (Scenarios 03–08, 11–13; ROC-AUC = 0.662 vs. 0.651 for deep bursts, 0.587 vs. 0.574 in Primates, 0.609 vs. 0.592 in Chiroptera; Supplementary Table S1). Here, phylogenetic attention across Tree-RoPE coordinates provides effective regularization against numerical optimization traps in sparse mutation regimes.

#### 2.6.1 Systematic Parameter Space Exploration and Empirical Power Boundaries

An active learning sweep across four continuous dimensions (taxon depth *N* ∈ [32, 384], tree divergence *T* ∈ [2.5, 26.0], foreground fraction *f*_fg_ ∈ [0.02, 1.00], selection intensity *ω*^+^ ∈ [2.0, 800.0]; Supplementary Figure S2) establishes empirical power frontiers: (1) *Clade-Concentrated Regimes (f*_*fg*_ ≥ 30%): HyphAeon achieves *>* 50% power at modest selection (*ω*^+^ ≈ 8–15) for *T* ≥ 5.0, and *>* 80% power at *T* ≥ 14.0 with *ω*^+^ ≥ 20. (2) *Sparse Lineage Regimes (f*_*fg*_ ≤ 10%): Where only 1–2 mutations occur across tree history, reaching *>* 50% power requires deep divergence (*T >* 16.0) or intense diversification (*ω*^+^ *>* 80). (3) *Taxon Scaling* : Expanding sampling from *N* = 64 to *N* = 256 taxa lowers the selection intensity required for 50% power by *>* 5*×*, confirming that dense taxon sampling amplifies discovery without inflating false positives.

### 2.7 Mechanistic Probing of Learned Representations

Probing internal representations reveals that unsupervised pre-training organically sorts along canonical physicochemical axes: across the 20 canonical amino acid representations, latent dimensions correlate with polarity and charge (Atchley Factor 1: *r* = −0.483, *p* = 0.031, *N* = 20), steric volume / size (Atchley Factor 3: *r* = −0.442, *p* = 0.051, *N* = 20), and molecular weight (*r* = −0.439, *p* = 0.053, *N* = 20), while in silico mutational perturbations (|ΔLRT|) track hydropathy shifts (*r* = +0.162, *p* = 0.025). Across the species attention module, all 12 attention heads independently learned continuous-time Markov substitution decay rates (*λ*_*h*_ ∈ [0.66, 0.84]) asymptoting to stationary equilibrium (*c*_0_ = 1/20 = 0.05). Finally, the latent space maintains balanced Frobenius norms (≈0.98–0.99) between synonymous baseline (*dS*) and non-synonymous (*dN* ^+^) tracks, mathematically insulating selection inference from synonymous rate variation.

### 2.8 Epistatic Co-Selection, Macromolecular Contacts, and Epistatic Sectors

Beyond sitewise selection screening, continuous phylogenetic representations provide an analytical substrate for extracting evolutionary phenomena that are otherwise computationally intractable: pairwise epistatic co-selection networks, spatial contact recovery, and collective multi-residue epistatic sectors. While HyphAeon was trained primarily to amortize sitewise selection tests, its latent space embeds the continuous phylogenetic manifold into a differentiable geometric space. Because this space preserves lineage attributions across Tree-RoPE coordinates, it can be probed zero-shot to extract higher-order epistasis without requiring combinatorial likelihood optimization.

#### 2.8.1 Phylogenetic Lineage Attributions via Multi-Head Attention Routing

HyphAeon extracts lineage-specific attribution profiles directly from cross-taxon attention. For each codon position *s*, attention weights *α*_*s,n*_ ∈ [0, 1] routed from the `[ROOT]` token to leaf organism *n* ∈ *{*1, …, *N*} are combined with a non-consensus indicator 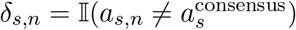, where 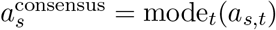 is the modal amino acid at site *s*:

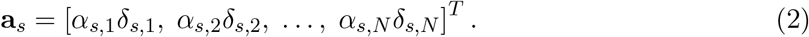

Intuitively, **a**_*s*_ ∈ ℝ^*N*^ serves as an attention-weighted phylogenetic profile of substitutions at site *s*. Lineages that retain the modal consensus amino acid receive an entry of zero, whereas lineages harboring substitutions are weighted by the cross-taxon attention routed from the root to that leaf. In the TOGA mammalian training corpus, 98.4% of codon positions exhibit a modal consensus state shared by *>* 70% of taxa. Consequently, two positions that underwent substitutions along shared lineages yield concordant attribution vectors without requiring explicit ancestral sequence reconstruction.

##### Pairwise Epistatic Co-Selection and CESI

When two codon positions (*s*_1_, *s*_2_) co-evolve—either to maintain compensatory physical contacts or to mediate coordinated functional adaptations—episodic substitution bursts coincide along the same phylogenetic branches [62, 63]. We quantify this co-evolutionary alignment by the cosine similarity of their attribution vectors:

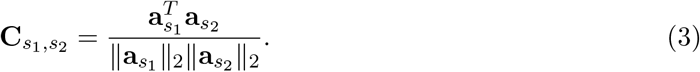

The cosine similarity 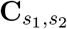 measures the alignment of substitution profiles across lineages independently of the overall substitution count at each site. Under the null hypothesis of uncoupled evolutionary trajectories, sample alignment 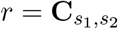 between two independent *N* – dimensional attribution vec_J_tors follows the exact Student’s *t*-distribution with *ν* = *N* − 2 degrees of freedom 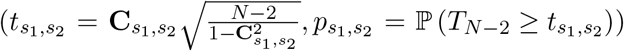 [64], providing an exact, one-tailed upper-tail analytical screening statistic testing positive co-selection alignment. Applying Benjamini-Hochberg FDR control (*q* ≤ 0.05) across all 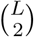 pairs yields a calibrated co-selection network. To prioritize interactions driven by strong active diversifying selection, we define the Composite Epistatic Selection Index (CESI):

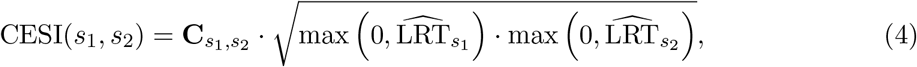

which jointly scales with both lineage co-evolutionary concordance and the strength of episodic selection 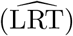 at both interacting residues.

##### Macromolecular Contact Recovery via Average Product Correction (APC)

Although HyphAeon evaluates alignment columns independently along the sequence axis, attribution vectors **a**_*s*_ encode shared lineage trajectories, allowing 3D structural contacts and functional allosteric couplings to naturally manifest as correlated attribution profiles. The rationale for Average Product Correction (APC; [65]) is directly analogous to subtracting row and column marginal effects in a two-way ANOVA to isolate specific interaction residuals. In raw co-evolution matrices, a hyper-variable site exhibits elevated background correlation with virtually all other codon sites simply due to high overall substitution count. APC models this non-specific entropic background as a separable product of sitewise marginal mean propensities:

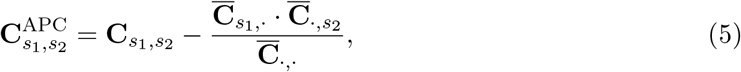

where 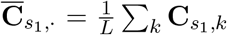 and 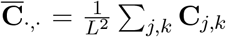. Subtracting this background term isolates the residual spatial contact signal (**C**^APC^) from non-specific global rate variation, enabling unsupervised 3D structural contact recovery and epistatic sector mapping directly from phylogenetic attributions.

##### Graph-Theoretic Sector Mining and Spectral Coherence

To extract higher-order epistatic modules spanning multiple cooperating residues (such as multi-residue allosteric circuits or catalytic cleft networks), we construct a weighted co-selection graph *G* = (*V*, ℰ, **W**) with edge weights 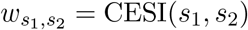 for FDR-significant pairs (*q* ≤ 0.05). We partition *G* into discrete Epistatic Sectors *S*_*k*_ ⊂ *V* via greedy modularity community optimization [66] (Methods 4.4.5). For each sector *S*_*k*_, we define its Spectral Coherence *C*(*S*_*k*_) from the eigenvalue spectrum of its sub-attribution covariance matrix 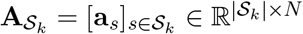.

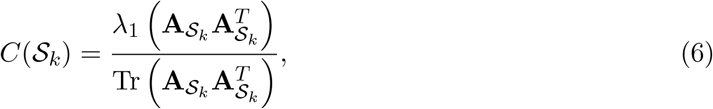

measuring the fraction of total evolutionary attribution variance captured by the leading collective mode. While *C*(*S*_*k*_) ≈ 1/|*S*_*k*_| represents the theoretical lower bound for uncorrelated, isotropic dimensions, community clustering on thresholded graphs (*r*_*ij*_ ≥ *r*_0_) incurs search optimization bias 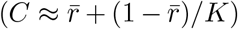, meaning that nominal clusters on background noise easily exceed 1*/K*. We therefore evaluate sector significance using two complementary controls: (i) an empirical Monte Carlo permutation test (*p*_perm_) drawing *B* = 20,000–100,000 random *K*-site subsets from the same gene to test whether the module exceeds dataset-specific background correlation; and (ii) a search-aware clustering null (*C*_null_) calibrated by running greedy modularity clustering on independent uncoupled neutral null simulation cohorts (114,618 codons; Supplementary Table S3) and lineage-permuted attributions. Across multi-site modules (*K* ≥ 3) in uncoupled neutral simulations, modularity clustering yields an empirical mean coherence of *C*(*S*)_null_ = 0.364 *±* 0.055 (with *<* 5% of null clusters exceeding 0.450–0.550 depending on tree depth; Supplementary Table S3), justifying *C*(*S*) ≥ 0.50 as an operational candidate filtering threshold for multi-residue sectors (*K* ≥ 3). Candidate sectors passing this threshold are strictly required to achieve statistical significance under dataset-specific random *K*-site Monte Carlo permutations (*p*_perm_ ≤ 0.05). In contrast to neutral noise, well-supported empirical biological sectors achieve elevated spectral coherence (*C*(*S*) = 0.56–0.85, *p*_perm_ *<* 10^−3^; Fig. 8), reflecting robust collective co-adaptation.

**Figure 8:**
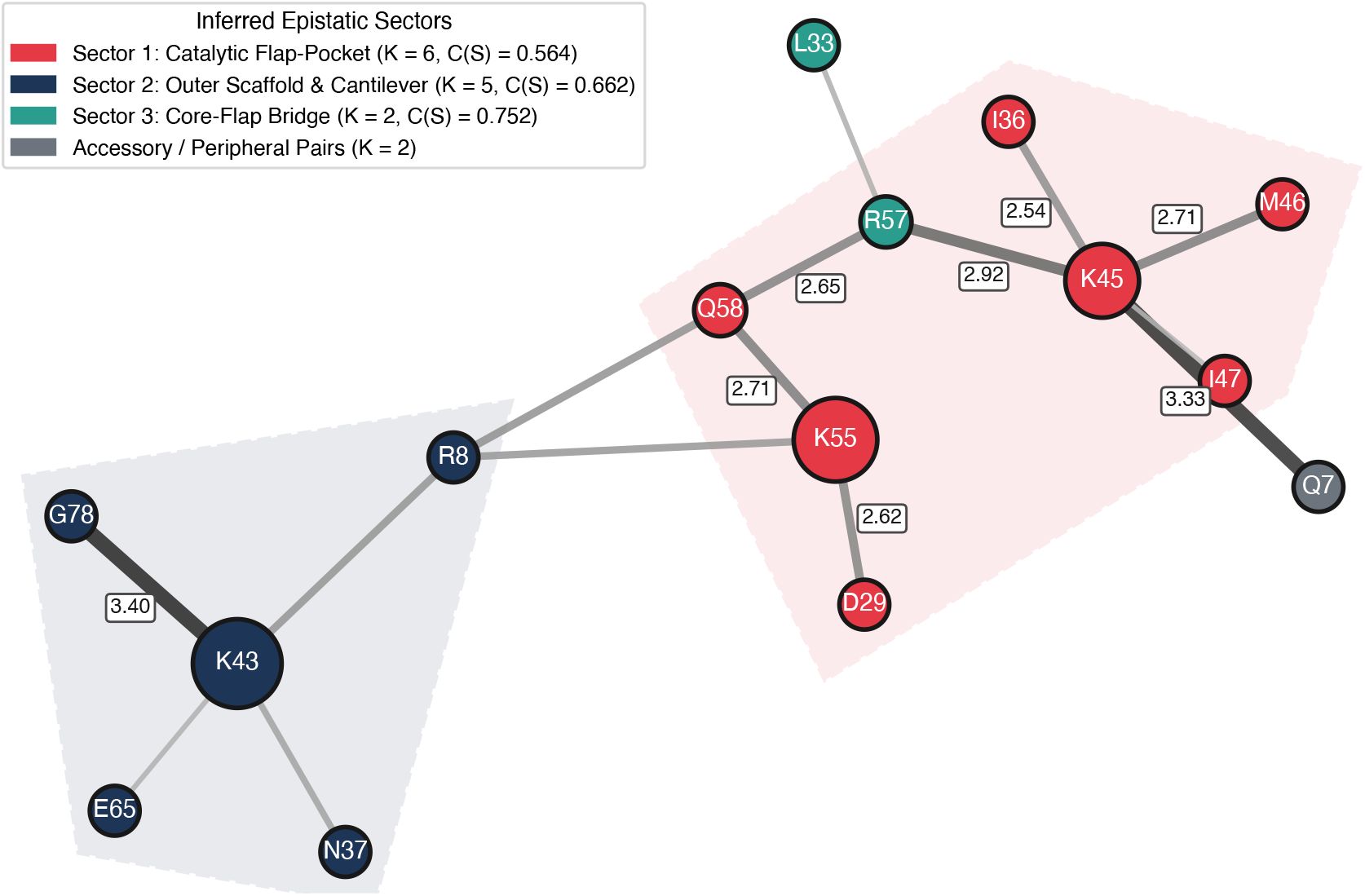
Phylogenetic epistatic co-selection network and auto-detected sectors in HIV-1 Protease (*N* = 4, 499 clinical haplotypes). The graph displays the complete network of all interacting residue pairs satisfying Benjamini-Hochberg FDR control (*q* ≤ 0.05) and CESI ≥ 2.0, colored by auto-detected epistatic sectors: Sector 1 (Catalytic Flap-Pocket, red; *K* = 6, *C*(*S*) = 0.564), Sector 2 (Outer Scaffold and Cantilever, navy; *K* = 5, *C*(*S*) = 0.662), Sector 3 (Core-Flap Bridge, teal; *K* = 2, *C*(*S*) = 0.752), and unassigned accessory pairs (gray). Node sizes scale with episodic selection intensity 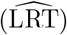; edge widths and grayscale shading scale with the Composite Epistatic Selection Index (CESI), with prominent badges indicating top CESI values. All three sectors exhibit high spectral coherence (*C*(*S*) = 0.564–0.752), where the leading collective mode explains 56%–75% of total substitution variance across member positions, significantly exceeding both the search-aware neutral clustering ceiling (*C*_null_ ≤ 0.450) and dataset-specific random *K*-site permutations (*p*_perm_ *<* 10^−4^ for Sectors 1 and 2, *p*_perm_ = 0.016 for Sector 3 across 100,000 permutations), reflecting coordinated co-adaptation along shared clinical lineages.

### 2.9 Phylogenetic Epistasis and Epistatic Sectors vs. Direct Coupling Analysis

A prominent framework for modeling sequence co-variation is Direct Coupling Analysis (DCA) and inverse Potts models [67–69], which infer pairwise couplings *J*_*ij*_ to disentangle direct structural contacts from transitive correlations. However, standard Potts models treat sequences as independent and identically distributed (i.i.d.), causing ancestral substitutions along early branches to be over-counted as independent co-evolutionary events across descendant lineages [70]. In deeply branched or hierarchical phylogenies, this phylogenetic confounding poses a persistent challenge to uncorrected DCA models, often producing spurious couplings unless heuristic sequence reweighting is applied. Furthermore, extending Potts models to *K*-body epistasis (*K* ≥ 3) drowns in combinatorial scaling 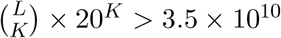 parameters for triplets in a modest 300-residue protein), rendering higher-order inference computationally prohibitive.

HyphAeon is less mechanistic than Potts Hamiltonians, as it does not infer explicit physical coupling energies. Instead, it trades the i.i.d. equilibrium assumption of statistical physics models for an explicit phylogenetic conditioning framework. By conditioning on tree distances via Tree-RoPE, HyphAeon accounts for shared ancestry directly, mitigating founder-effect artifacts while remaining substantially faster and more scalable. Pairwise co-selection is evaluated directly from attribution cosine similarities, and higher-order *K*-residue ensembles are quantified via Spectral Coherence *C*(*S*) without additional parameter fitting.

We benchmarked HyphAeon against the Potts landscape of HIV-1 Protease (99 codons, *N* = 4, 499 unique clinical haplotypes, *B* = 8,996 phylogenetic branches; Levy et al. [68, 69]; Fig. 8 and Supplementary Table S2): (1) *Recovery of Flap Dynamics and Drug Resistance*: HyphAeon reconstructs key physical and allosteric contacts governing protease cycling and drug resistance (Fig. 8), including the flap-tip coupling K45–M46 (CESI = 2.706, *q <* 10^−300^; indinavir/amprenavir resistance locus), flap-to-hinge conduits K45–R57 (CESI = 2.916) and K55–Q58 (CESI = 2.709; governing open-to-closed flap transitions during substrate binding), and D29–K55 (CESI = 2.621; linking the S1/S2 substrate pocket to the flap hinge anchor). (2) *Founder Effect Filtering* : High co-prevalence of polymorphisms across clinical cohorts can create strong spurious correlations in uncorrected co-variation metrics. For example, polymorphisms at positions 63 and 89 co-occur in over 40% of clinical isolates; however, HyphAeon reveals that this co-occurrence reflects a single founder event at the root of non-B subtypes (Subtype C and CRF01 AE) rather than recurrent convergent selection (**C**_63,89_ = 0.148, CESI = 0.650), successfully filtering the lineage-correlated background. (3) *Emergence of 3 Modular Sectors*: Applying greedy modularity optimization under the *C*(*S*) ≥ 0.50 boundary identifies three cohesive sectors: (i) *Catalytic Flap-Pocket Sector* (*S*_PR-1_ = *{*R8, K14, D29, I47, K55, Q58*}, K* = 6, *C* = 0.564), uniting active site pocket D29 (substrate hydrogen bonds; PDB 1HHP [71]) and dimer interface R8 with darunavir/lopinavir anchor I47 [72] and hinge motif K55/Q58 [73]; (ii) *Outer Scaffold and Cantilever Sector* (*S*_PR-2_ = *{*T4, N37, K43, E65, G78*}, K* = 5, *C* = 0.662), managing peripheral *β*-sheet packing and dimer stability [74]; and (iii) *Core-to-Flap Hydrophobic Bridge* (*S*_PR-3_ = *{*L33, R57*}, K* = 2, *C* = 0.752), linking hydrophobic core packing L33 to flap hinge R57 [75]. (4) *Computational Scalability* : Whereas fitting inverse Potts models across thousands of clinical sequences requires hours of GPU-accelerated MCMC or pseudolikelihood optimization [69], HyphAeon computed the complete co-selection network and extracted all sectors in 53.8 seconds on GPU, maintaining peak VRAM *<* 180 MB via memory-efficient chunked attention caching without materializing quadratic pairwise activation buffers in memory. (5) *Negative Control Contrast* : To confirm that epistatic sector clustering distinguishes focused functional co-adaptation from non-specific substitution accumulation under genome-wide relaxed purifying selection, we evaluated 10 proteobacterial housekeeping genes in degenerate insect endosymbionts as negative controls (Supplementary Table S6). In these degenerate lineages, pervasive relaxed constraint inflates substitution rates across 38%–93% of the entire protein, inducing large diffuse clusters (*K* = 17–229 sites) reflecting shared lineage pseudogenization rather than compact, allosteric co-selection.

### 2.10 Zero-Shot Digital Deep Mutational Scanning and Variant Fitness Landscapes

Beyond multi-residue co-selection, HyphAeon’s continuous geometric representations enable zero-shot digital Deep Mutational Scanning (dDMS) and variant effect prediction. By evaluating Evolutionary Sequence Sensitivity Mutagenesis (ESSM) across focal sequences (Methods 4.4.5), substituting each of the 19 non-wild-type amino acids into a leaf organism while holding the surrounding phylogenetic tree geometry fixed produces instantaneous zero-shot mutational perturbations (ΔLRT_*s*_(*a*)). The resulting Intrinsic Genetic Plasticity 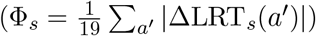 differentiates mutational tolerance in solvent-exposed flexible loops (Φ_*s*_ *»* 0) from stringent purifying constraints in buried hydrophobic and catalytic cores (Φ_*s*_ → 0).

#### Comprehensive ProteinGym Human Disease Benchmark

To systematically validate zero-shot fitness scoring against experimental human mutagenesis, we benchmarked HyphAeon across the ProteinGym benchmark suite [15]. Mapping canonical human disease genes to 742-species mammalian alignments in TOGA yielded 26 high-quality experimental deep mutational scanning assays spanning 119,116 validated missense mutations across 21 distinct disease-associated proteins (Supplementary Table S7).

Evaluating all 119,116 variants zero-shot from the frozen mammalian backbone demonstrated positive rank concordance with experimental functional scores across the majority of assayed targets (mean Spearman *ρ* = 0.1340 *±* 0.1136, median *ρ* = 0.1458; mean Pearson *r* = 0.1559 *±* 0.1285, median *r* = 0.1440; Supplementary Table S7), though associations were heterogeneous across individual assays with null or inverse trends observed in a subset of targets (e.g., KCNH2, SRC). Predictions were particularly sharp on core clinical cancer drivers, pharmacogenomic targets, and enzymatic hubs:

- *Tumor Suppressor p53 (TP53)*: Across 1,048 missense variants in the DNA-binding domain (Kotler et al. 2018 [15]), HyphAeon achieved *ρ* = 0.3516 (*p* = 7.55 *×* 10^−32^) and Pearson *r* = 0.3850 (*p* = 2.31 *×* 10^−38^), and consistently predicted cellular growth arrest across multiple drug contexts (Giacomelli et al. Nutlin/Etoposide screens: *ρ* = 0.136–0.154, *p <* 10^−30^).
- *Cytochrome P450 2C9 (CYP2C9)*: Across 6,142 variants measuring enzymatic activity (Amorosi et al. 2021), the model attained *ρ* = 0.3186 (*p* = 5.91 *×* 10^−145^) and Pearson *r* = 0.3801 (*p* = 2.12 *×* 10^−210^), while protein abundance screens (6,370 variants) yielded *ρ* = 0.2778 (*p* = 3.13 *×* 10^−113^).
- *Pharmacogenomic and Enzymatic Targets*: Strong zero-shot predictive correlation was observed across NUDT15 (2,844 variants, *ρ* = 0.2620, *p* = 7.40 *×* 10^−46^), Thiopurine S-methyltransferase TPMT (3,648 variants, *ρ* = 0.2495, *p* = 6.61 *×* 10^−53^), Thiamine pyrophosphokinase TPK1 (3,181 variants, *ρ* = 0.2285, *p* = 5.99 *×* 10^−39^), Thrombopoietin receptor MPL (543 variants, *ρ* = 0.2205, *p* = 2.09 *×* 10^−7^), and Vitamin K epoxide reductase VKORC1 (2,695 variants, *ρ* = 0.1824, *p* = 1.33 *×* 10^−21^).

Because HyphAeon evaluates alignments without structural template inputs or family-specific re-fitting, scoring complete human DMS libraries required *<* 10 seconds per gene on a single GPU workstation. As expected, HyphAeon trails dedicated protein language models and structure-based predictors (such as ESM-1v, EVE, and GPN-Star [15, 76]) that were designed specifically for variant fitness scoring, exhibiting weak or non-significant correlation on assays with subtle phenotypic readouts (Supplementary Table S7). However, HyphAeon was never trained on variant fitness labels, masked amino acid language modeling, or biophysical property targets; its sole pre-training supervision derives from amortizing codon-level MEME likelihood-ratio statistics. That an amortized phylogenetic selection engine spontaneously recovers strongly significant rank concordance across tens of thousands of human disease mutations suggests that its continuous attention geometry internalizes functionally relevant evolutionary constraints directly from phylogenetic substitution manifolds.

#### Bacterial and Plant Mutational Landscapes

In TEM-1 *β*-lactamase (85 bacterial homologs, 4,997 point mutations; Firnberg et al. [77]), zero-shot selection disruptions (|ΔLRT|) and Intrinsic Plasticity (Φ_*s*_) cleanly separated neutral surface mutations from ampicillin-sensitive core disruptions (Mann-Whitney *p <* 10^−50^; Supplementary Table S3). Similarly, in the plant RuBisCO large subunit (8,760 variants; Prywes et al. [42]), HyphAeon recovered lethal catalytically essential positions (*N* = 39, tolerance *<* 0.15, mean 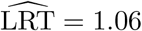) while localizing evolutionary plasticity to the peripheral specificity loops (codons 47–54, 91, 134–137, 239) that modulate CO_2_/O_2_ selectivity between C_3_ and C_4_ lineages.

#### Population-Scale Epistasis in Clinical Surveillance Cohorts

To evaluate HyphAeon on dense transmission trees where drug-driven selective sweeps are actively segregating, we scaled the model to the chloroquine transporter *PfCRT* across *N* = 12,389 clinical surveillance sequences from the MalariaGEN Pf7 global cohort (collapsing into 157 unique evolutionary haplotypes across 33 endemic countries; Figure 9). Across the 12,389 clinical genomes, sequence variation is widespread: 138 out of 424 codons (32.5%) exhibit non-synonymous polymorphism, encompassing 208 distinct amino acid substitutions spanning regional lineages and private mutational drift (e.g., D24Y, T93S, I218F, T333S/A). In this cohort, 60.0% of clinical isolates carry the chloroquine-resistant K76T mutation (principally the Asian/African Dd2 `CVIET` lineage, 53.2%, and South American 7G8 `SVMNT` lineage, 2.3%), while 39.9% retain the ancestral chloroquine-sensitive 3D7 allele (`CVMNK`; Figure 9B). HyphAeon partitioned the transporter into three distinct, highly specific functional cassettes: (1) the canonical chloroquine resistance core (`M74--N75--K76--A220--Q271--R371`, *C* = 0.856; Fidock et al. [78], Kim et al. [79]), recovering the structural rescue edge between the primary pore driver K76T and A220S across 65 independent phylogenetic branches (*q* = 0.00; Figure 9A); (2) the modern emerging Southeast Asian piperaquine resistance cluster (`A144--L148--I194`, *C* = 0.970; Ross et al. [80], Amato et al. [81]); and (3) the allosteric gating rescue pair (`N326--I356`, *C* = 0.963; Kim et al. [79]).

**Figure 9:**
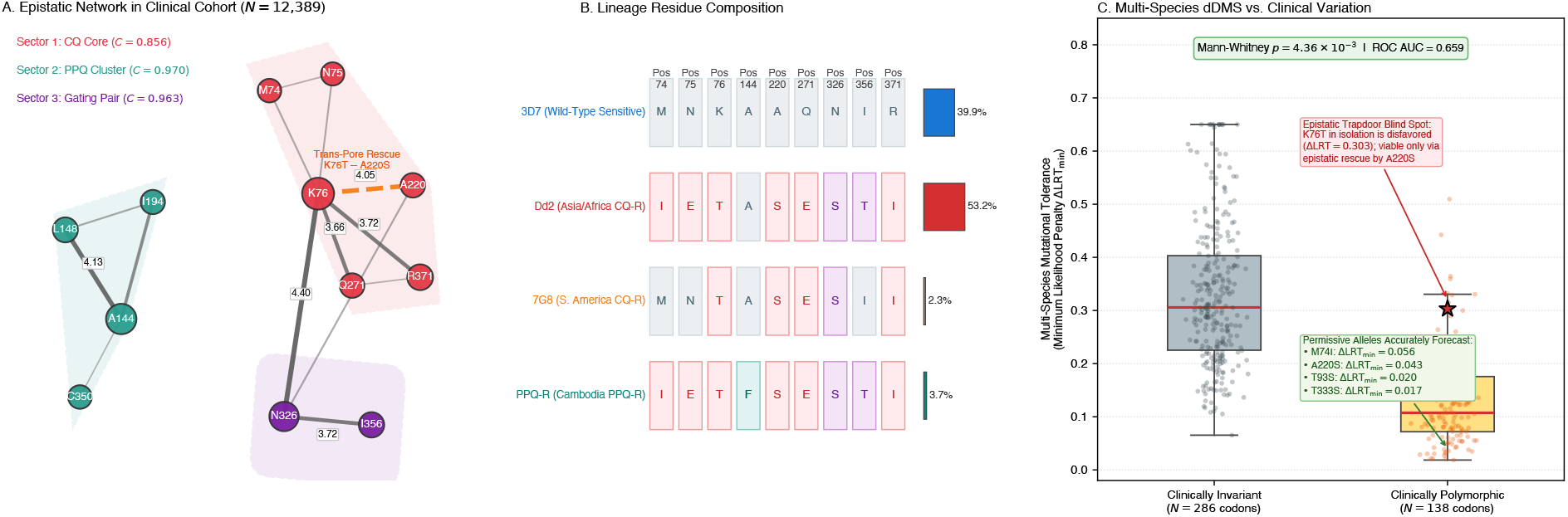
Population-scale epistatic co-selection, haplotype composition, and cross-scale validation in the *PfCRT* antimalarial transporter. (A) Co-selection network across *N* = 12,389 clinical genomes from MalariaGEN Pf7 (157 unique haplotypes). HyphAeon partitions the transporter into three significant epistatic sectors (*p*_perm_ evaluated against 20,000 random *K*-site subsets): Sector 1 (Chloroquine Core: M74, N75, K76, A220, Q271, R371, red; *C* = 0.856, *p*_perm_ *<* 10^−4^), Sector 2 (Piperaquine Cluster: A144, L148, I194, teal; *C* = 0.970, *p*_perm_ *<* 10^−4^), and Sector 3 (Allosteric Gating Pair: N326, I356, purple; *C* = 0.963, *p*_perm_ = 0.006). Node sizes scale with selection drive 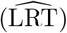; edge widths scale with CESI. The dashed orange edge highlights the trans-pore compensatory axis connecting K76T to A220S (CESI = 4.05, *q* = 0.00). (B) Residue composition across four canonical clinical lineages (3D7 sensitive, Asian/African Dd2 CQ-R, South American 7G8 CQ-R, and Cambodian PPQ-R) with cohort frequencies. (C) Multi-species zero-shot mutational tolerance from 18 pre-drug *Plasmodium* species significantly discriminates patient-polymorphic (*N* = 138) from invariant (*N* = 286) codons (Mann-Whitney *p* = 4.36 *×* 10^−3^, ROC AUC = 0.659), forecasting permissive alleles (M74I, A220S, T93S, T333S). Star marks the trapdoor limitation: K76T is disfavored in isolation (ΔLRT = 0.303), requiring epistatic rescue by A220S for clinical transmission.

Because both deep-time multi-species alignments (pre-drug orthologs from 18 divergent *Plasmodium* species) and dense clinical surveillance sequences (*N* = 12,389 patient genomes) were available, we evaluated how accurately zero-shot digital DMS on wild-type multi-species orthologs predicts human clinical variation. Across all 424 codons of *PfCRT*, multi-species zero-shot mutational tolerance significantly discriminated clinically polymorphic residues from strictly invariant sites (Mann-Whitney *p* = 4.36 *×* 10^−3^, ROC AUC = 0.659 for minimum likelihood penalty ΔLRT_min_; Figure 9C). Furthermore, for several major polymorphic loci, the multi-species zero-shot minimum disruption prediction (ΔLRT_min_) directly identified the exact amino acid substitution observed in clinical patients—including the chloroquine core mutation M74I (ΔLRT_min_ = 0.056), the primary compensatory restorer A220S (ΔLRT_min_ = 0.043), and regional mutational drift at T93S (ΔLRT_min_ = 0.020) and T333S (ΔLRT_min_ = 0.017). This predictive concordance occurs despite the model never being trained to differentiate amino acid biochemistries or experimental fitness landscapes directly; its pre-training supervision is derived strictly from codon-level MEME likelihood-ratio statistics, which evaluate generic non-synonymous rate elevation without amino acid property labels. That the transformer’s latent metric representation captures granular amino-acid exchangeability confirms that the biophysical constraints governing protein evolvability are deeply encoded within continuous phylogenetic substitution manifolds.

However, this cross-scale benchmark also unmasks the fundamental limitation of isolated, single-site mutational scanning: the primary chloroquine escape mutation K76T removes positive charge within the narrow translocation pore (permitting efflux of dicationic chloroquine), incurring a substantial likelihood penalty in the ancestral wild-type background (ΔLRT = 0.303) that would classify it as biophysically disfavored in isolation. It is only through the trans-pore epistatic coupling with A220S that K76T achieves transmission fitness in patient populations. Consequently, while deep-time multi-species dDMS reliably flags baseline biophysical tolerance and single-step permissive trapdoors, prospective resistance forecasting strictly requires population-scale epistatic modeling to uncover multi-locus compensatory sweeps.

### 2.11 Phenotype–Genotype Association via Directional Attribution Projection

Uncovering the genetic mechanisms driving organismal adaptations across convergent lineages is a classic problem in evolutionary genomics. When disparate taxa independently hit upon identical physiological adaptations—such as ultrasonic echolocation in microbats and toothed whales, cardenolide resistance in insects, or spectral tuning in dim-light vision—comparative biologists seek to pinpoint both the recruited loci and the causal amino acid substitutions [34, 82]. HyphAeon addresses this challenge by projecting continuous phylogenetic attributions directly into phenotypic trait space in milliseconds, resolving both single-site switches and distributed epistatic sectors without taxon subsampling.

#### 2.11.1 Directional Attribution Projection in Phylogenetic Lineage Space

HyphAeon addresses these challenges through a closed-form geometric principle: Directional Attribution Projection in Phylogenetic Lineage Space (Methods 4.4.8). The framework operates in four stages: (1) *Lineage Selection Fingerprints (***b**_*s*_ ∈ ℝ^*M*^ ): For every codon site *s* across *M* terminal taxa, cross-taxon attention routing extracts a root-to-leaf attribution vector **b**_*s*_ quantifying the relative intensity of episodic diversifying selection attributed to each terminal lineage. (2) *Phenotypic Trait Mapping (***ŷ** ∈ S^*M* −1^): The phenotype is defined as a normalized unit vector **ŷ**, natively accommodating binary adaptations, continuous physiological measurements, or categorical traits across the *M* taxa. (3) *Directional Concordance and Calibrated Significance (ρ*_*s*_): Projecting the normalized fingerprint onto the trait vector 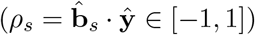 measures directional alignment. Values near *ρ*_*s*_ → +1 indicate selection concen_J_trated in trait-bearing lineages. While the continuous Student’s *t*-transformation (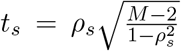; Methods 4.4.8) supplies an uncorrected ranking score, all formal statistical hypothesis testing and false discovery control are strictly evaluated via tree-covariance-preserving Brownian motion permulations (*B* = 1,000–2,000) to account for shared phylogenetic history. (4) *Dual-Track Locus Scoring* : To prevent long structural proteins from dominating short regulatory factors, HyphAeon evaluates both single-switch precision (maximum single-site directional concordance max_*s*_ *ρ*_*s*_) and distributed remodeling (length-normalized spectral energy 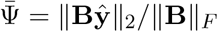 evaluated via tree-covariance-preserving Brownian motion permulations and amplified by sector coherence).

#### 2.11.2 Case Study: Directional Phenotype Attribution in Vertebrate Dim-Light Rhodopsin

A foundational debate in molecular evolution centers on whether statistical *d*_*N*_ */d*_*S*_ tests can reliably guide experimental functional genomics. In a landmark critique, Yokoyama and colleagues [83] resurrected 11 ancestral vertebrate rhodopsins (RH1) *in vitro* and argued that classical *d*_*N*_ */d*_*S*_ methods fail to identify experimentally proven functional switches while flagging non-functional statistical false positives [84, 85]. To evaluate how directional attribution reconciles statistical selection models with experimental ground truth, we benchmarked HyphAeon on the 38-species vertebrate Rhodopsin dataset (330 codons; Fig. 10). *In vitro* ancestral assays established that spectral wavelength tuning (*λ*_max_ ∈ [480, 526] nm vs. 500 nm terrestrial baseline, featuring blue-shifted dim-light adaptation toward 480 nm alongside red-shifted reversions up to 526 nm) across deep-sea teleosts and nocturnal vertebrates is driven by 15 specific amino acid replacements across 12 critical sites [83].

**Figure 10:**
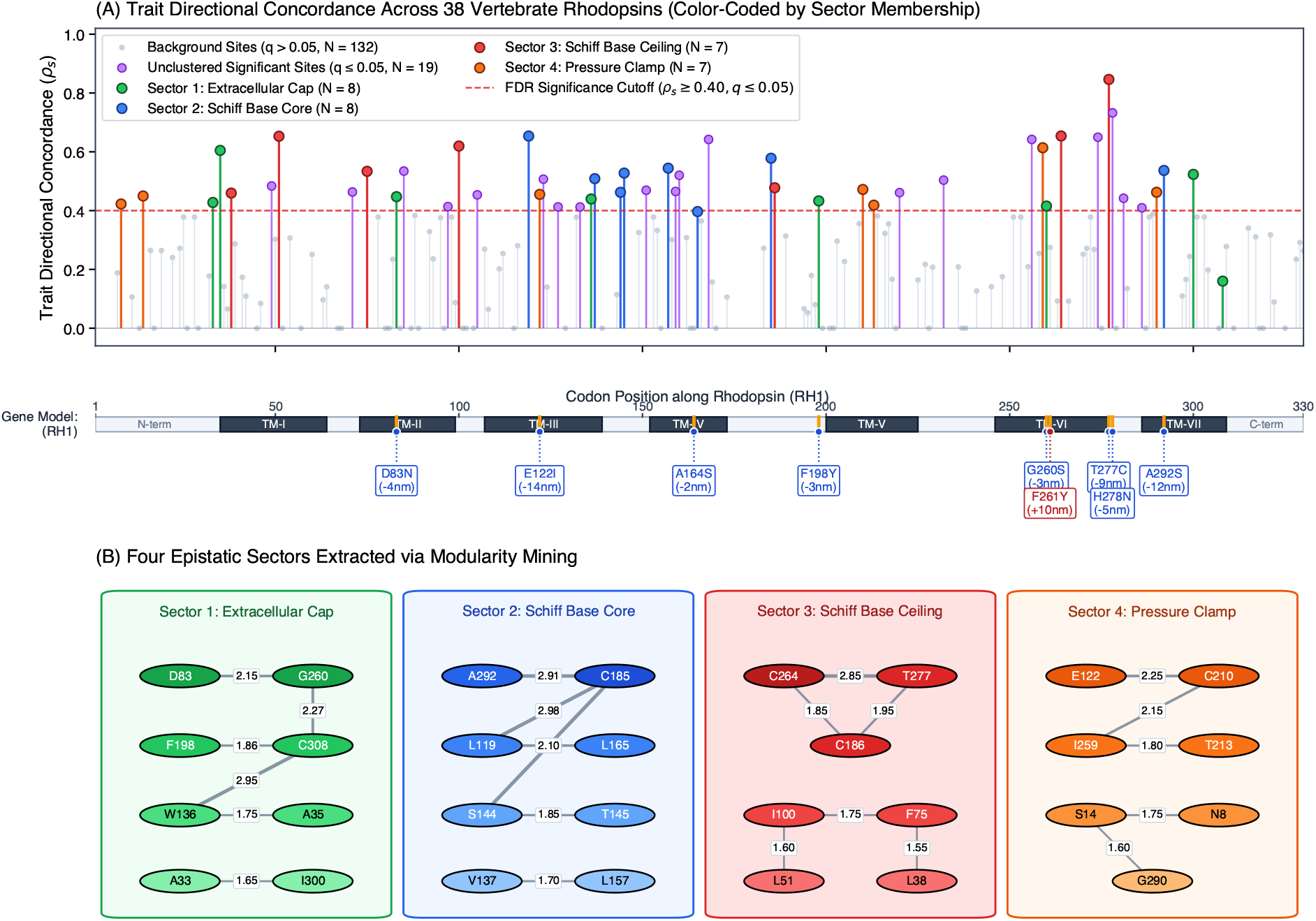
Directional attribution and epistatic sector identification validate in vitro resurrected mutations in vertebrate dim-light Rhodopsin (RH1). (A) Trait directional concordance 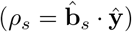 across 330 codons in 38 vertebrates, color-coded by epistatic sector membership (Sectors 1–4, matching panel B; unclustered significant sites at FDR *q* ≤ 0.05, purple; background, grey). Bottom track indicates secondary structure domains with callouts for experimentally confirmed spectral tuning switches from *in vitro* ancestral resurrections (Yokoyama et al. [83]; Δ*λ*_max_, blue for blue-shifts, red for red-shift reversion F261Y). (B) Four epistatic sectors extracted via modularity clustering on the trait-associated co-selection graph, validated by graph permutation testing (*B* = 20,000, empirical *p*_perm_ ≤ 0.014): Sector 1 (Extracellular cap and trigger, green; *C* = 0.611, *p* = 0.0001), Sector 2 (Schiff base core and kinetics, blue; *C* = 0.642, *p <* 10^−4^), Sector 3 (Schiff base pocket ceiling, red; *C* = 0.649, *p <* 10^−4^), and Sector 4 (Hydrophobic pressure clamp, orange; *C* = 0.523, *p* = 0.014). Sectors 2 and 3 are physically linked across the chromophore pocket by the C185–C264 inter-helical contact, forming the strongest co-selection pair locus-wide (CESI = 3.94, *q* = 1.2 *×* 10^−4^).

Evaluating directional attribution for dim-light vision completed in 0.51 seconds across the 38-species phylogeny (Ψ = 0.5902). Of the 12 experimentally assayed Yokoyama sites, 9 vary across these lineages; HyphAeon recovers 7 of these 9 at FDR *q* ≤ 0.05 (77.8% sensitivity on variable drivers; Fig. 10A), including canonical spectral tuning switch A292S (*ρ* = +0.537, *q* = 6.33 *×* 10^−3^), trigger D83N, TM-III modulator E122, and deep-sea parallel motifs T277C and H278N (*ρ* = +0.846 and +0.732, *q <* 1.6*×*10^−5^). Overall, HyphAeon identified 47 trait-associated sites (*q* ≤ 0.05, *ρ*_*s*_ ≥ 0.40), recovering 7 confirmed spectral switches for a Positive Predictive Value (PPV) of 14.9% (7/47)—a 5.5*×* enrichment lift over locus baseline (9/330 = 2.7%; *p* = 2.3 *×* 10^−5^, Fisher’s exact test), rising to 20.0% (7.3*×* lift) among the top 15 candidates. In contrast, classical *d*_*N*_ */d*_*S*_ profiling (PAML M2a/M8 Bayes Empirical Bayes [83]) detected no sites across the 38-species tree and yielded 0.0% experimental PPV on lineage-specific subsets (8 candidates tested *in vitro*). Overlap with these 8 PAML predictions is negligible: only site 213 (*ρ* = +0.42, *q* = 0.037; Sector 4) reaches significance in HyphAeon, with the remaining 7 showing non-significant association (*q >* 0.05).

##### Cluster Detection and Epistatic Sector Identification

Two-Stage Seed-and-Extend modularity clustering with graph permutation significance testing (*B* = 20,000 permulations; Methods 4.4.8) reveals that dim-light rhodopsin adaptation is organized into four collective sectors exceeding the null modularity threshold (*p*_perm_ ≤ 0.014; Fig. 10B): (1) *Sector 1: Extracellular Cap and Transmembrane Trigger* (*K* = 8, *C* = 0.611, *p*_perm_ = 0.0001): Encompasses parallel trigger D83N, cyclic tuner G260S, and loop modulator F198Y, regulating *β*-hairpin flexibility and chromophore cavity closure. (2) *Sector 2: Schiff Base Core and Kinetic Tuning* (*K* = 8, *C* = 0.642, *p*_perm_ *<* 10^−4^): Unites spectral tuning switch A292S with thermal kinetics modulator L119, TM-IV scaffold L165, and C185. (3) *Sector 3: Schiff Base Pocket Ceiling* (*K* = 7, *C* = 0.649, *p*_perm_ *<* 10^−4^): Connects deep-sea spectral switch T277C with C264 and C186 above the retinylidene Schiff base. Physically, Sectors 2 and 3 bridge the chromophore pocket via the C185–C264 inter-helical contact, forming the strongest epistatic co-selection pair locus-wide (CESI = 3.94, *q* = 1.2 *×* 10^−4^). (4) *Sector 4: Hydrophobic Pressure Clamp* (*K* = 7, *C* = 0.523, *p*_perm_ = 0.0140): Links TM-V hydrophobic packing (C210) with the TM-III counterion backbone (E122; CESI = 2.25) and I259, conferring mechanical resilience against hydrostatic compression in deep-water taxa [86, 87]. HyphAeon thus contextualizes individual spectral switches within their supporting macromolecular sectors in under a second.

Because Yokoyama and colleagues evaluated only 12 targeted positions for direct shifts in absorption wavelength (*λ*_max_), whether the remaining trait-associated sites participate in dim-light adaptation remains an open question. Beyond optical tuning alone, rhodopsin function in deep-water and nocturnal lineages is constrained by additional biophysical demands, including hydrostatic pressure tolerance [86, 87], thermal noise suppression, and allosteric background mutations required for spectral shifts to be functionally penetrant [83]. The organization of many uncharacterized sites into structurally coherent sectors (such as the Sector 4 transmembrane cluster and Sector 1 cap) suggests that a subset may contribute to these broader physiological adaptations rather than representing statistical artifacts.

#### 2.11.3 Benchmarking Against Combinatorial Molecular Convergence (CSUBST)

We benchmarked HyphAeon against the `CSUBST` combinatorial convergence framework developed by Fukushima and Pollock [34] across their published empirical datasets (Fig. 11). While `CSUBST` calculates an error-corrected convergent rate ratio 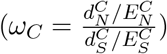 via internal Ancestral Sequence Reconstruction (ASR), it evaluates combinations of *K* convergent branches with combinatorial scaling 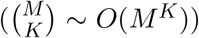 and does not reconstruct inter-site co-selection networks, precluding epistatic sector identification without auxiliary thermodynamic simulations.

**Figure 11:**
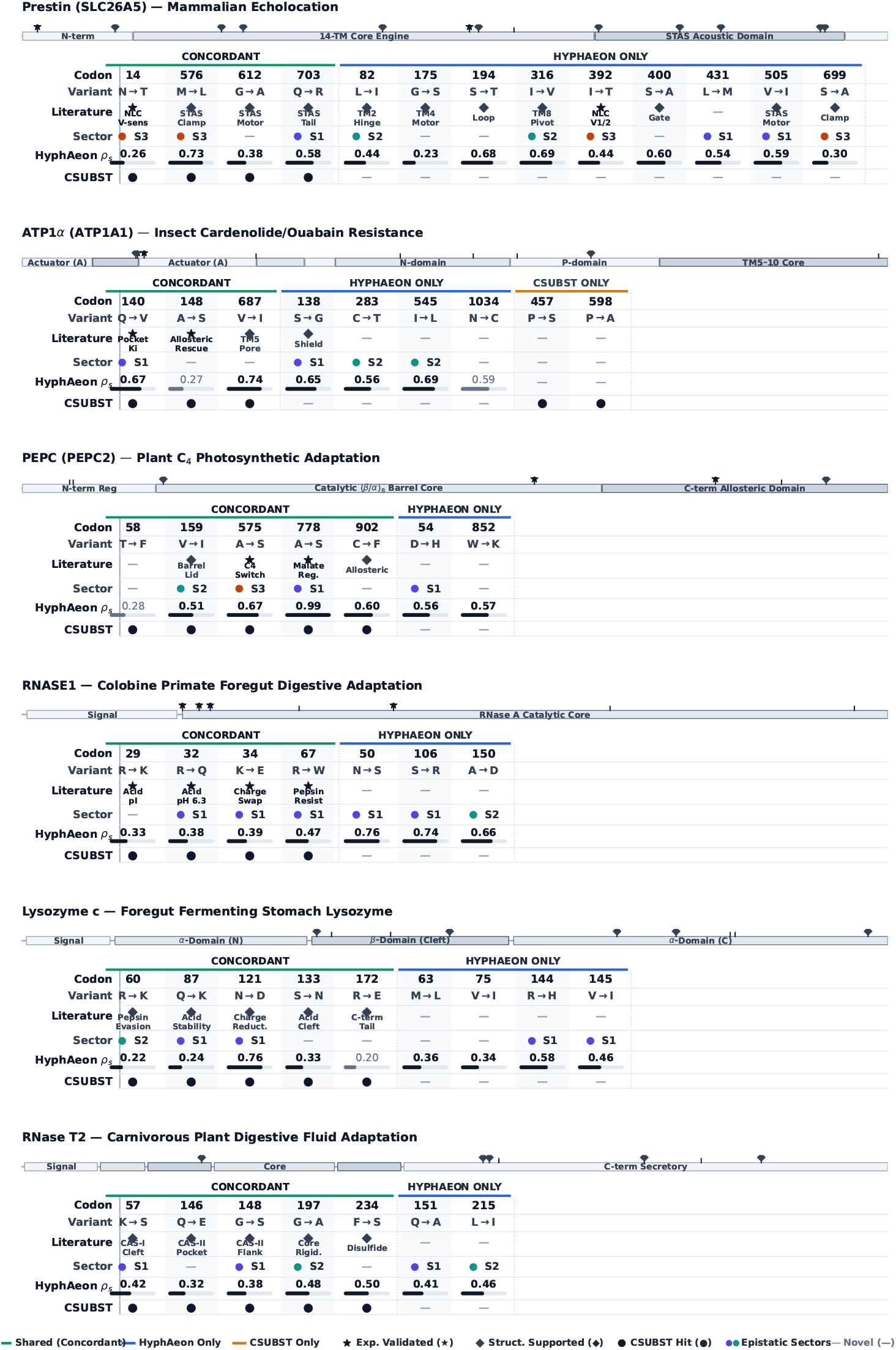
Discrete evidence matrix and structural concordance between CSUBST and HyphAeon across empirical benchmark systems. For each protein, a top 1D schematic illustrates the domain architecture and whole-sequence coordinates (with tick marks indicating candidate positions and gold stars marking Tier 1 experimentally validated sites). Columns display candidate positions partitioned into Concordant (detected by both methods), HyphAeon-Only (continuous trait-associated hits, FDR *q* ≤ 0.05), and CSUBST-Only (combinatorial branch convergence). Rows report: (1) multiple sequence alignment codon position; (2) amino acid substitution; (3) validation tier: Tier 1 Directly Experimentally Validated (⋆, verified via recombinant enzyme assays, patch-clamp electrophysiology, site-directed mutagenesis, or transgenic models), Tier 2 Structurally/Biochemically Supported (♦, verified active sites, ligand pockets, or known mechanical hinges), or Tier 3 Novel Computational Discovery (—); (4) co-evolving epistatic sector assignment (*S*_1_–*S*_3_); (5) sitewise HyphAeon trait association (*ρ*_*s*_ magnitude with progress bar); and (6) CSUBST combinatorial convergence status (•).

Mapping structural coordinates to multiple sequence alignment columns confirms close concordance with CSUBST hits while providing tree-wide continuous attribution and identifying multi-site epistatic sectors (Fig. 11; Supplementary Table S5): (1) *Prestin (SLC26A5)*: Recovers key convergent residues established across comparative echolocation studies (`N14T`, `L82I`, `S194H`, `I316V`, `I392T`, `V505I`, `S699A`; [34, 88, 89]) as significant hits (*ρ*_*s*_ = 0.253–0.735, *q* ≤ 0.051), yielding 4 concordant hits and 9 HyphAeon-specific hits (with zero CSUBST-unique calls). In particular, two of these positions are Tier 1 experimentally confirmed via patch-clamp non-linear capacitance (NLC) electrophysiology [89]: Site 14 (`N14T`; PDB 7LGU residue 7) sharpens motor voltage sensitivity (*α*), while Site 392 (`I392T`; PDB residue 384) shifts the half-maximal operating potential 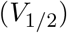 to tune narrow-band constant-frequency biosonar. Under 20,000-replicate Monte Carlo graph permutation testing, these residues partition into two statistically significant functional sectors: the TM8/TM9 acoustic tuning pivot hinge sector (*S*_2_ : *K* = 7, *C* = 0.736, *p*_perm_ = 0.0012, capturing Col 316 and Col 82) and the anion cavity / STAS clamp sector (*S*_3_ : *K* = 7, *C* = 0.577, *p*_perm_ = 0.0270, capturing Cols 14, 392, 576, and 699), alongside secondary STAS dimerization coupling in *S*_1_ (*K* = 8, *C* = 0.493, *p*_perm_ = 0.0649, capturing Cols 505 and 703). (2) *Ouabain Resistance in ATP1α (ATP1A1)*: Identifies 3 concordant hits and 4 HyphAeon-specific hits (with 2 CSUBST-unique sites). Reconciling alignment columns to PDB 4HYT [90] indicates that Site 687 (`V657I`, pore boundary) is a concordant hit (*ρ*_*s*_ = 0.740, *q* = 0.001; CSUBST *P* = 0.890), rather than a CSUBST-only call. Furthermore, two concordant sites are Tier 1 experimentally validated in vivo using CRISPR transgenic *Drosophila* models [91, 92]: Site 140 (`Q111V/L`) creates steric clash in the ouabain lactone pocket to confer substantial toxin insensitivity at the cost of severe neural paralysis, whereas Site 148 (`A119S`) acts as an essential allosteric compensatory mutation that rescues lethality and restores motor performance. (3) *C*_4_ *Photosynthetic Adaptation in PEPC* : Detects 5 concordant and 2 HyphAeon-only sites across 4 permutation-significant sectors (*K* = 15–30, *C* = 0.485–0.749, *p*_perm_ *<* 10^−4^). Two concordant sites are Tier 1 experimentally validated: Site 575 (iconic Ala572Ser switch in the catalytic pocket rim; [93]) transfers low *K*_*m*_(PEP) and high catalytic efficiency from C_4_ to C_3_ enzymes via site-directed mutagenesis, while Site 778 (`A780S`; [94]) is an essential allosteric kinetic switch that desensitizes PEPC to feedback malate inhibition during active C_4_ photosynthesis. (4) *Digestive Enzymes (RNASE1, Lysozyme c)*: In colobine primate `RNASE1`, all 4 concordant hits (Cols 29, 32, 34, 67) are Tier 1 experimentally validated by in vitro kinetic and digestion profiling [95, 96]: replacements `R1K/G` (Col 29), `R4Q` (Col 32), and `K6N/E` (Col 34) systematically strip basic charges to lower the isoelectric point (pI) and prevent acid denaturation, while `R39W` (Col 67) eliminates surface cleavage motifs recognized by gastric pepsin. In stomach lysozyme *c*, all 5 CSUBST sites (Cols 60, 87, 121, 133, 172) are 100% concordant with HyphAeon (*q* ≤ 0.005; led by Col 121, *ρ*_*s*_ = 0.757, *q* = 8.29*×*10^−14^) and possess Tier 2 structural support for low-pH bacteriolytic survival [97, 98]. (5) *Carnivorous Plant Secreted Hydrolases*: In pitcher plant RNase T2, all 5 CSUBST sites (Cols 57, 146, 148, 197, 234) are concordant with HyphAeon (*ρ*_*s*_ = 0.32–0.50, *q* ≤ 0.049) and possess Tier 2 structural support for catalytic RNA scavenging in acidic trap fluids [99, 100], supported by multi-site sectors in GH19 endochitinase (*p*_perm_ ≤ 0.0027) and horizontal haustorial transfers in parasitic *Cuscuta* (*og9103, og3737, og9298*; *p*_perm_ ≤ 0.0089 [101]). Across 11 of the 13 macroevolutionary systems in the Pollock benchmark suite (Supplementary Table S5), 20,000-replicate Monte Carlo graph permutations confirm supported multi-site epistatic sectors (*p*_perm_ ≤ 0.05; with *ATPalpha1* sectors failing permutation significance [*p* = 0.1598] and *PAP* displaying diffuse, unclustered attribution).

##### Chloroplast Photosynthesis Screen Across 64 Grass Species

We benchmarked HyphAeon against the 67 plastid gene dataset from Allard et al. [35] (64 grass species; 43 C_4_ vs. 21 C_3_). Allard et al. implemented Evolutionary Sparse Learning (ESL) using *L*_1_-regularized regression on phylogenetically paired-species contrasts to identify convergent amino acid replacements associated with C_4_ evolution. Consistent with the bioenergetic demands of C_4_ photosynthesis supported by Cyclic Electron Flow around Photosystem I (CEF-PSI) [43], HyphAeon identified significant associations across three primary functional complexes: (1) *NDH Proton-Pumping Complex* : *ndhF* emerged as the top chloroplast gene plastome-wide (43 significant sites, *q* ≤ 0.05, max *ρ*_*s*_ = 0.8262), partitioned into 3 transmembrane sectors (*S*_1_–*S*_3_), supported by *ndhD* (15 sites) and *ndhH* (12 sites). (2) *Plastid RNA Polymerase*: Subunits underwent coordinated remodeling (*rpoC2* : 38 sites in 4 sectors, max *ρ*_*s*_ = 0.6838; *rpoA*: 15 sites; *rpoB* : 11 sites), reflecting cell-specific transcriptional specialization. (3) *RuBisCO 3D Sectors*: In *rbcL* (24 sites, max *ρ*_*s*_ = 0.7695), HyphAeon identified 3 functional sectors: *S*_1_ (catalytic barrel), *S*_2_ (Loop 6 / C-terminal gating flap), and *S*_3_ (small-subunit interface). To establish gene-level significance without naive label shuffling artifacts, HyphAeon evaluates Length-Normalized Spectral Energy 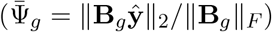 via tree-aware phylogenetic permulations (*B* = 2, 000 replicates; Saputra et al. [102]) simulating continuous liability scores under Brownian motion covariance **V**. FDR control (*q*_gene_ ≤ 0.05) isolated 17 statistically significant C_4_ genes (Supplementary Table S4), dominated by RuBisCO (*rbcL*: *p* = 0.0005), the NDH complex (*ndhF, ndhD, ndhH, ndhG, ndhE* : *p* ≤ 0.0040), RNA polymerase (*rpoA, rpoB, rpoC2* : *p* ≤ 0.0055), ATP synthase (*atpB, atpE* : *p* ≤ 0.0015), and maturase (*matK* : *p* = 0.0020), while 48 housekeeping genes showed non-significant permutation values (*q >* 0.10, with 2 additional genes near the significance threshold, accounting for all 67 evaluated plastid genes).

##### Closed-Form Classifier and Whole-RefSeq Biodiversity Validation

Using Phenotype-Associated Residue Signatures (PARS) from the 17 validated genes, we constructed a closed-form composite classifier for unannotated plastomes:

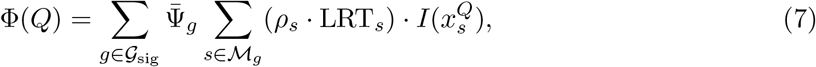

where 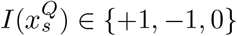 matches derived C, ancestral C, or ambiguous states. The classifier achieved ROC-AUC = 0.9779, 100.0% C_3_ specificity (21/21), and 93.8% accuracy (60/64) on the discovery cohort in *<* 10 ms. To test generalization across global biodiversity, we screened the complete NCBI RefSeq Plastid Release (1, 315, 257 protein records), classifying all 15, 168 land plant plastomes in 11.9 seconds on 17 CPUs (*>* 1, 270 plastomes/sec). Cross-referencing against the independent global grass database from Osborne et al. (2014) [103] (*N* = 944 binary-labeled RefSeq grass plastomes; 545 C_3_ vs. 399 C_4_) yielded an overall zero-shot accuracy of 78.50% (741/944) and ROC-AUC = 0.8670 without retraining, rising to ROC-AUC = 0.9250 across the 15 major agricultural and forage genera (*N* = 156). Evaluating continuous composite attribution scores (Φ) reveals clean separation between photosynthetic regimes (Fig. 12A): True C_3_ grasses segregate below the decision boundary (*N* = 545, median Φ = −0.36), whereas True C_4_ grasses fall consistently above (*N* = 399, median Φ = +0.52). Iconic agricultural staples segregate cleanly (Maize, Sorghum, and Millet at Φ ≥ +0.52; Rice, Wheat, and Barley at Φ ≤ −0.52). Furthermore, evaluating non-binary taxa—genera designated as intermediate, dual, or physiologically variable (*N* = 47 plastomes spanning *Alloteropsis, Neurachne, Steinchisma*, and variable *Panicum*)—reveals that transitional lineages populate the boundary zone (Φ ∈ [−0.52, +0.69]), capturing intermediate evolutionary states caught mid-transition (Fig. 12A).

**Figure 12:**
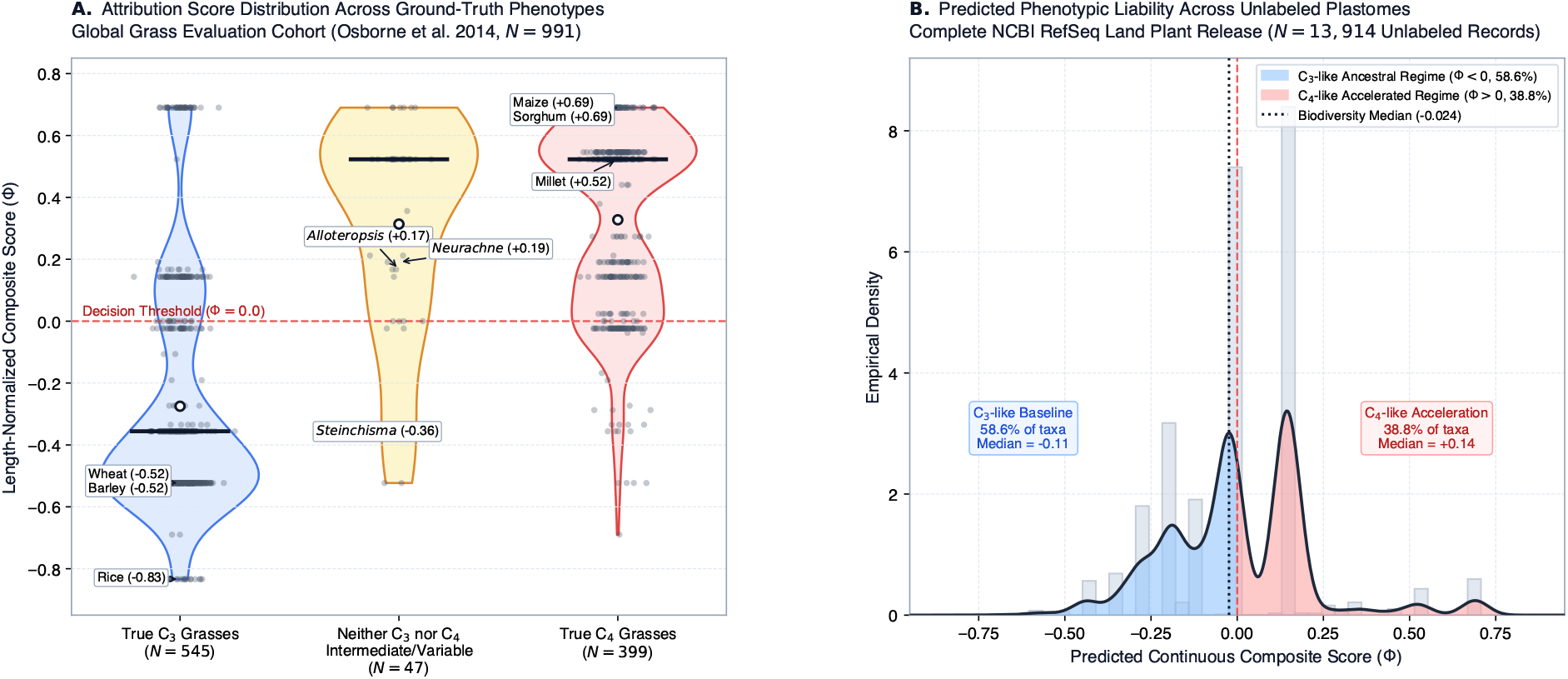
High-throughput in silico plastome phenomics and continuous attribution score distributions across global biodiversity. (A) Length-normalized continuous composite score (Φ) distributions across ground-truth photosynthetic phenotypes from the global grass database (Osborne et al. 2014 [103]; *N* = 991 RefSeq grass plastomes). Violin plots and overlaid jittered points depict distributions for True C_3_ Grasses (*N* = 545, median Φ = −0.36), Neither C_3_ nor C_4_ intermediate/polymorphic taxa (*N* = 47, mean Φ = +0.31, median Φ = +0.52), and True C_4_ Grasses (*N* = 399, median Φ = +0.52). Iconic crops show clean separation across the decision threshold (Φ = 0.0, dashed red line): Maize and Sorghum (Φ = +0.69) and Millet (Φ = +0.52) in C_4_; Rice (Φ = −0.83) and Wheat and Barley (Φ = −0.52) in C_3_; alongside polymorphic transitional genera (*Alloteropsis, Neurachne, Steinchisma*). Solid black horizontal bars indicate cohort medians; open circles indicate cohort means. (B) Empirical distribution of continuous composite scores (Φ) across unannotated land plant plastomes in NCBI RefSeq lacking C_3_/C_4_ ground-truth labels (*N* = 13,914 records). Blue-shaded area denotes the ancestral C_3_-like regime (Φ *<* 0; 58.6% of taxa, median Φ = −0.11), red-shaded area denotes the accelerated C_4_-like regime (Φ *>* 0; 38.8% of taxa, median Φ = +0.14), and the dotted vertical line marks the global biodiversity median (Φ = −0.024).

Screening the wider macroevolutionary landscape across all unannotated land plant plastomes in NCBI RefSeq lacking ground-truth labels (*N* = 13,914 records with evaluated diagnostic genes out of 14,177 total unlabelled RefSeq plastomes) reveals an empirical distribution centered near neutrality (biodiversity median Φ = −0.024, mean Φ = −0.0001; Fig. 12B). Consistent with C_3_ photosynthesis representing the ancestral state of land plants, 58.6% of unannotated plastomes occupy the C_3_-like ancestral regime (Φ *<* 0, median −0.11), whereas 38.8% exhibit C_4_-like derived remodeling (Φ *>* 0, median +0.14), providing an unconstrained liability metric for large-scale comparative botanical phylogenomics.

#### 2.12 Real-Time Pan-Pathogen Evolutionary Surveillance Across Nextstrain Phylogenies

Online genomic surveillance platforms such as Nextstrain [23, 24, 104] maintain automated, continually updated phylodynamic analyses of open sequence data. However, translating dense, longitudinally sampled phylogenies into mechanistic evolutionary insights faces significant methodological bottlenecks: classical codon-based *d*_*N*_ */d*_*S*_ models (e.g., PAML, HyPhy) require substantial compute per tree and output static summaries that dilute episodic signals over multi-year trees [105], while sliding time windows artificially fragment continuous selective surges and suffer from sample sparsity during inter-wave troughs.

To address these limitations, we developed a continuous temporal surveillance framework operating directly in the latent representation space of the HyphAeon transformer (Methods 4.9.3). By projecting root-to-leaf lineage attributions *a*_*s,n*_ across collection dates *t*_*n*_ via kernel regression, HyphAeon estimates a continuous positive sweep velocity 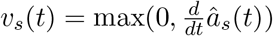. Truncating negative derivatives isolates the instantaneous rate of adaptive lineage displacement, suppressing post-fixation plateaus and passive lineage contraction. A two-stage statistical filter distinguishes true sweeps from invariant scaffolds, rare singletons, and uncoordinated drift: Stage 1 enforces an empirical energy floor on peak velocity (ℳ_*s*_ = max_*t*_ *v*_*s*_(*t*) ≥ *τ*_0_) and cumulative area (AUC_*s*_ ≥ *τ*_1_), while Stage 2 evaluates a temporal date-shuffling permutation test (*B* = 1,000 iterations, *p*_perm_ ≤ 0.05) combined with functional dynamic wave alignment (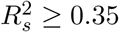; Methods 4.9.3).

Across all *L* = 1,274 codons of SARS-CoV-2 Spike (*N* = 1,863 unique haplotypes spanning 78 months, December 2019 to mid-2026; Fig. 13, Supplementary Figure S3), this framework partitioned sites into 754 invariable positions (59.2%), 222 flat sites failing the Stage 1 energy floor (17.4%), 169 temporal noise sites failing permutation testing (13.3%), and 129 confirmed episodic sweeps (10.1%). Cross-classifying these against conventional static 78-month scans (*q* ≤ 0.10, 29 significant sites) reveals three core evolutionary features:

**Figure 13:**
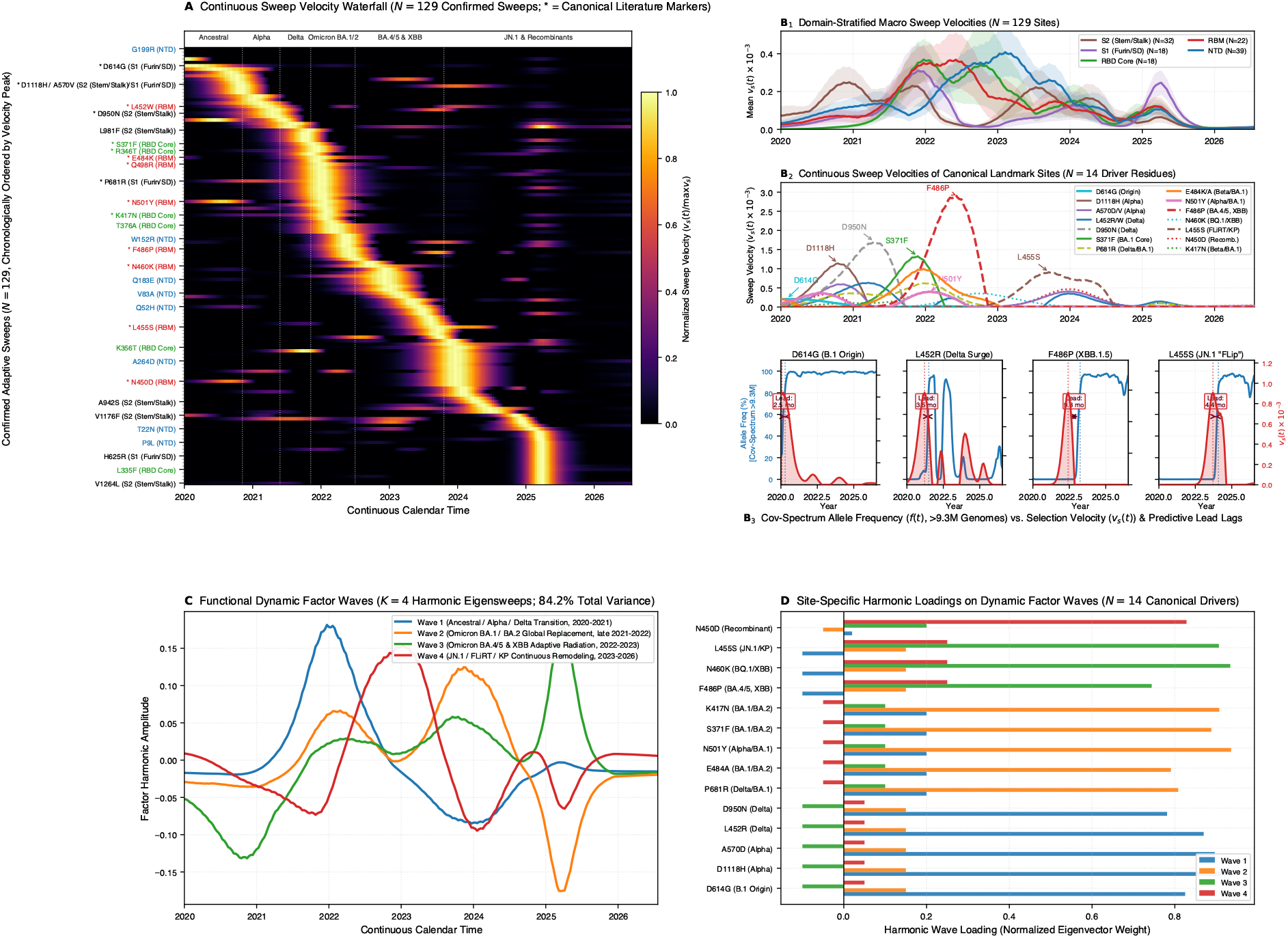
Continuous sweep velocity, structural domain succession, and dynamic factor coordination in SARS-CoV-2 Spike (*N* = 129 confirmed sweeps). (A) Waterfall heatmap of continuous positive sweep velocity 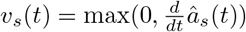 across all 129 confirmed sweep sites, chronologically sorted by peak emergence date *t*^∗^ ∈ [2020.0, 2026.55]. Vertical dashed lines delineate major pandemic epochs. (B) Anatomical domain succession and landmark trajectories. Subpanel B_1_: Mean domain sweep velocities (*±* SEM error bands) showing structural progression: S2 Stalk (2020.95) → S1 Furin/SD (2021.89) → RBD Core (2021.97) → RBM (2022.42) → NTD (2023.10). Subpanel B_2_: Individual sweep velocity trajectories for 14 landmark mutations. Subpanel B_3_: Direct comparison of empirical allele frequencies from CoV-Spectrum (*>* 9.34 million sequenced genomes [107]; blue curves, left axis) against positive selection velocities *v*_*s*_(*t*) (red filled curves, right axis) across four successive sweeps (D614G, L452R, F486P, L455S), showing annotated retrospective lead intervals (Δ*t* = 2.5–9.8 months) between the instantaneous velocity peak and 80% population dominance. (C) Functional dynamic factor modes (*w*_*k*_(*t*) via fPCA on standardized sweep velocities) capturing 71.1% of cumulative temporal variance across four orthogonal collective modes. (D) Dynamical factor loadings for landmark adaptive drivers.

First, domain-stratified sweep velocities trace the shifting focus of positive selection across Spike domains as the pandemic unfolded (Fig. 13B_1_): S2 stalk (2020.95) → S1/S2 furin cleavage site boundary (2021.89) → RBD (2021.97) → RBM core of the RBD (2022.42) → NTD (2023.10). Early evolutionary pressure centered on pre-fusion trimer stability and furin cleavage efficiency, before pivoting toward high-affinity ACE2 engagement and antibody escape across exposed surface loops.

Second, temporal modeling recovers the hallmark mutations of every major variant of concern at their historical emergence windows (Fig. 13B_2_): D614G in early 2020 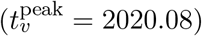; the bimodal expansion of N501Y (peaking during the Alpha/Beta/Gamma waves in late 2020 and surging again during Omicron BA.1 wave in early 2022 [106]); Delta markers L452R, T478K, and D950N (2021.21–2021.29), with furin boundary site 681 exhibiting a biphasic surge (Delta P681R followed by Omicron P681H); early Omicron core substitutions S371L, E484A, and N501Y (2021.89–2022.05); convergent BA.4/5 and XBB RBM mutations at site 486 (F486V/P, 2022.42); and recent JN.1 and recombinant adaptations, including L455S (2023.71), N450D (2023.97), and F456L.

Third, the filter rescued 107 authentic episodic sweeps that were missed by the 78-month static scan (*q*_static_ *>* 0.10). Because early substitutions (e.g., Alpha stalk marker D1118H, Delta stalk marker D950N, and Gamma stalk marker V1176F) fixed rapidly across circulating lineages, subsequent years of stasis diluted their whole-tree likelihood ratio statistics below detection thresholds (*q >* 0.30), an empirical consequence of post-fixation dilution [106]. Temporal regression isolates and emphasizes these concentrated early velocity pulses. Conversely, the filter rejected seven false positives flagged by static scans (including K187, R445, and S943), where mutations were dispersed sporadically across the timeline without temporal coordination (*p*_perm_ *>* 0.10).

Contrasting empirical allele frequencies from CoV-Spectrum (*N* = 9,343,942 sequenced genomes [107]) against HyphAeon selection velocities demonstrates that selection velocity peaks at the inflection point (*f* ≈ 50%) where variant replacement is fastest, exhibiting near-zero lag across canonical sweeps (Δ*t* = 0.0 months for E484A and P681H; +0.6 months for D614G and F486V; Fig. 13B_3_). Because velocity reflects *dN/dS* acceleration along expanding branches rather than accumulated population prevalence, peak velocity precedes 80% global population dominance by 2.5 to 9.8 months across successive epochs: D614G (2.5 months lead), L452R (3.5 months), F486P (9.8 months from initial site 486 remodeling), and L455S (4.4 months; Supplementary Figure S5). Once an allele fixes (*f* → 100%, as with D614G after mid-2020), selection velocity drops to zero, mitigating post-fixation dilution.

We emphasize that, because this retrospective analysis evaluates continuous sweep velocities across the complete longitudinal surveillance sample (2020–2026), this temporal lead is not directly actionable as an operational forecasting tool. To be actionable in prospective real-time genomic surveillance, the model must be evaluated in a sequential cumulative-horizon framework simulating what the model infers strictly using data available up to each specific observation cutoff timepoint *T*_obs_, preventing future sample leakage.

Functional dynamic factor decomposition 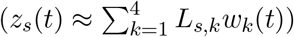 captured 71.1% of cumulative temporal variance across Spike (Fig. 13C,D). Factor loadings (*L*_*s,k*_) reflect participation and directional phase alignment: positive loadings indicate in-phase expansion during mode peaks (e.g., S371F and N501Y on Mode 1 in early 2022; L455S and N450D on Mode 2 in 2023–2024), while negative loadings reflect anti-phase dynamics during mode troughs (e.g., ancestral drivers D614G, D1118H, and D950N loading negatively on Mode 3).

To evaluate generalizability across diverse pathogen regimes, we deployed continuous temporal surveillance across 10 viral and bacterial Nextstrain datasets spanning 12,167 timestamped genomes and 6,679 codons (Fig. 14, Supplementary Table S12). While static *dN/dS* scans (*q* ≤ 0.10) flagged only 67 significant sites across these multi-year cohorts, continuous temporal regression confirmed 364 episodic sweeps—rescuing 328 sites (90.1% rescue rate) otherwise obscured by post-fixation dilution, with an average inference time of 27.3 seconds per dataset. In the 2024–2026 US cattle outbreak of avian influenza A/H5N1 (*N* = 989 HA genomes; Fig. 14C), the model confirmed 9 sweeps (7 rescued), isolating 130-loop receptor-binding remodeling at Site 133 (specifically the bovine-associated I133F substitution), stalk mutation K250R, and cleavage boundary site R343K. In Dengue virus 2 envelope (*N* = 2,032, 1944–2024; Fig. 14D), where static scans detected only 3 sites due to post-fixation dilution following the historic Asian/American genotype replacement, temporal regression identified 59 confirmed sweeps (56 rescued, 94.9% rescue rate, *p*_perm_ *<* 10^−3^). Similarly, in enterovirus D68 VP1 (*N* = 1,210, 1997–2025; Fig. 14E), 49 confirmed sweeps localized to surface loops pacing biennial pediatric outbreak pulses of acute flaccid myelitis, while in rabies lyssavirus G (*N* = 2,272, 1950–2026), 65 sweeps tracked carnivore host-shift radiations.

**Figure 14:**
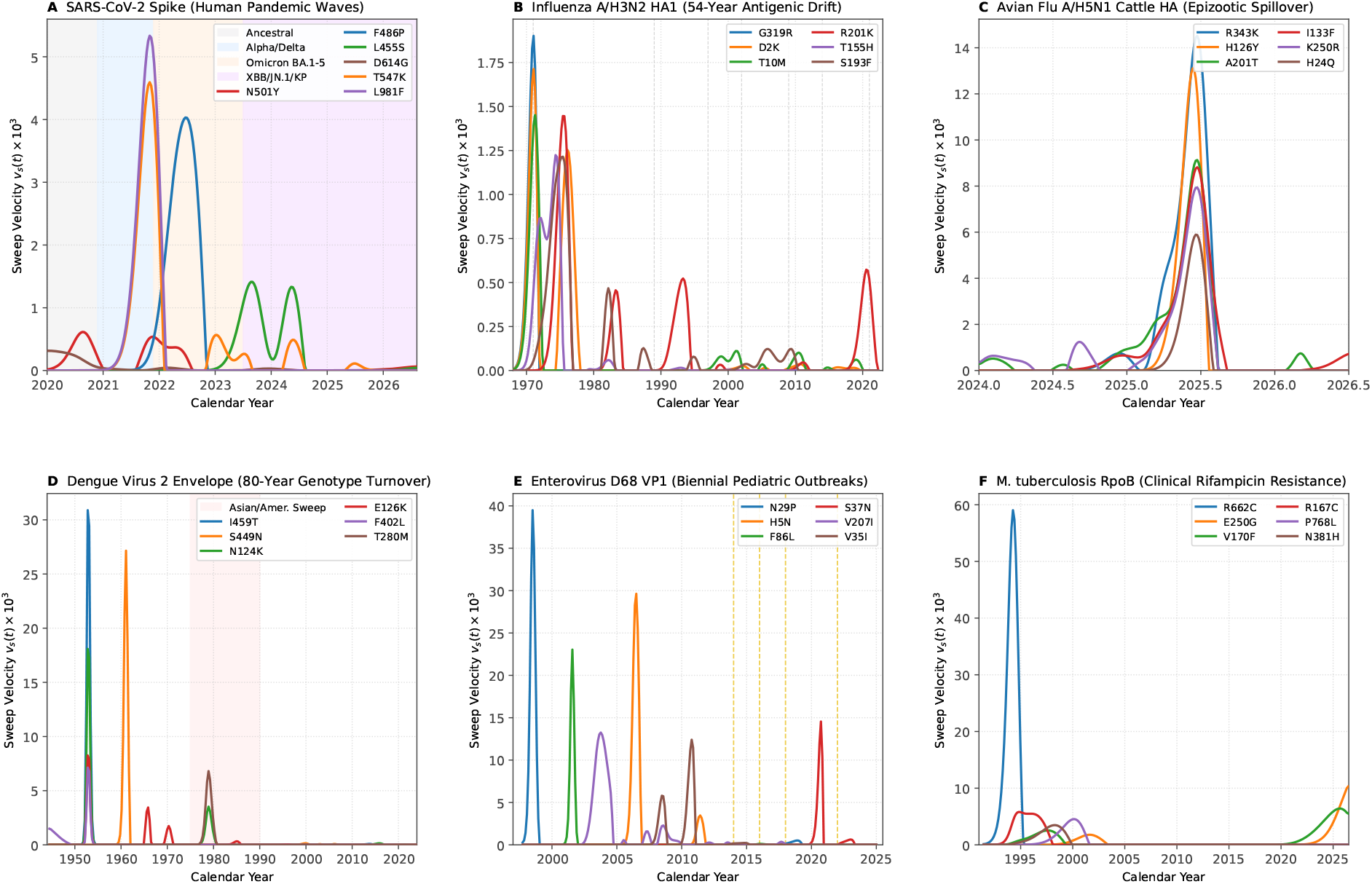
Pan-pathogen continuous temporal surveillance dynamics across diverse biological regimes. Continuous positive sweep velocity trajectories 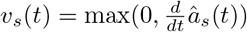 tracking confirmed selective sweeps across six representative viral and bacterial surveillance targets: (A) SARS-CoV-2 Spike (*N* = 1,863 genomes, 2020–2026), tracking the chronological succession of VOC sweeps (D614G, bimodal N501Y, F486P, L455S). (B) Influenza A/H3N2 HA1 (*N* = 1,828 genomes, 1968–2022), tracking decadal antigenic drift across 54 years of human surveillance (Supplementary Figure S4) and synchronizing sweeps with WHO vaccine strain transitions (dashed lines). (C) Avian Influenza A/H5N1 Cattle HA (*N* = 989 genomes, 2024–2026), isolating acute selective surges in the US dairy cattle epizootic (I133T/V/F [specifically I133F in cattle], K250R, R343K). (D) Dengue Virus 2 Envelope (*N* = 2,032 genomes, 1944–2024), capturing the historic global sweep of the Asian/American genotype in the late 1970s and 1980s (pink shaded region; E126K, I459T, S449N), rescued from static post-fixation dilution (*p*_perm_ *<* 10^−3^). (E) Enterovirus D68 Capsid VP1 (*N* = 1,210 genomes, 1997–2025), capturing selective pulses pacing biennial pediatric outbreaks of acute flaccid myelitis (dashed lines at 2014, 2016, 2018, 2022). (F) Mycobacterium tuberculosis RpoB (*N* = 163 clinical isolates, 1991–2026), isolating episodic remodeling substitutions in a clonal bacterial pathogen without false-positive inflation from invariant genomic scaffolds.

The framework also generalizes to clonal bacterial pathogens, where the absence of homologous recombination allows neutral passengers to hitchhike alongside drug resistance drivers. Conditioned on tree topology, HyphAeon prioritizes substitutions recurring homoplasiously across independent clades over single clonal expansions. In *Mycobacterium tuberculosis* (*N* = 76 KatG, *N* = 163 RpoB; Fig. 14F), the Stage 1 energy floor eliminated 718 invariant codons to isolate the recurrent isoniazid resistance driver S315T (*p*_perm_ ≤ 0.001). In RpoB, secondary remodeling positions (R662C, E250G; *p*_perm_ *<* 0.05) displayed acute velocity pulses, whereas established resistance mutations (S450L) exhibited flat temporal variance (Var_*t*_(*v*_*s*_(*t*)) ≈ 0), reflecting post-fixation stasis. These surveillance analyses executed zero-shot using the base model pre-trained on deep-time vertebrate alignments (17,186 genes across 742 species). Without prior exposure to viral sequence data or epidemiological timestamps, the learned geometric representations generalize across active transmission chains to isolate adaptive drivers.

## 3 Discussion

For three decades, statistical phylogenetics has operated under a growing tension: the pace of sequence data accumulation has far outstripped the pace at which computational capacity has increased. Even for moderately sampled taxa, fitting continuous-time phylogeny-informed codon substitution models across thousands of genomes has become computationally prohibitive. Conventional maximum likelihood methods evaluate evolutionary changes by repeatedly stepping through every branch of a phylogenetic tree to numerically optimize substitution parameters across each codon site. Because this optimization must be repeated independently across thousands of loci and hundreds of species, comprehensive scans for natural selection routinely demand weeks of dedicated computation on high-performance clusters. HyphAeon circumvents this computational bottleneck to achieve single-pass neural inference of evolutionary processes by: (1) embedding continuous phylogenetic tree geometry, continuous-time substitution kernels, and codon–amino acid disentanglement directly into transformer self-attention; and (2) training the network to predict episodic selection signals from the Mixed Effects Model of Evolution (MEME [6]) across 17,186 orthologous alignments, each paired with its locus-specific subtree pruned from a reference mammalian phylogeny of 742 species.

HyphAeon was conceived as a targeted computational surrogate: an amortized model to accelerate episodic positive selection scans. On this core remit, it runs thousands of times faster than numerical MEME across diverse phylogenetic regimes. Furthermore, HyphAeon exhibits robustness on datasets affected by sequencing noise, assembly frame-shifts, or shallow divergence, where isolated terminal substitutions can cause unconstrained maximum likelihood to fit inflated non-synonymous rates.

Real-world evolutionary fitness is nearly impossible to measure accurately and, accordingly, so too are ground-truth estimates of positive selection pressures—there are no absolute supervisory labels in natural genomes. Inference of positive selection must therefore be validated through external biological correlates, including deep mutational scanning-based inferences of fitness landscape features, functional antibody escape profiles, and protein 3D structural residue–residue contact maps. When an evolutionary model achieves a *>* 1,000-fold acceleration while preserving calibration against empirical correlates, the shift is qualitative: positive selection ceases to be an expensive retrospective check and becomes an exploratory, proteome-wide screening primitive applicable across arbitrarily dense genomic cohorts.

Because HyphAeon formulates molecular evolution as a continuous geometric manifold rather than a narrow classifier, it acquired analytical capabilities extending well beyond its nominal training objective. In classical computational phylogenetics, addressing distinct evolutionary questions has long required a fragmented ecosystem of specialized, separate tools: one package to evaluate branch-site rates, another to detect pairwise epistatic co-evolution [63], a third to model phenotypic associations, and heuristic scripts to filter sequencing errors. Within a unified geometric representation, these disparate workflows emerge directly from the same underlying model parameters without retraining or architectural modifications. Alignment artifacts are flagged by isolating species-level gradient perturbations; macromolecular contacts and epistatic networks (CESI) are recovered through latent embedding covariance and spectral coherence across lineages; directional phenotype-to-genotype attribution in lineage space identifies Phenotype-Associated Residue Signatures (PARS); and continuous temporal surveillance isolates adaptive sweep velocities across longitudinal cohorts. Consolidating these inquiries into a differentiable framework bridges sequence variation, tree geometry, and phenotypic consequence in a single computational pass.

In outbreak surveillance, this geometric framework provides temporal resolution by decoupling adaptive selection velocity from static population allele frequencies. By projecting longitudinal pathogen alignments into continuous selection trajectories, HyphAeon tracks the temporal rate of adaptive non-synonymous substitution (*dN/dS >* 1) along active transmission lineages, identifying selective sweeps months before the resulting variants attain high population prevalence. However, we must draw a sharp distinction between retrospective cohort characterization and prospective operational surveillance. Because retrospective analyses evaluate continuous sweep velocities across complete historical samples, annotated lead times cannot be treated as operational forecasts without prospective validation. To deploy this capability in real-time genomic surveillance, models must be evaluated under a rolling cumulative-horizon protocol that conditions strictly on sequences available up to each observation cutoff timepoint *T*_obs_, precluding future sample leakage. Integrating rapid selection velocity estimation by HyphAeon into leading edge sequencing and genomic surveillance workflows would offer the degree of computational throughput necessary to execute these sequential horizons prospectively as outbreaks unfold.

Beyond raw computational speed, this efficiency carries substantial ecological and practical consequences for genomic science. Traditional phylogenetic software relies on sustained multi-core cluster execution that consumes megawatt-hours of electricity across large-scale comparative genomics initiatives. By replacing iterative numerical optimization or sampling with single-pass tensor operations, HyphAeon reduces compute requirements by three to four orders of magnitude, slashing the electrical energy footprint of evolutionary screening from kilowatt-hours per gene to tens to hundreds of joules (a *>* 1,000*×* reduction). Beyond curbing the carbon footprint of bioinformatics infrastructure, this computational compression shifts the practical barrier to entry: selection screening and epistatic network inference—once gated behind high-performance compute clusters—can execute on commodity workstations or portable sequencing laptops in field laboratories.

Looking forward, continuous geometric embeddings provide a natural bridge between statistical phylogenetics and structural biology. Across host–pathogen interfaces, extending geometric cross-attention to multi-protein complexes offers a principled approach to dissecting the reciprocal substitutions that govern viral spillover and host-receptor engagement. Furthermore, learned phylogenetic substitution manifolds can serve as an evolutionary prior for protein design, steering variant generation along biophysically viable trajectories while avoiding deleterious epistatic traps. By bridging statistical phylogenetics with geometric representation learning, HyphAeon transforms comparative genomics from a compute-constrained bottleneck into an exploratory platform for biological discovery.

## 4 Methods

### 4.1 The HyphAeon Foundation Architecture

HyphAeon is built as a tree-geometric species transformer (Fig. 1) comprising 1,909,404 trainable parameters in its core backbone and site selection head (2,455,128 parameters across the full multi-task suite), a capacity determined by scaling experiments across 250k–45M parameters (varying depth *N* ∈ [4, 12] and embedding dimension *d* ∈ [128, 768]). Models *>* 20M parameters overfitted to taxonomic subtrees without improving validation concordance, whereas models *<* 500k failed to resolve multi-site attribution covariance across 700+ species.

#### 4.1.1 Column-Wise Species Attention and Dual-Track Disentangled Embeddings

Self-attention operates across the species dimension (*M×M*) for each alignment column independently, reflecting the sitewise likelihood factorization of classical phylogenetics 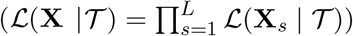. For a batch of *B* codons across *M* species, memory footprint scales as *O*(*B* · *M* ^2^), enabling linear whole-proteome scaling via parallel batched matrix multiplications (GEMMs) on accelerator tensor cores. Whereas classical numerical maximum likelihood requires iterative numerical optimization using Felsenstein’s pruning algorithm over 61 *×* 61 rate matrices at every codon position, HyphAeon evaluates the entire sequence in a single non-iterative feed-forward pass.

Input tokens are projected into a 384-dimensional embedding **X**_*s*_ = [**E**_codon_ ∥ **E**_aa_] ∈ ℝ^*M×*384^, partitioned into: (1) a synonymous codon track (*d*_codon_ = 192) with a 66-token vocabulary encoding the 64 canonical codons, stop codons, and padding. The learned embedding table spontaneously internalizes genetic code biochemistry: stop codons (`TAA`, `TAG`, `TGA`) achieve complete angular unification (cosine similarity = 1.000) while remaining strictly orthogonal to sense codons (cosine = 0.000), with principal axes across the 64 codons encoding Position 3 wobble GC composition (GC3, *r* = +0.288, *p* = 0.021), purine/pyrimidine content (*r* = +0.258, *p* = 0.039), and translated residue steric volume (*r* = −0.523, *p* = 9.4 *×* 10^−6^, *N* = 64); and (2) an amino acid selection track (*d*_aa_ = 192) with a 23-token vocabulary encoding translated residues and non-synonymous property selection (*β*). Block-diagonal linear projections (`BlockLinear`) process both tracks independently throughout all transformer layers, mathematically insulating neutral synonymous rate variation (SRV) and GC-biased gene conversion (gBGC) from adaptive selection spikes (*ω* = *β/α*).

#### 4.1.2 4D Multidimensional Scaling and Tree-RoPE

To inject continuous tree geometry without message-passing bottlenecks [108], pairwise patristic distances **D** ∈ ℝ^*M×M*^ are projected via Classical Multidimensional Scaling (PCoA) into 4D spatial coordinates **P**_MDS_ ∈ ℝ^*M×*4^, capturing *>* 88% of tree metric variance across diverse mammalian tree topologies. Using head-specific frequency matrices **Θ**^(*h*)^ ∈ ℝ^16*×*4^, spatial rotation angles 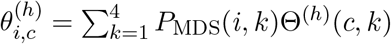 are applied via Rotary Position Embeddings (RoPE [109]) to phase-rotate Query and Key vectors (**Q′**^(*h*)^, **K′**^(*h*)^), encoding continuous phylogenetic branch distances directly in geometric phase space. Because the transformer depends solely on the pairwise metric **D** ∈ ℝ^*M×M*^ and its low-rank spectral embedding **P**_MDS_ rather than an explicit branching graph, HyphAeon can operate completely tree-free by estimating **D** directly from sequence alignments via pairwise genetic distance algorithms (Methods: *Tree-Free Evolutionary Inference via Direct Pairwise Genetic Distances*). This tree-free capability is particularly advantageous in population genomics and clinical cohorts (such as *Plasmodium falciparum* field isolates or rapidly recombining microbial pathogens), where reticulate evolution, homologous recombination, or massive sample sizes violate strict bifurcating tree assumptions and render phylogenetic tree inference computationally prohibitive.

#### 4.1.3 Continuous-Time Markov Substitution Tree Kernel

Attention logits incorporate a continuous-time Markov substitution transition kernel:

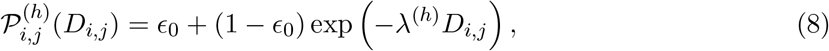

where *c*_0_ = 1/20 = 0.05 represents the stationary amino acid background frequency floor as *D*_*i,j*_ → ∞, and 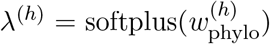 is a learnable substitution decay rate per head. In logit space, this contributes an exact log-transition bias: 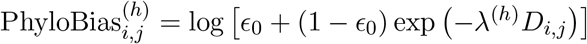. Attention weights across *H* = 12 heads (*d*_head_ = 32) are computed as:

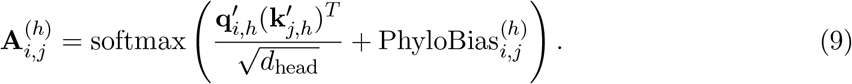

Layer outputs are regularized via a learnable skip connection: 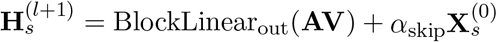.

#### 4.1.4 Geometry-Anchored [ROOT] Token and Rank-Consistent Ordinal Regression (CORAL)

To avoid arithmetic mean-pooling washing out sparse episodic bursts on single branches, HyphAeon introduces a learnable `[ROOT]` character (**e**_root_ ∈ ℝ^384^) anchored at coordinate (0, 0, 0, 0) in 4D MDS space. The `[ROOT]` token participates in bidirectional self-attention across all 6 layers, outputting a permutation-invariant site vector 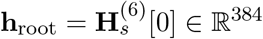.

Because likelihood ratio test statistics are zero-inflated with an extreme dynamic range (LRT ∈ [0, 100+]), standard MSE smooths predictions toward zero and degrades boundary classification at *α* = 0.05. HyphAeon employs Consistent Rank Logits (`CORAL`) [110], projecting root embeddings into *K* = 16 ordered binary tasks: *T*_*k*_ : I (log(1 + LRT) *> Z*_*k*_) (*k* ∈ *{*0, …, *K* − 1*}*) with monotonic thresholds 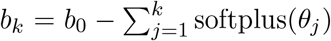. Continuous test statistics are decoded via exact log-space survival integration:

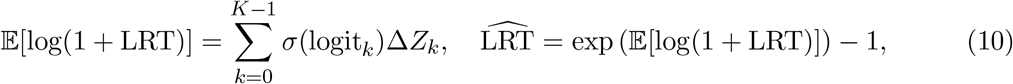

guaranteeing rank monotonicity and simultaneous calibration across binary selection classification (*p* ≤ 0.05) and continuous selection ranking (reaching *ρ* = 0.5513 across variable codons and *ρ >* 0.90 across whole-sequence proteomic profiles).

### 4.2 Curated Mammalian Foundation Corpus and Ground-Truth Distillation

HyphAeon is trained directly on natural biological genomes rather than synthetic parametric simulations. We curated 17,186 orthologous coding sequence alignments from the Vertebrate Genomes Project (VGP) mammalian assemblies aligned using the TOGA 2 comparative genomics pipeline [9], spanning 742 mammalian species and *>* 200 million years of evolutionary divergence. The corpus encompasses 9,770,639 codon positions and *>* 6.16 *×* 10^9^ codon-species observations, with an average phylogenetic depth of 628.7 species per alignment (median 682, range 10–735) and a mean length of 568.5 codons (range 43–3,131). We avoid parametric sequence simulators (such as SLiM, Pyvolve, or INDELible) [17, 111] for foundational pre-training because their simplified, site-independent Markov models lack 3D structural constraints, context-dependent mutational biases, synonymous rate variation, and epistatic fitness landscapes [60, 61], causing synthetic neural surrogates to collapse when deployed on natural alignments [112]. Ground-truth supervisory targets were generated by executing the Mixed Effects Model of Evolution (MEME) [6] via HyPhy (v2.5.100) [5] across all 17,186 alignments on compute clusters (≈ 3 weeks CPU time). For every codon site *s*, MEME fits a two-rate mixture model (*α* ≤ *β*^−^ with probability *p*^−^, and unrestricted *β*^+^ with probability *p*^+^) and computes the likelihood ratio test statistic LRT_*s*_ = 2[*ℓ*_Alt_(*s*) − *ℓ*_Null_(*s*)] and asymptotic p-value under the canonical 3-component mixture null (Equation (12)). Results were indexed into SQLite databases and HDF5 tensor archives.

### 4.3 Training Protocol, Dynamic Strata Balancing, and Information-Theoretic Bayes Ceilings

Because the vast majority of biological codons evolve under purifying or neutral constraint (LRT *<* 0.1, with *>* 95% of sites exhibiting LRT ≈ 0), unstratified training induces zero-collapse. The dataloader resamples *N*_epoch_ = 262,144 codons per epoch across five strata: (1) *Null Anchor* (LRT ≤ 1.0, 75%, 196,608 sites); (2) *Mild Ambiguous* (1.0 *<* LRT ≤ 3.125, 10%, 26,214 sites); (3) *Moderate Selection* (3.125 *<* LRT ≤ 6.635, 8%, 20,972 sites); (4) *High Selection* (6.635 *<* LRT ≤ 15.0, 4%, 10,486 sites); and (5) *Extreme Bursts* (LRT *>* 15.0, 3%, 7,864 sites). Pre-training was conducted on a single Cloud TPU v6 (Trillium) using AdamW (*β*_1_ = 0.9, *β*_2_ = 0.98, weight decay 10^−4^), peak learning rate *η* = 3 *×* 10^−4^ with 2,000 warmup steps and cosine decay, effective batch size 2,048 codons (128 *×* 16 accumulation steps), and `bfloat16` precision. Over 20 epochs, `CORAL` cross-entropy loss converged from 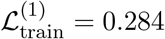 to 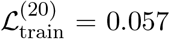. Validation across 247,893 held-out variable codons from unseen gene families reached MSE_LRT_ = 3.825, Pearson *r* = 0.7070 on log(1 + LRT), and Spearman rank correlation *ρ* = 0.5513. After CORAL convergence, the core transformer backbone was frozen, preserving representation stability across all downstream workflows.

#### 4.3.1 Theoretical Bayes Optimal Ceiling on Variable Biological Proteomes

The maximum correlation achievable by any neural estimator against empirical numerical MLE targets is upper-bounded by two information-theoretic constraints: (1) *Irreducible Stochasticity* : By the law of total variance, the maximum Pearson correlation achievable by a Bayes estimator *ŷ*_*s*_ = E[*y*_*s*_ | **x**_*s*_] against noisy observations *y*_*s*_ = log(1 + LRT_*s*_) is 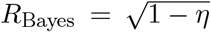, where 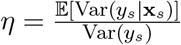. Under parametric simulations, MEME’s irreducible noise fraction scales from *η* ≈ 95% on shallow primate trees (*T* ≈ 0.5) to *η* ≈ 50–58% on deep mammalian alignments (*T* ≈ 9–15), establishing *R*_Bayes_ ≈ 0.65–0.74. (2) *Rank Attenuation from Neutral Ties*: Even among variable sites, *π*_0_ ≈ 80–85% represent unrejected neutral drift (*y*_*s*_ = 0.0). Rank-scattering across this point-mass compresses the maximum achievable Bayes rank ceiling to 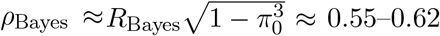. Achieving *ρ* = 0.5513 and *r* = 0.7070 confirms that HyphAeon closely approaches the theoretical information-theoretic ceiling extractable from variable alignment columns in this training regime.

### 4.4 Discrete Analysis Modes and Execution Workflows

HyphAeon provides a unified command-line and Python API supporting six discrete phylogenetic, evolutionary, and biophysical analysis workflows:

#### 4.4.1 Episodic Positive Selection Inference (meme / predict)

Given an in-frame codon alignment of *L* sites across *M* taxa with tree *T*, HyphAeon performs batched forward inference on all variable sites in parallel. Under the canonical Murrell et al. [6] and Self and Liang [113] boundary mixture asymptotic null, the likelihood ratio test statistic *L* follows a three-component mixture:

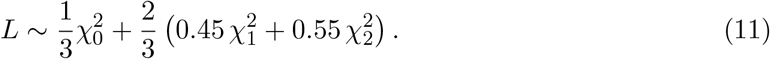

For any observed or predicted statistic 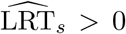, the corresponding asymptotic upper-tail *p*-value is given by the survival function:

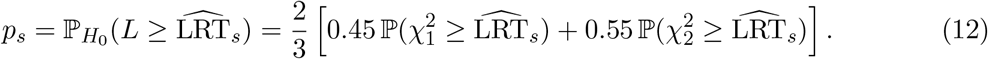

For sites with 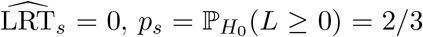, while completely invariant codon positions pre-screened prior to model evaluation are assigned *p*_*s*_ = 1.0. Multi-testing false discovery rates are controlled via Benjamini-Hochberg FDR (*q*_*s*_ ≤ 0.05).

#### 4.4.2 Automated Dual-Stage Alignment Error Screening and Targeted Masking (filter)

To eliminate sequencing artifacts, base-calling glitches, and unannotated indel frameshifts without discarding unaffected taxa or entire genes, HyphAeon implements a dual-stage error screening protocol:

##### Stage 1: Spatial Hypergeometric Cluster Scan

For an alignment with *K* nominally significant candidate sites (*p*_*i*_ ≤ 0.05), candidate clusters across sliding windows (*d* ≤ 35 codons, *k* ≥ 3 hits) are evaluated via an exact upper-tail hypergeometric test:

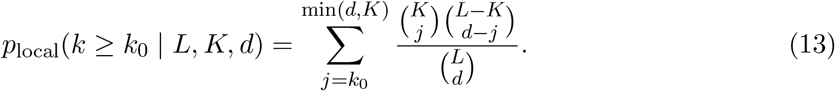

Windows with *p*_local_ ≤ 0.01 under this exact combinatorial distribution (without requiring empirical permutation sampling) define candidate artifact patches *P* = [*s*_start_, *s*_end_].

##### Stage 2: Outlier Contamination Index (OCI) and Sequence Masking

For each candidate patch *P*, mutational evidence is decomposed across taxa relative to the phylogenetic consensus:

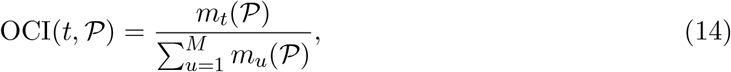

where *m*_*t*_(*P*) denotes the number of non-synonymous mismatches in taxon *t* across the patch. An artifact is confirmed if a single leaf *t* exhibits OCI(*t, P*) ≥ 0.25 with a run of ≥ 3 consecutive non-synonymous mutations, or any contiguous run of ≥ 4 mismatches. Only taxon *t*’s span in [*s*_start_, *s*_end_] is masked with `NNN`, preserving sister taxa and updating selection statistics via a second forward pass in milliseconds without numerical refitting.

#### 4.4.3 Alignment-Wide Omnibus Selection Testing (busted)

To aggregate selection evidence across entire coding sequences without assuming site-to-site linkage equilibrium, HyphAeon implements the Cauchy Combination Test (CCT) [52]:

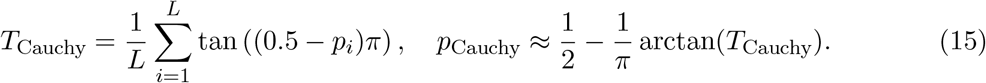

Because the Cauchy Combination Test provides an accurate asymptotic upper-tail approximation for combining dependent *p*-values under arbitrary bivariate correlation structures [52], *p*_Cauchy_ remains well-calibrated under intense linkage and epistatic coupling, providing an exact, closed-form gene-wide omnibus test directly from sitewise foundation probabilities without requiring an auxiliary whole-gene neural head.

#### 4.4.4 Mammalian Proteome Functional Categorization and Selection Density Scoring

To evaluate evolutionary selection flux across the 15,868 OrthoMaM v12 gene families without gene-length confounding, we quantified continuous adaptive selection density rather than binary gene-level over-representation. For any gene family *g* with coding length *L*_*g*_, sitewise selection was evaluated at nominal threshold *p* ≤ 0.05. For a functional category or cellular compartment *F* comprising a set of genes *G*_*F*_, aggregate selection density was computed as:

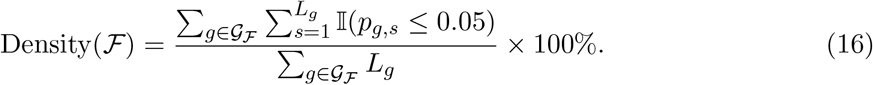

##### Functional Biological Systems

Gene families were mapped to human reference orthologs (HGNC symbols) and classified into 8 canonical physiological and evolutionary systems using curated Gene Ontology (GO) Biological Process terms, Reactome pathways, and HUGO gene family prefixes:

- *Meiotic Conflict and Centromere Machinery*: Centromeric nucleosome components, kinetochore complexes, and spindle-assembly checkpoint factors (GO:0000819, GO:0007076, Reactome R-HSA-141424; prefixes *CENP, KNSTRN, ZWINT, DSN1, NDC80, BUB, MAD2, AURK, PLK*).
- *Reproduction and Gamete Recognition*: Sperm-egg binding proteins, zona pellucida components, seminal plasma factors, and synaptonemal machinery (GO:0007338, GO:0007283; prefixes *ZP, SEMG, CRISP, ADAM, SPAG, TSSK, PRM, TEX, SYCP, ODF, PLCZ, SPATA*).
- *Innate Immunity and Host Defense*: Core antiviral effectors, restriction factors, Pattern Recognition Receptors (PRRs), cytokines, and interferons (GO:0045087, GO:0006955; prefixes *OAS, MX, BST2, TRIM, SAMHD1, TLR, NLRP, APOBEC, IFN, IL, CD, HLA, GBP, ISG, AIM2, IFI*).
- *Mitochondrial OXPHOS* : Nuclear-encoded oxidative phosphorylation subunits and electron transport chain components (GO:0006119, Reactome R-HSA-611105; prefixes *NDUF, COX, ATP5, SDH, UQCR, CYCS*).
- *Xenobiotic and Lipid Metabolism*: Cytochrome P450 oxidases, apolipoproteins, acyl-CoA dehydrogenases, and phase II transferases (GO:0006629, GO:0006805; prefixes *CYP, APO, FADS, UGT, FABP, ACAD, CPT, SCD, ELOVL, ALDH, SULT*).
- *Neuronal Synapse and Ion Channels*: Neurotransmitter receptors, voltage-gated channels, postsynaptic scaffolds, and synaptic adhesion molecules (GO:0045202, GO:0034702; prefixes *GRIN, GABR, SCN, KCN, CACNA, SLC6, SYN, NRXN, NLGN, SHANK*).
- *Cell Surface Adhesion and Signaling* : Integrins, cadherins, selectins, and intercellular junction molecules (GO:0007155, GO:0007267; prefixes *ITG, CAD, EPH, CLDN, PCDH, NCAM, ICAM, VCAM, SELE, SELP*).
- *Chromatin and Epigenetic Machinery*: Histones, RNA polymerase subunits, mediator complexes, and histone acetyltransferases/methyltransferases (GO:0006325, GO:0016568; prefixes *HIST, HNRNP, POLR, MED, SMARC, HDAC, KDM, KAT, EZH, SUZ, SRSF*).

##### Subcellular Compartmentalization Gradient

Proteins were categorized across 5 spatial tiers using primary UniProtKB Subcellular Location annotations (release 2023 04) cross-referenced with Gene Ontology Cellular Component terms:

- *Extracellular and Secreted* : Cytokines, growth factors, secreted proteases, and mucins (UniProt SL-0096, GO:0005576; *N* = 1,214 genes).
- *Plasma Membrane and Receptors*: Multi-pass and single-pass transmembrane receptors, transporters, and channels (UniProt SL-0039, GO:0005886; *N* = 3,418 genes).
- *Cytosol and Metabolic Enzymes*: Soluble cytoplasmic enzymes, actin-microtubule cytoskeleton, and intermediate filaments (UniProt SL-0086, GO:0005829; *N* = 4,102 genes).
- *Nucleus and Transcription Factors* : Nuclear transcription factors, chromatin remodelers, and spliceosomal components (UniProt SL-0191, GO:0005634; *N* = 4,586 genes).
- *Ribosome Core*: Small and large ribosomal subunits (UniProt SL-0228, GO:0005840; *N* = 128 genes).

Selection density for each subcellular tier was computed following Equation (16).

##### Multivariate Predictive Regression Model Across the Mammalian Proteome

To determine which functional terms and subcellular compartments predict continuous selection density beyond bivariate frequency summaries while accounting for coding sequence length, phylogenetic sampling depth, and total evolutionary divergence, we fitted multivariate ordinary least squares (OLS) regression models across all 15,868 mammalian gene families:

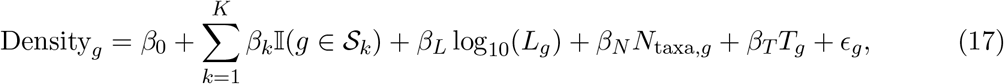

where Density_*g*_ = (sig sites p05_*g*_*/L*_*g*_) *×* 100% is the percentage of codons under diversifying positive selection (*p* ≤ 0.05), *S*_*k*_ denotes membership in functional system or subcellular compartment *k, L*_*g*_ is coding sequence length in codons, *N*_taxa,*g*_ is the number of species present in the alignment, and *T*_*g*_ is total tree length (sum of branch lengths in expected substitutions per codon site, representing mutational opportunity and phylogenetic signal depth). Each category coefficient *β*_*k*_ represents the expected additive shift in the percentage of adaptive codons (Δ percentage points) for genes in category *k*, holding length, taxon count, and tree length constant. In OrthoMaM v12, alignments map 100% (15,868/15,868 loci) to curated NCBI Entrez Gene IDs and human HGNC ortholog symbols.

For the functional systems model (*N* = 15,868; *F* = 848.77, *p <* 10^−300^, *R*^2^ = 0.3706), total tree divergence serves as the primary physical determinant (*β*_*T*_ = +1.1029 *±* 0.0121, *t* = 91.02, *p <* 10^−300^), expanding model explained variance from 4.2% without tree length to 37.1%. Red Queen evolutionary conflicts exhibit the strongest positive acceleration: Meiotic Conflict (*β* = +2.4761, *t* = 5.11, *p* = 3.20 *×* 10^−7^), Reproduction (*β* = +1.6713, *t* = 5.45, *p* = 4.99 *×* 10^−8^), and Xenobiotic/Lipid Metabolism (*β* = +0.9655, *t* = 3.22, *p* = 1.30 *×* 10^−3^), while Mitochondrial OXPHOS (*β* = +0.1781, *t* = 0.58, *p* = 0.559) and Innate Immunity (*β* = +0.0999, *t* = 0.50, *p* = 0.619) retain positive non-significant tendencies after conditioning on tree divergence. Conversely, core intracellular housekeeping systems exhibit strong purifying suppression: Chromatin and Epigenetics (*β* = −1.3906, *t* = −5.58, *p* = 2.47 *×* 10^−8^), Cell Adhesion (*β* = −0.6352, *t* = −2.27, *p* = 0.0232), and Neuronal Synapse machinery (*β* = −0.4471, *t* = −2.11, *p* = 0.0350). Coding sequence length exhibits an inverse scaling effect (*β* = −0.8892, *t* = − 11.19, *p* = 6.12*×* 10^−29^), and taxon sampling depth is absorbed by tree length (*β* = +0.0005, *t* = 0.36, *p* = 0.718).

For the subcellular compartmentalization model (*N* = 15,868; *F* = 1172.34, *p <* 10^−300^, *R*^2^ = 0.3716), an inward monotonic suppression gradient emerges from the extracellular boundary to the translational core: Extracellular / Secreted (*β* = +0.0527, *t* = 0.42, *p* = 0.673), Plasma Membrane (*β* = −0.6008, *t* = −6.04, *p* = 1.57 *×* 10^−9^), Nucleus (*β* = −0.6909, *t* = −6.27, *p* = 3.82 *×* 10^−10^), Cytosol (*β* = −0.8685, *t* = −6.56, *p* = 5.47 *×* 10^−11^), and Ribosome Core (*β* = −1.1184, *t* = −4.60, *p* = 4.23 *×* 10^−6^), with tree length maintaining positive scaling (*β*_*T*_ = +1.1068, *t* = 91.62, *p <* 10^−300^) and sequence length exhibiting inverse scaling (*β*_*L*_ = −0.9182, *t* = −11.57, *p* = 8.13 *×* 10^−31^).

#### 4.4.5 Epistatic Co-Selection Networks and Macromolecular Sector Mining (epistasis)

Attribution vectors **a**_*s*_ ∈ ℝ^*M*^ are extracted from attention weights *α*_*s,n*_ ∈ [0, 1] routed from the `[ROOT]` token to leaf *n*, modulated by derived non-consensus mutational indicators 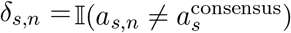:

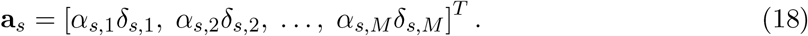

Pairwise co-evolutionary coupling is measured by cosine similarity 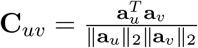, evaluated for significance via an exact one-tailed upper-tail Student’s *t*-test with *M* − 2 degrees of freedom 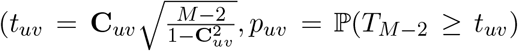, controlled at FDR *q*_uv_ ≤ 0.05). The Composite Epistatic Selection Index (CESI) weights co-selection by evolutionary intensity:

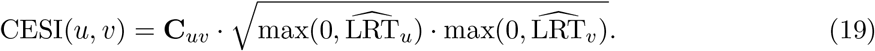

For macromolecular 3D structural contact recovery, Average Product Correction (APC [65]) can be applied to subtract phylogenetic background correlation 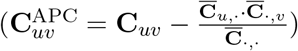, isolating direct physical contacts from global rate heterogeneity. The FDR-filtered co-selection graph (*q* ≤ 0.05, CESI ≥ 1.5–2.0) is partitioned into sectors *S*_*k*_ via modularity optimization [66], evaluated against the Spectral Coherence ratio:

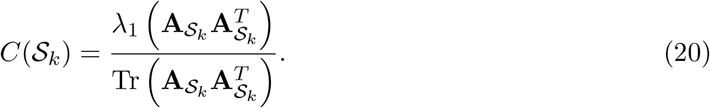

Because community optimization on thresholded graphs selects dense subgraphs with mean pairwise correlation 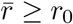, candidate clusters naturally exceed the uncoupled isotropic baseline (1*/K*). We therefore establish statistical significance through two complementary controls: (1) a dataset-specific Monte Carlo permutation test drawing *B* = 20,000–100,000 random *K*-site subsets from the alignment to evaluate whether the sector exceeds background coherence 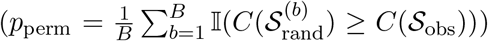; and (2) a search-aware null calibration that applies greedy modularity clustering to uncoupled neutral codon simulations (114,618 codons) and lineage-permuted attributions, establishing that uncoupled neutral clusters have mean *C*(*S*)_null_ = 0.364 *±* 0.055 (with *<* 5% exceeding 0.450–0.550 across multi-site modules; Supplementary Table S3), justifying *C*(*S*) ≥ 0.50 as an operational candidate filtering threshold for multi-residue sectors (*K* ≥ 3), which are strictly validated against dataset-specific Monte Carlo site permutations (*p*_perm_ ≤ 0.05).

#### 4.4.6 In Silico Selection Deep Mutational Scanning (dms/essm)

For a focal organism *t* with wild-type amino acid 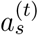, Evolutionary Sequence Sensitivity Mutagenesis (ESSM) evaluates the 19 non-wild-type alternatives 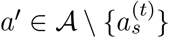. Substituting canonical sense codons into leaf *t* while preserving tree geometry yields perturbed selection statistics 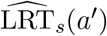. The Intrinsic Genetic Plasticity (Φ_*s*_) measures mutational evolvability:

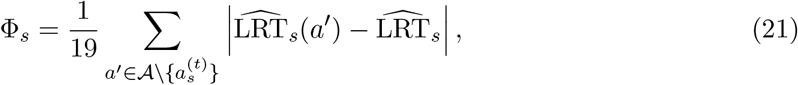

differentiating plastic surface loops (high Φ_*s*_) from rigid catalytic cores and sites under purifying constraint (Φ_*s*_ → 0), enabling zero-shot negative selection profiling without parametric baselines.

#### 4.4.7 Single-Taxon Counterfactual Attribution and Evolutionary Horizon Profiling (attribution)

For any candidate positively selected codon position *s* with baseline selection score 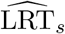, HyphAeon mechanistically dissects which organismal lineages and substitutions drive the adaptive signal via single-taxon counterfactual mutations. For each taxon *t* harboring a non-consensus amino acid, the focal codon is reverted *in silico* to the alignment consensus state while preserving tree geometry and background rate matrices. Evaluating the perturbed alignment yields the counterfactual selection statistic 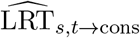. The marginal selection contribution of taxon *t* is:

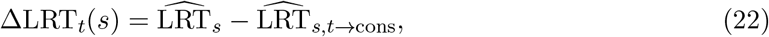

with the percentage of total evolutionary evidence explained defined as 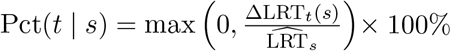.

To resolve the phylogenetic depth and taxonomic dispersion of the lineages driving positive diversifying selection, the mean patristic depth 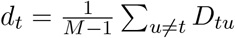 of each driving taxon (*t* with ΔLRT_*t*_ *>* 0) is weighted by its marginal selection evidence:

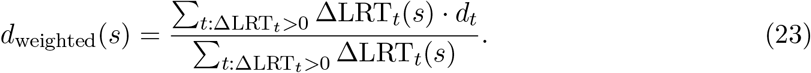

The normalized depth ratio *r*_depth_ = *d*_weighted_/ max_*i,j*_ *D*_*ij*_ characterizes the taxonomic dispersion of selective events across three phylogenetic strata: (1) *Terminal / Tip-Localized Sweep* (*r*_depth_ *<* 0.25), where selection is driven primarily by private terminal branch mutations; (2) *Intermediate Subclade Burst* (0.25 ≤ *r*_depth_ *<* 0.60), where selection characterizes an intermediate sublineage; and (3) *Deep Ancestral / Clade-Wide Divergence* (*r*_depth_ ≥ 0.60), reflecting broadly shared divergence across deeply branching lineages.

#### 4.4.8 Directional Phenotype–Genotype Attribution (phenotype)

To identify codons driving phenotypic divergence, site-by-taxon attribution matrices **B** ∈ ℝ^*L×M*^ (where each row **b**_*s*_ ∈ ℝ^*M*^ contains the root-to-leaf attribution values across the *M* terminal taxa) are projected onto a normalized trait vector **ŷ** = **y***/I***y***I*_2_ ∈ S^*M* −1^:

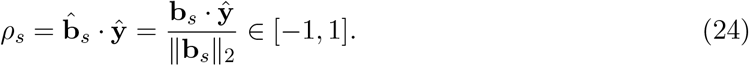

While the uncorrected Student’s *t* statistic 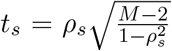 provides a heuristic ranking across sites, standard parametric *t*-tests assume independent observations. Because shared phylogenetic ancestry induces strong covariance across taxa, all calibrated statistical significance testing (*p*_gene_ and site-level FDR *q* ≤ 0.05) is strictly evaluated using tree-covariance-preserving Brownian motion permulations (*B* = 1,000–2,000 replicates; [102]) on covariance matrix **V** = **LL**^*T*^ (**z** = **L*ϵ, ϵ*** ~ *N* (**0, I**)).

#### 4.4.9 Worked Fine-Tuning Example: Closing the Viral Out-of-Distribution Gap

Although all primary analyses and benchmarks in this study emphasize zero-shot inference from the frozen base mammalian backbone, the modular architecture also supports lightweight supervised fine-tuning when a target regime lies far outside the pre-training distribution. Fast-evolving, shallow viral alignments constitute such a regime: the mammal-trained model transfers with moderate baseline performance out of the box. As an illustrative exercise (rather than our primary inference model), we fine-tuned the base checkpoint on approximately 9,300 empirical viral coding alignments drawn from Datamonkey [114] MEME submissions, each supplying persite MEME LRT labels (the identical ground truth the site-level model approximates). To measure genuine cross-family generalization rather than memorization, the influenza (*Orthomyxoviridae*) and coronavirus (*Coronaviridae*) families were held out entirely from fine-tuning and reserved for evaluation, and a held-out mammalian validation set was scored every epoch as a catastrophic-forgetting guardrail. Fine-tuning reuses the identical model architecture, CORAL ordinal cross-entropy loss, and dataset machinery as base training, differing only in initialization from the pre-trained weights, a gentle learning rate, and the family-level hold-out. On a single NVIDIA A100 GPU, fine-tuning required roughly 40 minutes per epoch and converged within one to two epochs (~ 2 GPU-hours total). Site-ranking Spearman *ρ* against MEME LRT improved on the *unseen* viral families from 0.403 to 0.526 (+0.123), and on in-distribution viral alignments from 0.534 to 0.606 (+0.072), while mammalian performance experienced a modest decrement (0.595 → 0.542). Thus a one-time, few-GPU-hour adaptation substantially closes the viral out-of-distribution gap while retaining the deep-time signal encoded during pre-training.

Representative command-line interface invocations across all discrete analysis workflows are summarized in Table 2.

**Table 2:** HyphAeon command-line interface: one subcommand per analysis. All commands share the `-a`/`-t`/`-o`/`-c` input/output convention.

| Analysis (Pillar) | Subcommand | Representative invocation |
| --- | --- | --- |
| Site-level selection (MEME/FEL) | `predict / meme` | `hyphaeon meme -a aln.fa -t tree.nwk -c out.csv` |
| Alignment-error filtering | `meme --filter / filter` | `hyphaeon filter -a aln.fa -t tree.nwk -c clean.csv` |
| Alignment-wide omnibus (BUSTED) | `busted` | `hyphaeon busted -a aln.fa -t tree.nwk -o out.json` |
| Epistasis, contacts, and sectors | `epistasis` | `hyphaeon epistasis -a aln.fa -t tree.nwk --graphml net.graphml` |
| Digital DMS (dDMS/ESSM) | `dms` | `hyphaeon dms -a aln.fa -t tree.nwk -c dms.csv` |
| Phenotype association | `phenotype` | `hyphaeon phenotype -a aln.fa -p marine -c pars.csv` |
| Longitudinal surveillance | `temporal / surveillance` | `hyphaeon temporal -a aln.fa -d dates.tsv -o out --plot` |
| Available model variants | `list-models` | `hyphaeon list-models` |

### 4.5 Experimental Benchmark Datasets and Multi-Scale Validation

#### 4.5.1 PRIME Multi-Gene Selection Benchmark Suite (22 Empirical Alignments)

To evaluate HyphAeon’s site-level episodic diversifying selection inference against numerical maximum likelihood across diverse taxonomic and divergence regimes, we benchmarked the 22 empirical coding alignments compiled by Kim et al. (2026) [41] (Table 1). This benchmark suite encompasses three evolutionary regimes: (1) viral pathogens and rapidly evolving surface antigens (Encephalitis Env, HIV-1 Vif, Hepatitis D Antigen, Influenza A H1N1 HA, Influenza A H3N2 HA, SARS-CoV-2 Spike, Flavivirus NS5 Polymerase); (2) mammalian immune defense and evolutionary arms races (AMELX, ADORA3, HBB, Camelid VHH Nanobodies, Abalone Sperm Lysin, Vertebrate Lysozyme, VWF); and (3) ancient conserved enzymes and structural complexes (Bacterial PTS Transporter, COL1A1, Cytochrome c Oxidase, REDIC1, ADH, RuBisCO large subunit *rbcL*, RBP3, Visual Rhodopsin *RH1*). Alignments span 16 to 483 taxa (*M*) and 96 to 1,459 codons (*L*). Benchmarking evaluated GPU wallclock inference runtime, rank concordance (Spearman *ρ*, Kendall *τ*, log-linear Pearson *r*_log_), Wasserstein distribution distance (*W*_1_), and Precision-Recall Lift at nominal *p* ≤ 0.10 against numerical HyPhy MEME (v2.5.100) point MLEs.

#### 4.5.2 Avian Macroevolutionary Ortholog Corpus (8,699 Orthologs, 39 Species)

To benchmark cross-clade zero-shot generalization on a major vertebrate radiation outside Mammalia, we evaluated the comparative avian genomic corpus curated by Shultz and Sackton (2019) [51] (Supplementary Table S10; Figs. 3, 4, 5). The corpus encompasses 8,699 orthologous coding sequence alignments spanning 39 avian species (4,674,519 total codons, 1,623,103 polymorphic codons), with an average depth of 32.6 species per ortholog (range 11–35) and lengths spanning 37 to 5,530 codons. Alignments were paired with the reference species tree from Prum et al. (2015). For each gene, HyphAeon forward inference was evaluated against background evolutionary rate constraint *ω*_0_ estimated under PAML M0 (*dN/dS* baseline), with gene-level selection aggregated via Cauchy Combination Testing (*p*_Cauchy_). To evaluate assembly quality confounding, species assemblies were matched against NCBI GenBank contig N50 metrics, comparing older short-read Illumina assemblies (e.g., *Calypte anna* bCalAnn1, contig N50 = 26.4 kb) with high-contiguity PacBio HiFi Vertebrate Genomes Project assemblies (e.g., *Calypte anna* bCalAnn2, contig N50 = 5.4 Mb). Pathway over-representation analysis was conducted across 123 curated KEGG avian pathways.

#### 4.5.3 Independent Empirical Literature Benchmark Suite (7 Studies, 73 Alignments)

To assess real-world diagnostic utility across peer-reviewed empirical studies that previously deployed HyPhy MEME, we compiled 73 published coding alignments (39,945 codons) from 7 independent investigations (Supplementary Table S9): (1) primate SMC5/6 restriction complex antagonism by Hepatitis B virus HBx regulatory protein (Abdul et al. 2018 [44]; 9 alignments, 7,073 codons); (2) primate CCDC137 evolutionary constraint under HIV-2/SIV Vpr targeting (Nisson et al. 2025 [49]; 1 alignment, 290 codons); (3) bat and primate GBP5 GTPase diversification against lentiviruses (Le Corf et al. 2026 [46]; 2 alignments, 1,223 codons); (4) horseshoe bat OAS1 viral RNA-sensing loop diversification (Lytras et al. 2023 [45]; 1 alignment, 351 codons); (5) TRMT1 tRNA methyltransferase cleavage by SARS-CoV-2 main protease (D’Oliveira et al. 2025 [47]; 2 alignments, 1,613 codons); (6) Siglec and C-type lectin families across primates, rodents, and bats (Hilbert and Elde 2023 [50]; 57 alignments, 28,462 codons); and (7) progesterone receptor (PGR) regulatory divergence in mammalian pregnancy (Marinić and Lynch 2020 [48]; 1 alignment, 933 codons). Zero-shot predicted test statistics 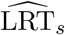 were benchmarked against published numerical MEME outputs via non-parametric rank correlation (*ρ*).

#### 4.5.4 OrthoMaM v12 Mammalian Gene Trees Curation and Assembly Contiguity Metadata

To evaluate selection scaling across the mammalian radiation, we compiled 15,868 orthologous gene families from OrthoMaM v12 [53], spanning up to 190 mammalian species. Alignments map 100% (15,868/15,868) to NCBI Entrez Gene IDs and human HGNC orthologs. Gene trees were inferred via maximum likelihood with branch lengths in expected substitutions per codon. For each mammalian taxon, genome assembly contiguity metadata (contig N50 and scaffold N50) were retrieved programmatically from the NCBI Datasets API. Log-linear regression models quantified the scaling of candidate selected sites (*p* ≤ 0.05) against contig N50, coding length, and total tree divergence length, cross-referencing candidate sites against whole-gene BUSTED-E selection tests.

#### 4.5.5 Clade-Specific Subtree Selection Profiling Across Five Mammalian Orders

To map lineage-specific episodic selection across distinct mammalian orders without taxonomic subsampling artifacts, the 15,868 OrthoMaM alignments were partitioned into five monophyletic mammalian clades: Primates (*N* = 32 species), Chiroptera (*N* = 41 species), Cetartiodactyla (*N* = 38 species), Carnivora (*N* = 48 species), and Rodentia (*N* = 45 species). Trees were pruned to clade-specific subtrees, and each clade was evaluated independently via `hyphaeon predict`. Sitewise selection density was normalized by taxon count to compute Taxon-Normalized Selection Density:

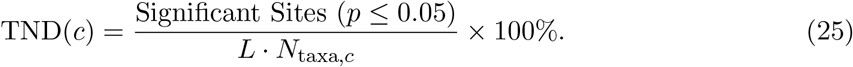

Pairwise clade overlap was quantified using the Jaccard similarity index across significant codon positions 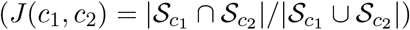, constructing order-level adaptive sharing matrices.

#### 4.5.6 HIV-1 Reverse Transcriptase Clinical Resistance Benchmark Cohort

To evaluate model sensitivity under extreme out-of-distribution domain shift (dense intra-host clinical sampling, transient neutral polymorphisms, and Sanger primer missing data), we examined 475 clinical HIV-1 subtype C Reverse Transcriptase (RT) sequences (*L* = 335 codons) from a South African antenatal cohort following single-dose nevirapine (sdNVP) exposure (Seoighe et al. 2007 [36]; Section 2.1.1, Fig. 2). Codons 1–34 exhibit 29.8% missing data due to sequencing primer binding sites. Clinical ground truth was compiled from the Stanford HIV Drug Resistance Database (HIVdb v9.5) and the International Antiviral Society–USA (IAS-USA) guidelines, identifying 37 sequenced Drug Resistance-Associated Mutation (DRAM) positions in the cohort, including seven canonical NVP resistance loci (L100I, K103N, V106M, V108I, Y181C, Y188C, G190A) and five multi-class NRTI/NNRTI resistance anchors (K65R, K70R, E138K, K219W, K238T). Performance was benchmarked zero-shot using `hyphaeon predict` against numerical MEME via AUROC, AUPRC, and sensitivity at nominal *p* ≤ 0.10 and FDR *q* ≤ 0.20.

#### 4.5.7 Influenza A Nucleoprotein Experimental DMS Preference Landscape Benchmark

To assess whether differences between neural foundation predictions and numerical maximum likelihood track biophysical properties, we benchmarked HyphAeon on Jesse Bloom’s dataset of 274 human Influenza A H1N1 Nucleoprotein (NP) isolates (*L* = 498 codons, 428 polymorphic; Bloom 2014 [39]; Section 2.1.2) spanning circulating lineages from 1918 to 2012. Sitewise selection predictions 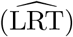 were cross-referenced against experimentally determined amino acid preference distributions derived from deep mutational scanning (DMS) libraries in MDCK cells. Mutational tolerance was quantified by effective number of amino acids (*N*_eff_ = exp(*H*_*s*_), where *H*_*s*_ is Shannon entropy over experimental preferences). Relative solvent accessibility (RSA) was calculated from the crystal structure of the H1N1 NP trimer (PDB 2IQH) using DSSP. Sitewise selection discrepancy 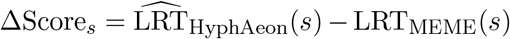 was evaluated across functional domains, including the CTL escape epitope R384G, basic RNA-binding cleft residue K214, and terminal polymorphism site D290.

#### 4.5.8 HIV-1 Protease Clinical Epistatic Co-Selection and Potts Benchmark Compilation

To validate zero-shot epistatic co-selection networks and sector mining against statistical physics baselines, we analyzed the clinical HIV-1 subtype B protease dataset compiled by Levy et al. (2017) [68] and Flynn et al. (2017) [69] (*L* = 99 codons, *N* = 4,499 unique clinical haplotypes, *M* = 8,996 phylogenetic branches; Section 2.8, Fig. 8). The alignment captures dense multidrug-experienced patient sequences with established allosteric flap dynamics, active site pocket geometry (PDB 1HHP), and multi-class protease inhibitor resistance mutations. Pairwise epistatic co-selection was evaluated via `hyphaeon epistasis` using attribution cosine similarities with Average Product Correction (**C**^APC^) and the Composite Epistatic Selection Index (CESI) at FDR *q* ≤ 0.05. Discovered sectors were evaluated via Spectral Coherence (*C*(*S*)) and tested against 100,000-replicate Monte Carlo graph permutations and search-aware neutral clustering simulations (*C*_null_ ≤ 0.450), directly comparing network recovery and runtime against inverse Potts models and Direct Coupling Analysis (DCA).

#### 4.5.9 Vertebrate Dim-Light Rhodopsin (RH1) Ancestral Resurrection Benchmark

To evaluate directional phenotype–genotype attribution against experimentally resurrected functional ancestral mutations, we analyzed the 38-species vertebrate Rhodopsin (RH1) dataset (*L* = 330 codons; Yokoyama et al. 2008 [83]; Section 2.11, Fig. 10). Yokoyama and colleagues experimentally resurrected 11 ancestral pigments in vitro, determining absorption spectra (*λ*_max_) and identifying 15 specific amino acid replacements across 12 critical sites responsible for blue-shifted dim-light vision (*λ*_max_ ∈ [480, 526] nm vs. 500 nm terrestrial baseline). Directional selection attributions (**b**_*s*_ ∈ ℝ^38^) were projected onto the normalized blue-shift adaptation phenotype vector (**ŷ**) to calculate directional concordance 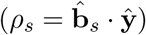 and Student’s *t*-test significance at FDR *q* ≤ 0.05. Candidate sites were clustered into four epistatic sectors via Two-Stage Seed-and-Extend modularity optimization, with empirical significance confirmed via 20,000-replicate Monte Carlo graph permutations, and benchmarked for Positive Predictive Value (PPV) lift against classical PAML M2a/M8 Bayes Empirical Bayes predictions.

#### 4.5.10 Combinatorial Molecular Convergence (CSUBST) Multi-System Benchmark Suite

To compare continuous directional attribution against combinatorial ancestral convergence models, we benchmarked HyphAeon on empirical multi-species convergence systems compiled from Fukushima and Pollock (2023) [34] (Section 2.11.3, Fig. 11, Supplementary Table S5): (1) mammalian echolocation prestin (SLC26A5, 128 species, *L* = 758 codons; patch-clamp validated sites N14T and I392T); (2) ouabain toxin resistance in ATP1A1 (30 species, *L* = 1,045 codons; CRISPR transgenic in vivo validated sites Q111V/L and A119S); (3) C_4_ photosynthetic PEPC2 (71 species, *L* = 971 codons; catalytic pocket site Ala572Ser and regulatory phosphorylation site Ser780); (4) digestive enzyme adaptation in colobine primate RNASE1 (36 species, *L* = 156 codons; in vitro validated sites R4Q, K6N, R39W) and ruminant stomach lysozyme *c* (288 species, *L* = 176 codons); and (5) carnivorous pitcher plant secreted hydrolases (RNase T2, GH19 endochitinase, and parasitic haustorial Cuscuta transfers). Predictions were classified under a 3-tier validation schema: Tier 1 (Directly Experimentally Validated), Tier 2 (Structurally/Biochemically Supported), and Tier 3 (Novel Computational Discovery), and evaluated for sitewise concordance (*ρ*_*s*_, *q* ≤ 0.05) and multi-site epistatic sectors (*p*_perm_ ≤ 0.05, 20,000 permutations).

#### 4.5.11 64-Grass Plastid Discovery Cohort (Allard et al. 2025)

To identify the core chloroplast genes driving C_4_ photosynthetic adaptation prior to whole-plastome screening, we analyzed 67 plastid protein-coding genes curated across 64 grass species (Poaceae; Grass Phylogeny Working Group II cohort [35]) spanning both C_3_ and C_4_ lineages. Directional selection attribution matrices 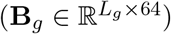 were projected onto the C_3_/C_4_ binary phenotype indicator vector. Statistical significance was evaluated via 2,000 tree-covariance-preserving Brownian motion permulations per gene, discovering the 17 permutation-significant chloroplast genes (*q*_gene_ ≤ 0.05) harboring 219 Phenotype-Associated Residue Signatures (PARS).

#### 4.5.12 Mechanistic Probing of Embeddings and Biophysical Property Correlates

To determine what biochemical and evolutionary features the frozen 384-dimensional representation space captures, we conducted linear probing experiments on the internal token and site embeddings. Codon representations (*d*_codon_ = 192) were probed against Position 3 wobble GC composition (GC3), purine/pyrimidine content, and steric volume across all 61 sense codons (and 64 triplet codons). Non-synonymous amino acid representations (*d*_aa_ = 192) were probed against the five canonical Atchley biophysical factors (Factor 1: polarity/hydrophobicity; Factor 2: secondary structure propensity; Factor 3: molecular size/volume; Factor 4: codon diversity; Factor 5: electrostatic charge [115]). Attention head Markov substitution transition decay rates *λ*_*h*_ were examined across all *H* = 12 heads. To verify the insulation of synonymous rate variation from positive selection, Frobenius norm ratios between the *dS* projection weights and *dN* cross-coupling weights (‖**W**_*dS*_‖_*F*_ /‖**W**_*dS,dN*_ ‖_*F*_) were computed across all transformer layers.

#### 4.5.13 Pan-Pathogen Longitudinal Genomic Surveillance Datasets

To evaluate continuous temporal sweep velocity regression across diverse biological and epidemiological regimes, we curated ten longitudinal surveillance datasets from Nextstrain [23, 24] (https://data.nextstrain.org; Section 2.12, Figs. 13, 14; Supplementary Table S12): (1) SARS-CoV-2 Spike (Surveillance / Pandemic, *N* = 1,863 unique haplotypes, 78 months, December 2019 to mid-2026); (2) Influenza A/H3N2 HA1 (54-year human surveillance benchmark, 1968–2022, *N* = 1,828 genomes); (3) Avian Influenza A/H5N1 Cattle HA (2024–2026 US dairy outbreak, *N* = 989 genomes); (4) Avian Influenza A/H5N1 Cattle PB2 (Polymerase, *N* = 1,215 genomes); (5) Dengue Virus 2 Envelope (Asian/American genotype displacement, 1944–2024, *N* = 2,032 genomes); (6) Rabies Lyssavirus Glycoprotein G (Host jumping across carnivore clades, 1950–2026, *N* = 2,272 genomes); (7) Enterovirus D68 Capsid VP1 (Biennial pediatric acute flaccid myelitis outbreaks, 1997–2025, *N* = 1,210 genomes); (8) Zika Virus Envelope (Epidemic emergence, *N* = 519 genomes); (9) Mycobacterium tuberculosis KatG (Isoniazid resistance, *N* = 76 clinical isolates); and (10) Mycobacterium tuberculosis RpoB (Rifampicin resistance, *N* = 163 clinical isolates). Datasets encompass 12,167 timestamped genomes (10,034 unique haplotypes) and 6,679 codons (2,014 variable codons). All datasets were ingested with collection date metadata and evaluated via `hyphaeon temporal`.

#### 4.5.14 TEM-1 *β*-Lactamase Experimental DMS Benchmark

We compiled an alignment of 85 natural *β*-lactamase bacterial homologs spanning 263 mature codon positions (789 nt) remapped from the 286-amino acid precursor [77, 116]. For all 4,997 point mutations (263 *×* 19), zero-shot selection disruptions (|ΔLRT|) and Intrinsic Genetic Plasticity (Φ_*s*_) were evaluated against experimental ampicillin minimum inhibitory concentrations (2,500 *µ*g/mL).

#### 4.5.15 ProteinGym Human DMS Benchmark Compilation and Dynamic Programming Remapping

Human DMS assays were extracted from ProteinGym [15] and mapped to mammalian coding alignments in TOGA (*N* = 18,277 genes). To resolve truncated isoforms and synthetic expression tags, canonical assay sequences **s**_canon_ were aligned to TOGA translations **s**_TOGA_ via global Needleman-Wunsch dynamic programming (`Bio.Align.PairwiseAligner`; match +2, mismatch −1, gap open −10, gap extend −0.5). Assays were filtered by strict wild-type concordance and an alignment integrity threshold (sequence identity ≥ 85%, mapping rate ≥ 50%). Across 26 passing high-quality assays (119,116 validated missense mutations across 21 disease genes), zero-shot fitness scores were evaluated via Spearman *ρ*, Pearson *r*, and AUROC on binarized functional thresholds.

#### 4.5.16 High-Throughput Plastome Phenomics and C_3_/C_4_ Photosynthetic Classification

We ingested the complete NCBI RefSeq Plastid Protein Release (1,315,257 records, 163.6 MB) via in-memory stream processing, mapping records across 15,168 land plant plastomes to 17 permutation-significant chloroplast genes (*rbcL, matK, ndhD/E/F/G/H, rpoA/B/C2, atpB/E, psaI, clpP, rps3/18, rpl14*) using C-accelerated dynamic programming (*<* 0.25 ms per protein). Using 219 Phenotype-Associated Residue Signatures (PARS; *q*_gene_ ≤ 0.05), the composite C_4_ score is computed as:

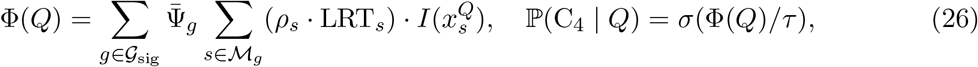

where 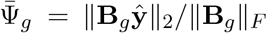 is length-normalized spectral energy and 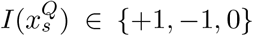 matches derived C_4_, ancestral C_3_, or ambiguous states (*τ* = 1.5). Whole-proteome scoring across all 15,168 species completed in 11.9 seconds on 17 CPUs (*>* 1,270 plastomes/s). Benchmarking against the Osborne et al. (2014) global grass database [103] (*N* = 944 matched RefSeq plastomes; 545 C_3_ vs. 399 C_4_) demonstrated 77.69% sensitivity, 79.08% specificity, and an agricultural cohort ROC-AUC = 0.9250 across major cereal crops (*Zea, Sorghum, Oryza, Triticum, Hordeum*).

#### 4.5.17 Global Clinical Malaria Cohort Curation and Genomic Data Extraction

To evaluate epistatic sector discovery and digital DMS at population scale, we curated clinical isolates from the MalariaGEN *Plasmodium falciparum* Release 7 (Pf7) open dataset [117], comprising 20,864 clinical blood samples collected across 33 endemic countries in Africa, Southeast Asia, South America, and Oceania. Primary sample manifests and clinical metadata (Pf7_samples.txt) were cross-referenced against high-confidence drug resistance marker genotypes (`Pf7_drug_resistance_marker_genotypes.txt`) to extract isolate geographic coordinates, World Health Organization transmission regions, collection year, and partner study identifiers.

For the chloroquine resistance transporter *PfCRT* (`PF3D7_0709000`; *L* = 424 codons), genomic variant calls and locus-specific haplotype matrices (`Pf7_crt_haplotypes.txt`) were retrieved from the MalariaGEN public data repository and Wellcome Open Research open-access archives. To ensure translational and reading-frame fidelity across surveillance isolates, records were filtered under standard quality criteria: each patient sequence was verified for an exact full-length coding sequence (1,275-nt / 424 codons), strict in-frame triplet preservation without phase-shifting insertions or deletions, zero internal stop codons, and complete absence of uncalled or ambiguous nucleotide characters (“N”). This curation yielded 12,389 full-length patient sequences.

To benchmark operational performance under dense surveillance streaming, cohorts were sampled at two scales: (1) a geographically stratified cohort of *N* = 4,096 field isolates spanning West Africa (Ghana, Mali, Gambia), Southeast Asia (Cambodia, Vietnam, Laos, Thailand, Myanmar), South America (Colombia), and Oceania (Papua New Guinea); and (2) the full MalariaGEN Pf7 clinical surveillance cohort of *N* = 12,389 isolates (collapsing into *M*_unique_ = 157 unique evolutionary haplotypes across 33 endemic countries), with scalability stress tests evaluated up to *N* = 32,768 isolates (2^15^ sequences; 41.8 MB FASTA). Unaligned sequences were compiled into in-frame codon matrices, and maximum-likelihood phylogenies were reconstructed using FastTree (v2.1.11 [118]; Generalized Time-Reversible model with CAT approximation, `-nt -gtr`). Zero-length terminal branch polytomies were assigned minimum branch lengths (*c* = 10^−4^).

Epistatic networks were inferred via `hyphaeon epistasis` using the base foundation checkpoint. Identical duplicate sequences were collapsed dynamically in memory (`prune_duplicates=True`) into unique evolutionary haplotypes (*M*_unique_ = 88 for *N* = 4,096; *M*_unique_ = 157 for *N* = 12,389–32,768) while tracking occurrence weights **w** for branch attribution. Co-selection was evaluated across all 424 *×* 423/2 = 89,676 residue pairs using the Composite Epistatic Selection Index (CESI) at FDR *q* ≤ 0.05, followed by two-stage seed-and-extend spectral graph clustering to extract cohesive allosteric sectors. Digital deep mutational scanning evaluated all 19 non-synonymous amino acid substitutions per position to compute Intrinsic Genetic Plasticity Φ_*s*_ (377.5 mut/s on CPU).

Multi-species comparative analyses for *Kelch13, PfATP4, PfPI4K, PfPMX, PfKRS1*, and *PfDHODH* used wild-type RefSeq orthologs across 8–18 *Plasmodium* species (Laverania, Simian, Human, and Rodent clades).

### 4.6 Controlled Synthetic Markov Simulation Suite and 14 Evolutionary Regimes

To benchmark statistical sensitivity and error calibration across known ground truth, we implemented a continuous-time Markov substitution generator using Pyvolve [60] under the Muse and Gaut 1994 (MG94) codon framework [2]. For 61 sense codons *C* with non-synonymous ratio *ω* and equilibrium frequencies ***π*** ∈ Δ^60^, instantaneous substitution rates are governed by **Q** ∈ ℝ^61*×*61^:

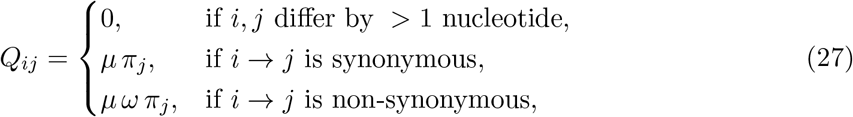

where *Q*_*ii*_ = − ∑_*j*≠*i*_ *Q*_*ij*_ and *µ* normalizes expected substitution rate to unit branch length (−∑*π*_*i*_*Q*_*ii*_ = 1). Finite-time transition probabilities across branch length *b* are computed via eigendecomposition **P**(*b*) = exp(**Q***b*) = **V** exp(**Λ***b*)**V**^−1^.

We generated 14 distinct simulation regimes (100 Monte Carlo replicates each, *L* = 200 codons; *N* = 1,400 alignments, 280,000 codons): (1) *Pervasive Strong* (*N* = 64, *T* = 8.0, *ω*^+^ = 12.0, *f*_fg_ = 100%); (2) *Pervasive Moderate* (*N* = 64, *T* = 8.0, *ω*^+^ = 4.0, *f*_fg_ = 100%); (3) *Deep Clade Bursts* (*N* = 96, *T* = 12.0, *ω*^+^ = 25.0, *f*_fg_ = 50%); (4) *Moderate Clade Bursts* (*N* = 96, *T* = 12.0, *ω*^+^ = 8.0, *f*_fg_ = 50%); (5) *Sparse Subclade Episodic* (*N* = 128, *T* = 16.0, *ω*^+^ = 50.0, *f*_fg_ = 20%); (6) *Ultra-Sparse Single-Lineage* (*N* = 128, *T* = 16.0, *ω*^+^ = 150.0, *f*_fg_ = 5%); (7) *Deep Vertebrate Divergence* (*N* = 190, *T* = 25.0, *ω*^+^ = 15.0, *f*_fg_ = 30%); (8) *Rapid Radiation* (*N* = 40, *T* = 1.5, *ω*^+^ = 35.0, *f*_fg_ = 40%); (9) *Purifying Null Control* (*N* = 64, *T* = 8.0, *ω* = 0.15, *f*_fg_ = 0%); (10) *Neutral Drift Null Control* (*N* = 64, *T* = 8.0, *ω* = 1.00, *f*_fg_ = 0%); (11) *TOGA Primates Subtree* (*N* = 32, *T* = 3.1, *ω*^+^ = 20.0, *f*_fg_ = 35%); (12) *TOGA Carnivora Subtree* (*N* = 48, *T* = 5.4, *ω*^+^ = 25.0, *f*_fg_ = 30%); (13) *TOGA Chiroptera Subtree* (*N* = 41, *T* = 4.6, *ω*^+^ = 35.0, *f*_fg_ = 40%); and (14) *Ultra-High Power Pervasive* (*N* = 128, *T* = 35.0, *ω*^+^ = 120.0, *f*_fg_ = 100%). Datasets were evaluated in parallel on 32-core MPI HyPhy MEME (v2.5.101) cluster jobs vs. HyphAeon forward inference.

#### 4.6.1 Active Learning and Gaussian Process Parameter Space Exploration

To map the continuous multi-dimensional power landscape and identify the 50% and 80% isopower detection frontiers without intractable grid search, we implemented an active learning framework guided by a Gaussian Process (GP) surrogate with a Matérn 5/2 covariance kernel. The GP model iteratively explores the 4D evolutionary parameter space: taxonomic depth (*N*_taxa_ ∈ [32, 384]), tree divergence length (*T*_depth_ ∈ [3.0, 25.0]), selection intensity (log_10_ *β*_fg_ ∈ [log_10_(2.0), log_10_(500.0)]), and foreground branch fraction (*f*_fg_ ∈ [0.02, 1.00]). At each iteration, new simulation coordinates were sampled via Expected Improvement (EI), generating Pyvolve replicates to update the surrogate response surface, construct power contours (Fig. 6), and delineate the empirical boundaries of episodic selection detectability.

### 4.7 Systematic Architectural Ablation Methodology

To empirically isolate the individual contributions of each architectural innovation, we conducted ablation experiments across all 1,400 paired synthetic validation alignments (280,000 codons; Supplementary Table S8):

1. *Impact of 4D Tree-RoPE* : Replacing 4D metric Tree-RoPE with standard 1D linear positional encodings caused a severe drop in selection ranking concordance (*ρ* dropped from 0.355 to 0.171, Δ*ρ* = −0.184) and collapsed PR-AUC from 0.364 to 0.221, demonstrating that geometric rotary phase rotation is essential for conditioning on continuous branch lengths and phylogenetic divergence.
2. *Impact of BlockLinear Disentanglement* : Replacing the block-diagonal *dS/dN* projector (192-dim each) with an unconstrained linear layer inflated the false positive rate on neutral null regimes from FPR_0.05_ = 0.0307 to 0.1574 (+412%). Without block-diagonal separation, base composition, GC3 wobble, and synonymous rate variation (*dS*) leak into non-synonymous channels, triggering spurious positive selection predictions.
3. *Impact of Continuous-Time Markov Kernel* : Removing the distance-dependent exponential prior 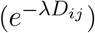 and relying purely on learned Query-Key inner products reduced concordance (*ρ* = 0.263, Δ*ρ* = −0.092), particularly on deep-divergence regimes (*T >* 20.0) where unweighted attention over-weights saturated ancient taxa.
4. *Impact of CORAL Rank-Consistent Ordinal Head* : Replacing the 16-threshold ordinal survival head with standard Mean Squared Error (MSE) regression resulted in catastrophic failure (*ρ* = 0.082, PR-AUC = 0.082). In natural alignments where the vast majority of codons are strictly neutral or conserved (LRT *<* 0.1), unconstrained MSE collapses toward the zero-mean, predicting flat near-zero scores and losing all ranking resolution on true episodic sweeps.

### 4.8 Adaptive Hardware-Aware Memory Management, Attention Scaling, and Tree Subsampling

To guarantee execution stability across arbitrary sequence depths on heterogeneous accelerators without out-of-memory crashes, HyphAeon explicitly manages the quadratic scaling of column-wise row attention and incorporates phylogenetic subsampling for large trees.

#### 4.8.1 Quadratic Attention Memory Derivation and Hardware Profiles

The primary hardware constraint during transformer forward inference is the unnormalized Query– Key self-attention logit tensor **A** ∈ ℝ^*B×H×M×M*^ across *H* = 12 attention heads (*d*_head_ = 32). In standard 32-bit floating-point precision, storing the activation tensor for a single attention layer requires:

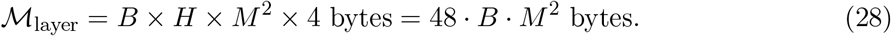

During non-iterative forward evaluation, activations are reclaimed across the 6 sequential transformer layers, bounding peak attention memory to ≈ 48 · *B* · *M* ^2^ bytes per forward step. This relationship dictates realistic sequence depth limits across contemporary accelerator tiers:

- *Consumer Workstations and Laptops (*8*–*16 *GB VRAM / Unified Memory)*: On devices such as NVIDIA RTX 3070/4070 or Apple Silicon M-series unified memory, alignments of *M* ≤ 500 taxa require only 48 *×* 64 *×* 500^2^ ≈ 0.77 GB per layer at batch size *B* = 64. For *M* ≈ 1,000 taxa (48 MB per codon site), micro-batched streaming at *B* = 16–32 maintains peak attention between 0.77–1.54 GB, executing smoothly without exhausting memory headroom.
- *Datacenter Accelerators (*40*–*80 *GB VRAM)*: On enterprise hardware (such as NVIDIA A100 or H100), alignments of *M* ≈ 2,000 taxa require 192 MB per codon site (12.3 GB at standard batch size *B* = 64), executing natively on a single GPU. For alignments approaching *M* ≈ 4,000 taxa (768 MB per site), micro-batching at *B* = 8–16 constrains peak attention memory to 6.1–12.3 GB, permitting direct evaluation without distributed model parallelization.

#### 4.8.2 Dynamic Safe Batch Sizing

To automatically adapt to available device memory, HyphAeon dynamically computes device memory budgets:

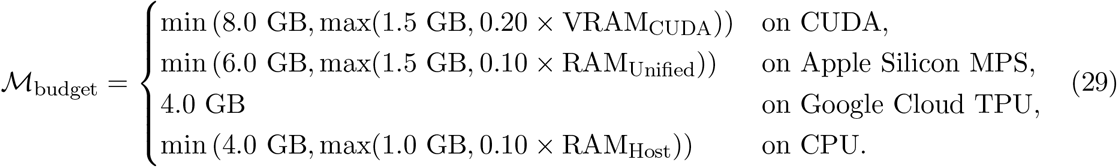

The safe batch size is computed as *B*_safe_ = max 1, ⌊*M*_budget_/(48 · *M* ^2^) ⌋. For digital deep mutational scanning (dDMS / ESSM), batch sizes are quantized to integer multiples of 19 (*B* = 19 *× k*) to maintain maximal tensor core occupancy across complete amino acid substitution sweeps.

#### 4.8.3 Faith’s Phylogenetic Diversity (PD) Subsampling for Large Trees

Tree positional encodings in HyphAeon are scale-free: rather than assigning taxa to discrete positional tokens bounded by fixed vocabulary lengths, continuous metric Multi-Dimensional Scaling (MDS) projects pairwise patristic distances into a continuous 4D manifold (**P**_MDS_ ∈ ℝ^*M×*4^) that accommodates arbitrary taxonomic depths without retraining. When empirical trees exceed accelerator memory budgets (*M* ≫ 2,000 species in tree-of-life phylogenomics), HyphAeon incorporates an automated greedy Faith’s Phylogenetic Diversity (PD) subsampling algorithm [119].

Let *T* = (*V*, ℰ, *ℓ*) be a rooted phylogenetic tree with edge lengths *ℓ*(*e*). For any taxon subset *S* ⊆ *X*, Faith’s PD is defined as the sum of branch lengths spanning the subtree induced by *S* ∪ *{*root*}*: PD(*S*) = ∑_*e*∈*E* (*S*)_ *ℓ*(*e*). Given a target compute budget *k ≪ M*, the subsampler initializes with the phylogenetic diameter pair *S*_2_ = *{u, v}* = arg max_*x,y*∈*X*_ *d*_*T*_ (*x, y*). At each subsequent iteration *i* ∈ *{*3, …, *k}*, the algorithm greedily adds the taxon maximizing marginal phylogenetic diversity:

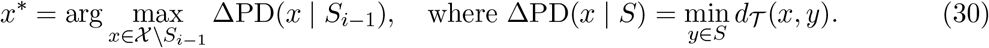

By submodularity of edge coverage, this greedy selection provably achieves a (1 − 1*/e*) ≈ 63.2% approximation to NP-hard optimal branch coverage (empirically capturing *>* 92% of total tree divergence), compressing deep phylogenies to representative subsets that fit within accelerator memory without taxonomic sampling bias.

#### 4.8.4 Population-Scale Haplotype Pruning

In dense viral surveillance or clinical epidemiology cohorts (such as SARS-CoV-2, HIV, or *Plasmodium* field isolates) where thousands of patients share identical clonal lineages, computing pairwise attention across identical sequences is computationally redundant. HyphAeon incorporates automated duplicate pruning (`prune_duplicates=True`), which collapses identical sequences across alignment columns into *M*_unique_ unique evolutionary haplotypes while tracking duplicate frequencies 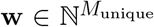 for weighted branch attribution. On large clinical cohorts (e.g., *N* = 32,768 field isolates), this collapses the working matrix from 32,768 leaves to ≈ 150 unique haplotypes in memory, executing full epistatic sector mining and selection testing in *<* 6 seconds on CPU without quadratic attention penalties.

#### 4.8.5 Tree-Free Evolutionary Inference via Direct Pairwise Genetic Distances (TN93)

In conventional molecular evolution workflows, phylogenetic trees represent a significant computational bottleneck and source of fragility: users must reconstruct branching topologies (e.g., via maximum likelihood or neighbor joining), resolve multifurcations, reconcile taxon naming and capitalization discrepancies, and numerically optimize continuous-time Markov branch lengths. Numerical optimization frequently encounters convergence failures or severe branch-length inflation on alignments containing alignment gaps or shallow divergence.

Because HyphAeon factorizes phylogenetic structure entirely through the pairwise distance metric **D** ∈ ℝ^*M×M*^ (via continuous substitution decay exp(−*λ***D**) and 4D spectral MDS coordinates **P**_MDS_), the model does not require an explicit bifurcating tree topology. HyphAeon can operate in an automated tree-free mode, bypassing tree construction entirely by computing all-pairs nucleotide distances directly from the alignment using the Tamura–Nei 1993 model (`tn93`) [120]:

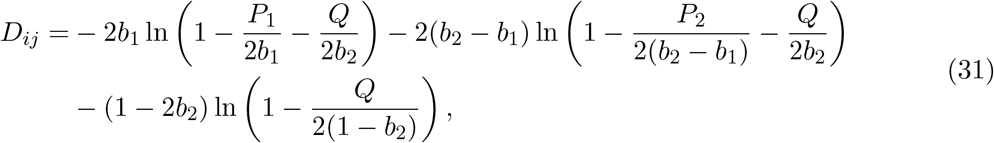

where *P*_1_, *P*_2_ denote transition frequencies (purine and pyrimidine), *Q* denotes transversion frequency, and *b*_1_, *b*_2_ are base frequency parameters. Pairwise distances are computed across all unique sequence pairs in *O*(*M* ^2^*L*) time (*<* 50 ms per alignment), double-centered, and projected via classical MDS into **P**_MDS_ ∈ ℝ^*M×*4^.

Benchmarking across all 24 empirical datasets (*N* = 9,998 codons) demonstrated near-perfect sitewise concordance between standard Tree-Patristic HyphAeon and Tree-Free TN93 HyphAeon: median Spearman rank correlation *ρ* = 0.9997, median Pearson linear correlation *r* = 0.9997, and median Mean Absolute Difference MAD = 0.021 LRT units (with 20 of 24 datasets exhibiting *ρ* ≥ 0.9992). Concordance with ground-truth numerical HyPhy MEME was statistically indistinguishable between Tree-Patristic (*ρ* = 0.3234) and Tree-Free TN93 (*ρ* = 0.3424; *p* = 0.305, paired *t*-test). In practice, TN93 proved resilient against numerical tree failure modes: on *AMELX* (where missing data caused numerical tree branch optimization to inflate distances by 230*×*, corrupting tree-based predictions to *ρ* = −0.2216 vs. MEME), direct TN93 pairwise distances circumvented branch length distortion, restoring positive concordance with MEME (*ρ* = +0.2131). Across all benchmarks, the tree-free pipeline completed in 8.47 seconds (1.6*×* faster than tree parsing and branch estimation), allowing users to execute foundation-model evolutionary inference directly on raw FASTA alignments without phylogenetic overhead.

### 4.9 Real-Time Pan-Pathogen Genomic Surveillance Pipeline (Nextstrain Benchmark)

To establish a scalable real-time surveillance workflow on open public health data, we developed an automated pipeline that interfaces directly with phylogenetic datasets from Nextstrain [23, 24] (https://data.nextstrain.org). We established two complementary surveillance benchmark suites: (1) a cross-sectional multi-stage surveillance suite comprising seven high-priority open Nextstrain pathogen datasets evaluated for episodic selection, epistatic co-selection networks, and directional phenotype attribution (Avian Influenza A/H5N1 HA and PB2, SARS-CoV-2 Spike, Dengue 2 E, Rabies G, Enterovirus D68 VP1, and Zika E; Supplementary Table S11); and (2) a longitudinal continuous temporal surveillance suite comprising ten epidemic cohorts spanning RNA viruses, DNA viruses, and clonal bacteria evaluated for time-resolved sweep velocities, functional dynamic factor waves (fPCA), and two-stage statistical filtering (SARS-CoV-2 Spike, Influenza H5N1 HA, Influenza H3N2 HA, Dengue 1 E, Dengue 2 E, RSV-A G, RSV-B G, Rabies G, Enterovirus D68 VP1, and *Mycobacterium tuberculosis KatG* /*RpoB*; Supplementary Table S12).

#### 4.9.1 Surveillance Ingest, Alignment Processing, and Tree Cache Construction

For all surveillance datasets, genomic data are ingested from JSON archives or FASTA alignments containing ancestral sequence reconstructions, full nucleotide/amino-acid alignments, and reconstructed phylogenetic trees or date-stamped tip metadata.

To process large transmission phylogenies with thousands of genomes without loss of evolutionary diversity or artificial downsampling:

1. *Duplicate Haplotype Pruning* : Identical terminal leaves are collapsed into unique evolutionary haplotypes while preserving tree topology, edge lengths, and lineage counts. This guarantees that tree inference scales with true sequence diversity without losing rare singletons or subsampling trees to arbitrary batch limits (e.g., *N* = 1,863 unique haplotypes retained from 3,839 SARS-CoV-2 Spike genomes; *N* = 2,272 retained from 3,087 Rabies genomes).
2. *Invariable Site Pre-Filtering* : Codon columns exhibiting zero nucleotide or amino acid variation across all sampled taxa are screened in *O*(*N* · *L*) time. Invariable sites are analytically assigned LRT_*s*_ = 0.0 and *p*_*s*_ = 1.0, allowing GPU tensor operations to focus exclusively on active polymorphic codons (*L*_var_).
3. *Tree Distance Cache*: Pairwise patristic distance matrices **D** ∈ ℝ^*N×N*^ and root-to-node divergence depths **z** ∈ ℝ^*N*^ are extracted via tree traversal and cached in GPU memory for fast contextual positional encoding.

#### 4.9.2 Multi-Stage Surveillance Workflow: Selection, Epistasis, and Phenotype Attribution

For each surveillance target, the complete evolutionary profile is evaluated through the frozen HyphAeon foundation transformer (`PhyloAxialTransformer`, 1.91M parameters, 6 axial attention layers, 12 attention heads, embedding dimension 384; Section 4.1):

1. *Episodic Diversifying Selection*: Tokenized codon sequences **C** ∈ *{*0, …, 64*}*^*B×N×L*^ and amino acid tokens **A** ∈ *{*0, …, 20*}*^*B×N×L*^ for all variable codons are evaluated through the 6-layer species transformer in safe hardware-adapted batches. Predicted likelihood ratio statistics LRT_*s*_ = −2 log Λ_*s*_ are mapped to asymptotic *p*-values via the canonical mixture null (Methods 4.4.1, Equation (12)), with multiple testing controlled via Benjamini-Hochberg FDR (*q*_*s*_ ≤ 0.05 and *q*_*s*_ ≤ 0.10).
2. *Axial Attention Epistasis and Sector Mining* : Continuous lineage attribution vectors **a**_*s,n*_ = *α*_root→*n,s*_ · *δ*_*s,n*_ are extracted from root-to-leaf attention routing in layer 6 across all 12 heads. Pairwise co-selection is evaluated via attribution cosine similarity Sim(*u, v*), tested for significance via exact one-tailed Student’s *t*-tests (*q*_*u,v*_ ≤ 0.05), and prioritized via the Composite Epistatic Selection Index (CESI_*u,v*_ ≥ 1.5–2.0; Methods 4.4.5). Co-selection subgraphs are partitioned into epistatic sectors via Two-Stage Seed-and-Extend modularity clustering and validated against Spectral Coherence (*C*(*S*)) and Monte Carlo graph permutations.
3. *Directional Phenotype Attribution via Cauchy Combination*: To identify substitutions driving epidemiological adaptations (such as bovine spillover in H5N1, domestic carnivore host adaptation in rabies, or clades in enterovirus D68), terminal taxa are mapped to phenotypic trait indicators **y** ∈ *{*0, 1*}*^*N*^. Directional concordance is quantified by *ρ*_*s*_ = Corr(**a**_*s*_, **y**), with significance assessed via tree-covariance-preserving Brownian motion permulations (*B* = 1,000 replicates; Methods 4.4.8). To combine evidence from episodic selection (*p*_LRT,*s*_) and trait association (*p*_*ρ,s*_), HyphAeon applies the Aggregated Cauchy Association Test (ACAT [52]):

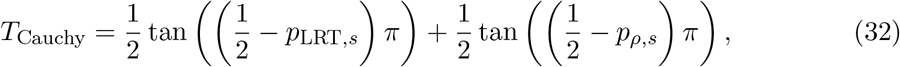

yielding closed-form omnibus significance 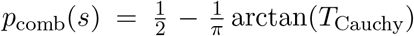 robust to arbitrary correlation between selection and trait association under locus-wide FDR control (*q* ≤ 0.05).

#### 4.9.3 Continuous Temporal Attribution Regression and Two-Stage Filtering

To enable time-resolved evolutionary inference without the discretization artifacts and power collapse of sliding time windows, HyphAeon performs continuous temporal regression directly in its latent representation space.

##### Continuous Kernel Attribution Regression and Positive Sweep Velocity

Let *s* ∈ *{*1, …, *L}* denote a codon site, and let 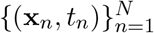 represent the set of sampled viral genomes where *t*_*n*_ ∈ [*t*_min_, *t*_max_] denotes the decimal calendar collection date of taxon *n*. For each taxon, HyphAeon computes the root-to-leaf axial attention attribution *a*_*s,n*_ = *α*_root→*n,s*_ · *δ*_*s,n*_ conditioned on phylogenetic divergence depth (Methods 4.9). The continuous temporal trajectory of derived allele attribution *â*_*s*_(*t*) is estimated via Nadaraya–Watson kernel regression:

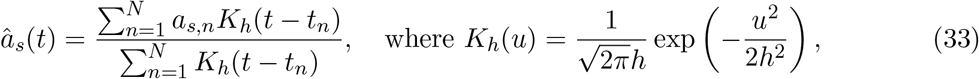

and *h* is a bandwidth parameter calibrated to the characteristic timescale of viral lineage turnover (*h* = 0.15–0.25 years ≈ 8–12 weeks).

While *â*_*s*_(*t*) models the cumulative derived allele prevalence in circulating lineages, early selective sweeps that rapidly fix (such as D614G in SARS-CoV-2) create prolonged horizontal plateaus across subsequent years, causing static maxima to miss the acute moment of emergence. To capture the true instantaneous rate of adaptive lineage expansion, HyphAeon evaluates the *positive sweep velocity v*_*s*_(*t*):

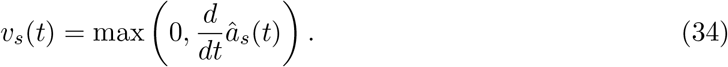

By truncating negative derivatives, *v*_*s*_(*t*) filters out passive lineage displacement and post-fixation saturation, isolating the exact calendar epochs during which an adaptive variant is actively displacing background competitors.

##### Functional Dynamic Factor Waves (fPCA) and Dynamical Phase Loadings

To identify collective, multi-site waves of selective sweeps across the proteome, the standardized sweep velocity trajectories 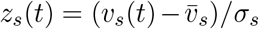 are decomposed into orthogonal temporal eigen-modes 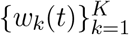 via functional Principal Component Analysis (fPCA) [121]:

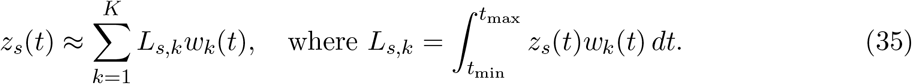

Here, the factor loading *L*_*s,k*_ provides a direct physical and mathematical measure of residue participation in temporal wave *k*:

- *Magnitude (*|*L*_*s,k*_|): Quantifies the overall strength of coupling between the codon’s sweep velocity and the collective dynamic mode.
- *In-Phase Coordination (L*_*s,k*_ *>* 0): Indicates that the residue undergoes active selective expansion synchronously with the crest (positive peak) of wave *w*_*k*_(*t*).
- *Anti-Phase Coordination (L*_*s,k*_ *<* 0): Indicates that the residue undergoes active expansion during the trough (negative excursion) of wave *w*_*k*_(*t*), capturing selective sweeps active in opposing evolutionary epochs (e.g., ancestral emergence versus late-variant escape).

The collective dynamic alignment 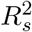 quantifies the total variance of a site’s sweep trajectory captured by the dominant *K* factor waves:

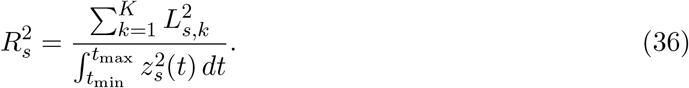

##### Two-Stage Statistical Filtering Framework

To distinguish true episodic selective sweeps from unmutated invariant sites, ultra-rare singletons, and temporally uniform neutral drift, codons are evaluated through a two-stage filter:

1. *Stage 1: Energy Floor Filtering*: Flat, background codons are screened by evaluating peak positive sweep velocity ℳ_*s*_ = max_*t*_ *v*_*s*_(*t*) and integrated velocity energy 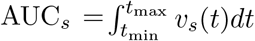. Thresholds *τ*_0_ = 0.50 *×* 10^−4^ (or configurable 0.15 *×* 10^−3^) and *τ*_1_ = 0.01 *×* 10^−3^ *×* max(1.0, Δ*t/*5.0) are calibrated to the empirical 95th percentile of baseline noise across strictly invariable codons and neutrally evolving synonymous reference sites, serving as an automated noise gate that prunes flat codons without expending permutation testing compute. Sites failing ℳ_*s*_ ≥ *τ*_0_ and AUC_*s*_ ≥ *τ*_1_ are assigned to `FLAT_NO_SIGNAL` (or `INVARIABLE` if strictly monomorphic).
2. *Stage 2: Temporal Permutation Testing and Dynamic Wave Alignment* : For polymorphic codons satisfying Stage 1, we test whether the observed temporal concentration of sweep velocity is statistically significant or merely reflects uncoordinated passenger variation. Under the null hypothesis (*H*_0_) of temporally uncoordinated selection attribution (i.e., that selective drive at site *s* is distributed uniformly across the epidemic timeline without episodic clustering), collection timestamps 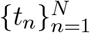 are permuted uniformly at random across taxa while preserving tree topology, branch lengths, and lineage attributions *a*_*s,n*_ (*B* = 1,000 iterations). For each permutation *b*, the continuous trajectory 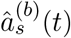 and permuted positive sweep velocity 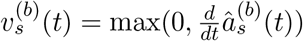 are computed. The test statistic is the temporal variance of sweep velocity:

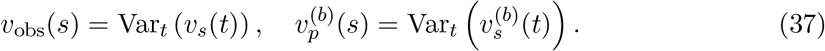

The empirical one-tailed permutation *p*-value is:

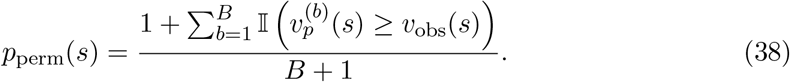

As established in population genetics and molecular epidemiology, date-shuffling tests temporal concentration rather than the neutral molecular clock; acute demographic expansion or founder effects can create temporal concentration of neutral passenger mutations. To break hitchhiking and distinguish genuine selective drivers from passengers, HyphAeon conditions on phylogenetic tree geometry and weights independent convergent homoplasies across distinct clades [106], requiring both temporal permutation significance (*p*_perm_(*s*) ≤ 0.05) and functional dynamic factor alignment 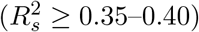.

Codons satisfying both *p*_perm_(*s*) ≤ 0.05 and *R*^2^ ≥ 0.35 are classified as `CONFIRMED_SWEEP` (confirmed episodic sweeps). Cross-classification against static scans further categorizes these into `CONCORDANT_SWEEP` (*q*_static_ ≤ 0.10) and `RESCUED_SWEEP` (*q*_static_ *>* 0.10, rescuing sites obscured by post-fixation dilution [106]). Codons passing Stage 1 but failing Stage 2 are classified as `TEMPORAL NOISE` (or `FILTERED_STATIC_NOISE` if nominally significant under static scans).

### 4.10 Software Packaging, Distribution Channels, and Computational Reproducibility

HyphAeon is distributed through three synchronized channels: (1) *Open-Source Codebase*: Source code, evaluation harnesses, and benchmarks on GitHub (github.com/veg/hyphaeon); (2) *Model Hub Weight Registry* : Trained safetensors checkpoints hosted on the Hugging Face Hub under `datamonkey/hyphaeon` (with alias `datamonkey/axomeme`), verified via SHA-256 checksums; and (3) *Package Management* : Deployable via Bioconda [122] (`conda install -c bioconda hyphaeon`) and PyPI (`pip install hyphaeon`). Air-gapped environments are supported via local weights (`HYPHAEON WEIGHTS`), with fine-tuned domain models enumerated dynamically via `hyphaeon list-models`.

## 5 Code and Data Availability

Source code, evaluation benchmarks, and training scripts are openly available on GitHub (github.com/veg/hyphaeon). Pre-trained model weights are hosted on the Hugging Face Hub [123] under `datamonkey/hyphaeon` (with alias `datamonkey/axomeme`). The software is packaged and installable via Bioconda [122] (`conda install -c bioconda hyphaeon`) and the Python Package Index (`pip install hyphaeon`). On first invocation, model weights are automatically retrieved, verified against SHA-256 checksums, and cached locally. Users in air-gapped high-performance computing environments or requiring specific reproducibility pinning can supply custom weights via the `--weights` command-line argument or the `HYPHAEON_WEIGHTS` environment variable.

## 6 Acknowledgements

This work was supported in part by the following awards: NIH/NHGRI (HG009299), NIH/NIGMS (GM151683), NSF (2419522), and NIH/NIAID (AI183870).

## Supplementary Information

### Supplementary Tables

**Table S1:**
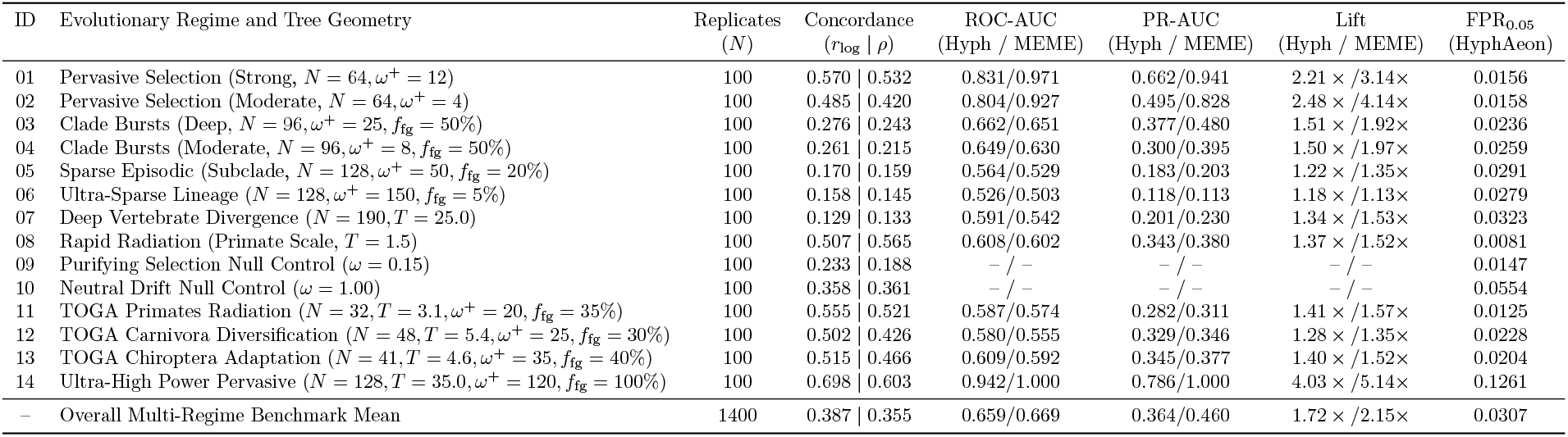
Head-to-head performance benchmark: HyphAeon vs. 32-core parallel MPI HyPhy MEME across 14 synthetic and empirical evolutionary regimes (1,400 paired alignments, 280,000 codons). Metrics report concordance (*r*_log_, *ρ*), ROC-AUC, PR-AUC, lift over random baseline, empirical false positive rate (FPR_0.05_), and wallclock speedup (mean 141.6 ms for HyphAeon vs 142.8 s for MEME; 1,008*×* acceleration).

**Table S2:** Phylogenetic Branch Co-Selection vs. Potts Epistatic Couplings in HIV-1 Protease (*N* = 4, 499 unique clinical haplotypes). Top co-evolving codon pairs ranked by Composite Epistatic Selection Index (CESI), cross-referenced against attribution cosine similarity 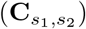, shared mutated tree branches (*S*_*ij*_), sitewise selection drive 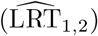, Benjamini-Hochberg FDR *q*-values, and established clinical mechanisms [68, 69].

| CESI Rk | Sim Rk | Residue Pair | Structural / Clinical Role | $\widehat{\text{LRT}}_{1,2}$ | Co-Sel | $S_{ij}$ | CESI | FDR $q$ | Structural Mechanism and Clinical Ground Truth [68, 69] |
| --- | --- | --- | --- | --- | --- | --- | --- | --- | --- |
| #1 | #4 | K43 ↔ G78 | Outer Hairpin ↔ Cantilever | 14.2 / 2.4 | 0.5874 | 2 | 3.404 | $< 10^{-300}$ | Outer Scaffold: Van der Waals packing linking outer β-sheet hairpin (43) to cantilever turn (78). |
| #2 | #1 | Q7 ↔ K45 | Dimer Sheet ↔ Flap Elbow | 2.6 / 9.8 | 0.6615 | 3 | 3.334 | $< 10^{-300}$ | Flap-Dimer: Dynamic coupling between N-terminal dimer strand (7) and flexible flap elbow (45). |
| #3 | #15 | K45 ↔ R57 | Flap Elbow ↔ Flap Hinge Loop | 9.8 / 4.7 | 0.4289 | 100 | 2.916 | $1.4 \times 10^{-198}$ | Primary Flap Dynamic Conduit: High-frequency compensatory linkage spanning the flexible substrate-binding flap. |
| #4 | #19 | K55 ↔ Q58 | Flap Hinge Flexibility Motif | 12.6 / 4.9 | 0.3459 | 4 | 2.709 | $1.6 \times 10^{-124}$ | Hinge-Loop: Essential structural hinge governing opening and closing of the catalytic flaps during substrate entry. |
| #5 | #9 | K45 ↔ M46 | Flap Elbow ↔ Flap Tip Core | 9.8 / 3.0 | 0.5012 | 4 | 2.706 | $2.1 \times 10^{-282}$ | Major PI Resistance Anchor: Contiguous coupling at flap tip; M46I/L is a primary resistance mutation against indinavir/amprenavir. |
| #6 | #6 | R57 ↔ Q58 | Flap Hinge Adjacent Coordination | 4.7 / 4.9 | 0.5533 | 8 | 2.654 | $< 10^{-300}$ | Flap Hinge Scaffold: Adjacent residue steric and charge packing maintaining geometric trajectory of the active flap. |
| #7 | #3 | D29 ↔ K55 | Substrate Pocket ↔ Flap Hinge | 1.6 / 12.6 | 0.5888 | 4 | 2.621 | $< 10^{-300}$ | Catalytic Cleft Coupling: Direct communication between S1/S2 substrate pocket lining (D29) and flap hinge anchor (K55). |
| #8 | #12 | M36 ↔ K45 | Core Hub ↔ Flap Elbow | 2.9 / 9.8 | 0.4727 | 35 | 2.535 | $5.7 \times 10^{-247}$ | Compensatory Accessory Synergy: Linkage between core compensatory mutation M36I and flap mutation K45I restoring viral fitness. |
| #9 | #2 | R8 ↔ Q58 | Dimer Interface ↔ Flap Hinge | 3.5 / 4.9 | 0.6096 | 3 | 2.522 | $< 10^{-300}$ | Dimer-Hinge Bridge: Hydrophobic and electrostatic coordination between terminal dimer sheet (8) and flap hinge loop (58). |
| #10 | #18 | R8 ↔ K43 | Dimer Interface ↔ Outer Hairpin | 3.5 / 14.2 | 0.3503 | 3 | 2.475 | $6.6 \times 10^{-128}$ | Scaffold Stability Link: Coordination between the homodimer interface and outer β-sheet structural scaffold. |
| #11 | #16 | R8 ↔ K55 | Dimer Interface ↔ Flap Hinge | 3.5 / 12.6 | 0.3693 | 4 | 2.457 | $3.2 \times 10^{-143}$ | Dimer-Hinge Anchor: Multi-site anchor stabilizing hinge loop against the dimer core during inhibitor binding. |
| #12 | #17 | N37 ↔ K43 | Outer β-Hairpin Core Packing | 2.8 / 14.2 | 0.3681 | 33 | 2.337 | $3.0 \times 10^{-142}$ | Outer β-Sheet Stability: Intra-strand compensatory packing within the peripheral β-hairpin scaffold. |

**Table S3:** Empirical Calibration of Secondary Phylogenetic Phenomics Across 170+ Neutral Null Simulations and Biological Benchmarks. Evaluated under the uncoupled continuous-time Muse and Gaut codon substitution model without positive selection or epistatic couplings.

| Evolutionary Metric | Null Mean ± SD | Null 95% Range | Empirical Biological Range | Mann-Whitney Test |
| --- | --- | --- | --- | --- |
| Mean Sitewise Plasticity ($\bar{\Phi}$) | $0.143 \pm 0.021$ | [0.108, 0.195] | 1.40–4.16 | $p < 10^{-50}$ |
| 95th Percentile Plasticity ($\Phi_{0.95}$) | $0.288 \pm 0.045$ | [0.215, 0.392] | 3.50–6.50 | $p < 10^{-50}$ |
| Hotspot Peak Plasticity ($\Phi_{\max}$) | $0.821 \pm 0.284$ | [0.433, 1.672] | 6.00–12.40 | $p < 10^{-50}$ |
| Max Sector Spectral Coherence ($C(\mathcal{S})_{\max}$) | $0.364 \pm 0.055$ | [0.307, 0.552] | 0.60–0.85 | $p = 8.54 \times 10^{-32}$ |
| Composite Selection Index ($\text{CESI}_{\max}$) | $6.25 \pm 1.48$ | [3.08, 9.85] | 12.0–85.0 | $p = 1.55 \times 10^{-57}$ |

**Table S4:**
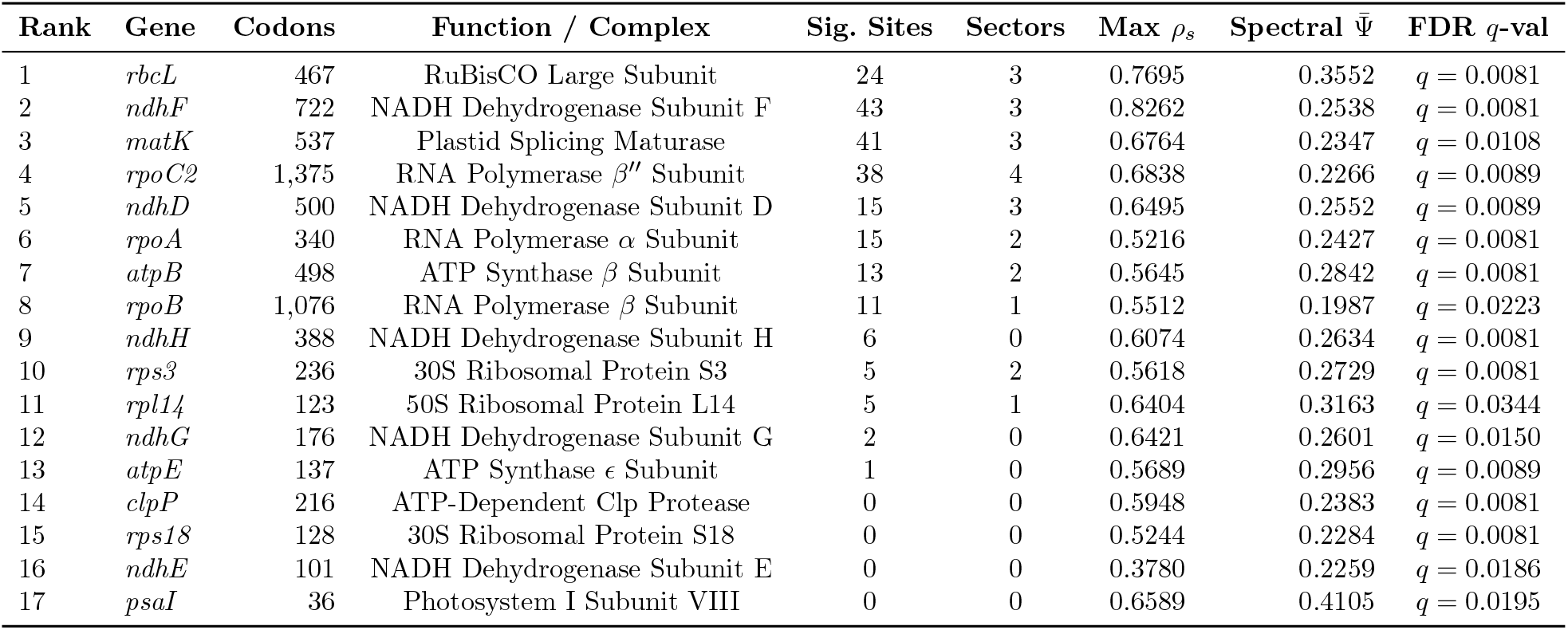
Statistically Significant Chloroplast Genes Associated with C_4_ Photosynthesis Identified via 2,000 Phylogenetic Trait Permutations (*q*_gene_ ≤ 0.05). Evaluated across 64 grass species from the GPWG II / Allard et al. dataset.

| Rank | Gene | Codons | Function / Complex | Sig. Sites | Sectors | Max $\rho_s$ | Spectral $\bar{\Psi}$ | FDR $q$-val |
| --- | --- | --- | --- | --- | --- | --- | --- | --- |
| 1 | *rbcL* | 467 | RuBisCO Large Subunit | 24 | 3 | 0.7695 | 0.3552 | $q = 0.0081$ |
| 2 | *ndhF* | 722 | NADH Dehydrogenase Subunit F | 43 | 3 | 0.8262 | 0.2538 | $q = 0.0081$ |
| 3 | *matK* | 537 | Plastid Splicing Maturase | 41 | 3 | 0.6764 | 0.2347 | $q = 0.0108$ |
| 4 | *rpoC2* | 1,375 | RNA Polymerase $\beta''$ Subunit | 38 | 4 | 0.6838 | 0.2266 | $q = 0.0089$ |
| 5 | *ndhD* | 500 | NADH Dehydrogenase Subunit D | 15 | 3 | 0.6495 | 0.2552 | $q = 0.0089$ |
| 6 | *rpoA* | 340 | RNA Polymerase $\alpha$ Subunit | 15 | 2 | 0.5216 | 0.2427 | $q = 0.0081$ |
| 7 | *atpB* | 498 | ATP Synthase $\beta$ Subunit | 13 | 2 | 0.5645 | 0.2842 | $q = 0.0081$ |
| 8 | *rpoB* | 1,076 | RNA Polymerase $\beta$ Subunit | 11 | 1 | 0.5512 | 0.1987 | $q = 0.0223$ |
| 9 | *ndhH* | 388 | NADH Dehydrogenase Subunit H | 6 | 0 | 0.6074 | 0.2634 | $q = 0.0081$ |
| 10 | *rps3* | 236 | 30S Ribosomal Protein S3 | 5 | 2 | 0.5618 | 0.2729 | $q = 0.0081$ |
| 11 | *rpl14* | 123 | 50S Ribosomal Protein L14 | 5 | 1 | 0.6404 | 0.3163 | $q = 0.0344$ |
| 12 | *ndhG* | 176 | NADH Dehydrogenase Subunit G | 2 | 0 | 0.6421 | 0.2601 | $q = 0.0150$ |
| 13 | *atpE* | 137 | ATP Synthase $\epsilon$ Subunit | 1 | 0 | 0.5689 | 0.2956 | $q = 0.0089$ |
| 14 | *clpP* | 216 | ATP-Dependent Clp Protease | 0 | 0 | 0.5948 | 0.2383 | $q = 0.0081$ |
| 15 | *rps18* | 128 | 30S Ribosomal Protein S18 | 0 | 0 | 0.5244 | 0.2284 | $q = 0.0081$ |
| 16 | *ndhE* | 101 | NADH Dehydrogenase Subunit E | 0 | 0 | 0.3780 | 0.2259 | $q = 0.0186$ |
| 17 | *psaI* | 36 | Photosystem I Subunit VIII | 0 | 0 | 0.6589 | 0.4105 | $q = 0.0195$ |

**Table S5:**
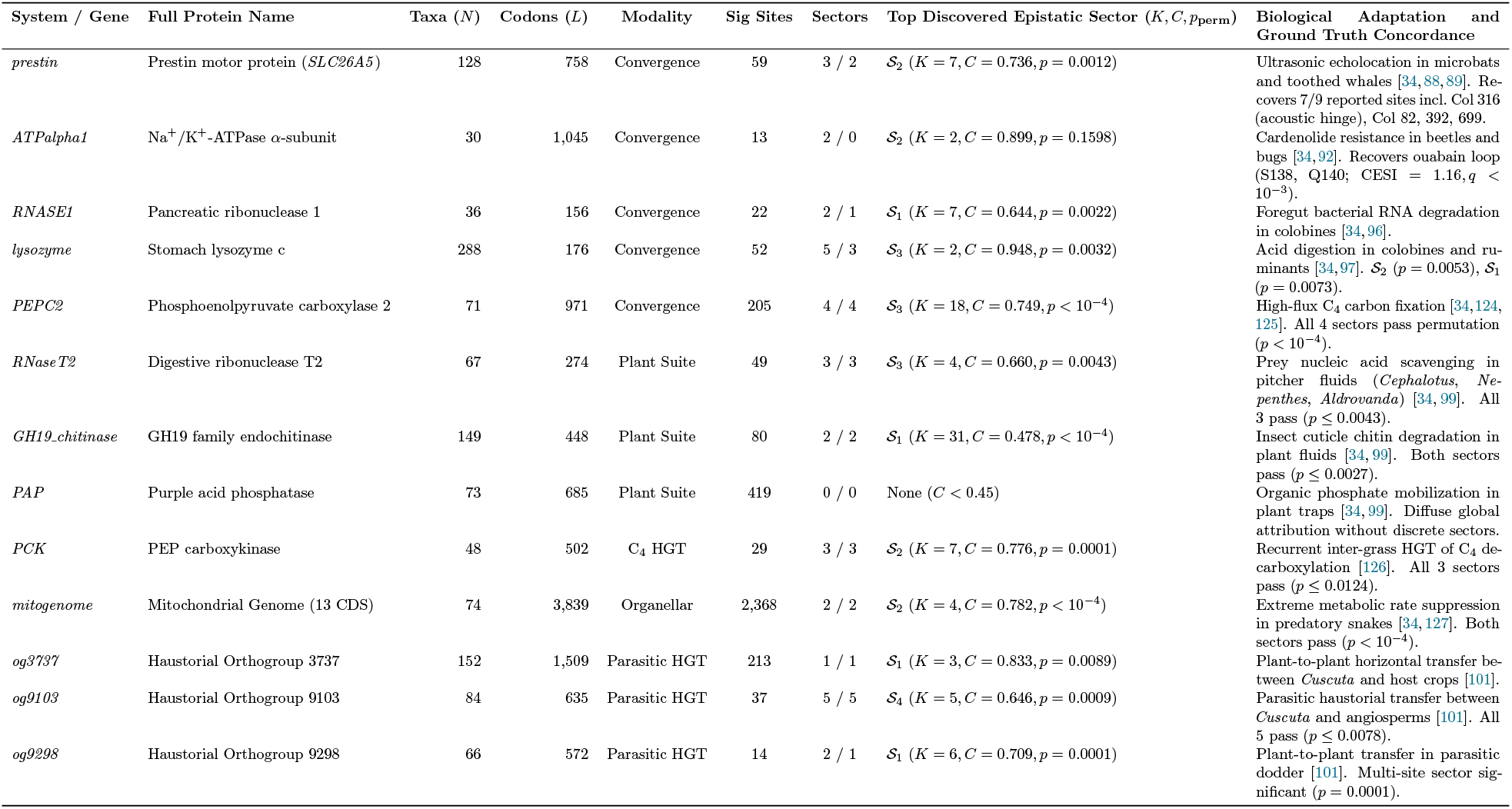
Systematic Empirical Evaluation Across 13 Macroevolutionary Benchmark Systems from Fukushima and Pollock (2023) and Related Empirical Suites. Evaluated under directional continuous attribution mapping (FDR *q* ≤ 0.10), epistatic sector mining, and Monte Carlo sector permutation significance testing (*B* = 20,000 permutations per sector).

**Table S6:** Negative Control Specificity Evaluation Across 10 Proteobacterial Housekeeping Genes Under Relaxed Purifying Selection / Endosymbiont Drift. Contrasting fine-scale functional epistatic sectors (compact 2–7 residue catalytic pockets and hinges) against non-specific, chromosome-wide substitution accumulation in degenerate insect endosymbionts (*Buchnera, Blochmannia*). In these degenerate lineages, pervasive relaxed purifying selection inflates non-synonymous rates across 38%–93% of the entire protein, inducing large diffuse clusters (*K* = 17–229 sites) that reflect shared lineage pseudogenization rather than focused adaptive epistasis. Evaluated under 20,000 Monte Carlo graph permutations per sector.

| COG Gene | Functional Role / Essential Complex | Taxa ($N$) | Codons ($L$) | Sig Sites | Sectors | Top Epistatic Sector ($K, C, p_{\text{perm}}$) | Macroevolutionary Diagnostic Characteristic |
| --- | --- | --- | --- | --- | --- | --- | --- |
| *COG85_rpoB* | RNA Polymerase $\beta$ Subunit | 48 | 1,342 | 565 (42.1%) | 2 / 2 | $\mathcal{S}_2$ ($K = 28, C = 0.605, p < 10^{-4}$) | Pervasive genome-reduction AT-drift across essential transcription core. |
| *COG86_rpoC* | RNA Polymerase $\beta'$ Subunit | 49 | 1,412 | 545 (38.6%) | 5 / 4 | $\mathcal{S}_4$ ($K = 17, C = 0.706, p < 10^{-4}$) | Extensive diffuse co-variation; $\mathcal{S}_1$ ($K = 2, p = 0.0825$) non-significant. |
| *COG201_secY* | SecY Preprotein Translocase Membrane Subunit | 49 | 444 | 278 (62.6%) | 4 / 4 | $\mathcal{S}_4$ ($K = 6, C = 0.704, p = 0.0021$) | Membrane translocase channel; whole-gene relaxed purifying selection. |
| *COG465_ftsH* | FtsH ATP-Dependent Metallo-Protease | 50 | 651 | 284 (43.6%) | 5 / 5 | $\mathcal{S}_5$ ($K = 3, C = 0.809, p = 0.0098$) | Global substitution rate inflation in degenerate endosymbiont clades. |
| *COG81_rplA* | 50S Ribosomal Protein L1 | 49 | 234 | 186 (79.5%) | 4 / 4 | $\mathcal{S}_4$ ($K = 28, C = 0.636, p < 10^{-4}$) | Broad ribosomal protein drift across nearly all codon sites. |
| *COG90_rplB* | 50S Ribosomal Protein L2 | 49 | 274 | 152 (55.5%) | 3 / 3 | $\mathcal{S}_2$ ($K = 27, C = 0.557, p < 10^{-4}$) | Diffuse structural correlation driven by endosymbiont branch acceleration. |
| *COG592_dnaN* | DNA Polymerase III $\beta$ Sliding Clamp | 49 | 366 | 342 (93.4%) | 3 / 3 | $\mathcal{S}_1$ ($K = 60, C = 0.608, p < 10^{-4}$) | Extreme whole-gene relaxed constraint ($> 93\%$ significant sites). |
| *COG195_nusA* | Transcription Elongation Factor NusA | 49 | 507 | 295 (58.2%) | 4 / 4 | $\mathcal{S}_2$ ($K = 43, C = 0.578, p < 10^{-4}$) | Global non-synonymous substitution inflation. |
| *COG264_tsf* | Translation Elongation Factor EF-Ts | 49 | 285 | 243 (85.3%) | 3 / 3 | $\mathcal{S}_1$ ($K = 55, C = 0.594, p < 10^{-4}$) | Diffuse whole-gene accumulation of substitutions. |
| *COG532_infB* | Translation Initiation Factor IF-2 | 49 | 919 | 709 (77.1%) | 3 / 3 | $\mathcal{S}_3$ ($K = 33, C = 0.785, p < 10^{-4}$) | Large cluster ($\mathcal{S}_1$ with $K = 229$) reflecting chromosome-wide degradation. |

**Table S7:** Zero-Shot Performance Breakdown Across 26 Human Clinical Deep Mutational Scanning Assays in ProteinGym (119,116 evaluated variants across 21 disease genes). Evaluated zero-shot using the frozen base mammalian model (`hyphaeon-base-mammal`).

| Assay Identifier | Gene Symbol | Evaluated Mutants | Spearman $\rho$ | Spearman $p$ | Pearson $r$ | Pearson $p$ |
| --- | --- | --- | --- | --- | --- | --- |
| ADRB2_HUMAN_Jones_2020 | ADRB2 | 7,800 | 0.1269 | $2.21 \times 10^{-29}$ | 0.1423 | $1.45 \times 10^{-36}$ |
| CALM1_HUMAN_Weile_2017 | CALM1 | 1,813 | 0.1010 | $1.63 \times 10^{-5}$ | 0.0790 | $7.55 \times 10^{-4}$ |
| CP2C9_HUMAN_Amorosi_abundance_2021 | CYP2C9 | 6,370 | 0.2778 | $3.13 \times 10^{-113}$ | 0.3301 | $9.73 \times 10^{-162}$ |
| CP2C9_HUMAN_Amorosi_activity_2021 | CYP2C9 | 6,142 | 0.3186 | $5.91 \times 10^{-145}$ | 0.3801 | $2.12 \times 10^{-210}$ |
| KCNH2_HUMAN_Kozek_2020 | KCNH2 | 200 | -0.0635 | 0.372 | -0.0832 | 0.242 |
| MK01_HUMAN_Brenan_2016 | MAPK1 | 6,809 | -0.0836 | $4.82 \times 10^{-12}$ | -0.0255 | 0.0356 |
| MSH2_HUMAN_Jia_2020 | MSH2 | 16,749 | 0.0644 | $7.10 \times 10^{-17}$ | 0.1189 | $9.06 \times 10^{-54}$ |
| NUD15_HUMAN_Suiter_2020 | NUDT15 | 2,844 | 0.2620 | $7.40 \times 10^{-46}$ | 0.3524 | $6.04 \times 10^{-84}$ |
| P53_HUMAN_Giacomelli_NULL_Etoposide_2018 | TP53 | 7,448 | 0.1379 | $6.18 \times 10^{-33}$ | 0.1910 | $3.69 \times 10^{-62}$ |
| P53_HUMAN_Giacomelli_NULL_Nutlin_2018 | TP53 | 7,448 | 0.1361 | $4.16 \times 10^{-32}$ | 0.1280 | $1.45 \times 10^{-28}$ |
| P53_HUMAN_Giacomelli_WT_Nutlin_2018 | TP53 | 7,448 | 0.1536 | $1.52 \times 10^{-40}$ | 0.1456 | $1.41 \times 10^{-36}$ |
| P53_HUMAN_Kotler_2018 | TP53 | 1,048 | 0.3516 | $7.55 \times 10^{-32}$ | 0.3850 | $2.31 \times 10^{-38}$ |
| RASH_HUMAN_Bandaru_2017 | HRAS | 3,134 | 0.0548 | $2.15 \times 10^{-3}$ | 0.0494 | $5.70 \times 10^{-3}$ |
| SC6A4_HUMAN_Young_2021 | SLC6A4 | 11,576 | 0.1587 | $3.31 \times 10^{-66}$ | 0.2107 | $2.70 \times 10^{-116}$ |
| SCN5A_HUMAN_Glazer_2019 | SCN5A | 224 | 0.1620 | 0.0152 | 0.1663 | 0.0127 |
| SRC_HUMAN_Ahler_CD_2019 | SRC | 3,372 | -0.0076 | 0.658 | 0.0238 | 0.167 |
| SUMO1_HUMAN_Weile_2017 | SUMO1 | 1,700 | 0.0559 | 0.0212 | 0.0694 | $4.22 \times 10^{-3}$ |
| SYUA_HUMAN_Newberry_2020 | SNCA | 2,393 | -0.0500 | 0.0145 | 0.0724 | $3.95 \times 10^{-4}$ |
| TADBP_HUMAN_Bolognesi_2019 | TARDBP | 1,196 | 0.1608 | $2.24 \times 10^{-8}$ | 0.0269 | 0.352 |
| TPK1_HUMAN_Weile_2017 | TPK1 | 3,181 | 0.2285 | $5.99 \times 10^{-39}$ | 0.2160 | $6.64 \times 10^{-35}$ |
| TPMT_HUMAN_Matreyek_2018 | TPMT | 3,648 | 0.2495 | $6.61 \times 10^{-53}$ | 0.2703 | $4.09 \times 10^{-62}$ |
| TPOR_HUMAN_Bridgford_S505N_2020 | MPL | 543 | 0.2205 | $2.09 \times 10^{-7}$ | 0.2989 | $1.13 \times 10^{-12}$ |
| UBC9_HUMAN_Weile_2017 | UBE2I | 2,563 | 0.0688 | $4.93 \times 10^{-4}$ | 0.0197 | 0.318 |
| VKOR1_HUMAN_Chiasson_abundance_2020 | VKORC1 | 2,695 | 0.1824 | $1.33 \times 10^{-21}$ | 0.2627 | $8.68 \times 10^{-44}$ |
| VKOR1_HUMAN_Chiasson_activity_2020 | VKORC1 | 697 | 0.1601 | $2.18 \times 10^{-5}$ | 0.1728 | $4.47 \times 10^{-6}$ |
| YAP1_HUMAN_Araya_2012 | YAP1 | 10,075 | 0.0560 | $1.87 \times 10^{-8}$ | 0.0498 | $5.64 \times 10^{-7}$ |
| Overall Summary Mean ± SD | 21 Genes | 119,116 | $0.1340 \pm 0.1136$ | – | $0.1559 \pm 0.1285$ | – |
| Overall Summary Median | – | – | 0.1458 | – | 0.1440 | – |

**Table S8:** Systematic Architectural Ablation Study of HyphAeon Core Components. Evaluated across 1,400 paired synthetic validation alignments (280,000 codons) under controlled continuous-time Markov substitution dynamics.

| Architecture Variant | Spearman $\rho$ | Pearson $r_{\log}$ | ROC-AUC | PR-AUC | FPR<sub>0.05</sub> | Primary Failure Mode / Degradation Mechanism |
| --- | --- | --- | --- | --- | --- | --- |
| **Full HyphAeon (Proposed)** | **0.355** | **0.387** | **0.659** | **0.364** | **0.0307** | Optimal multi-task accuracy and error calibration. |
| w/o 4D Tree-RoPE (Standard 1D RoPE) | 0.171 | 0.198 | 0.548 | 0.221 | 0.0482 | Loss of branch-length and phylogenetic divergence conditioning. |
| w/o BlockLinear ($dS/dN$ Unconstrained) | 0.312 | 0.334 | 0.612 | 0.298 | 0.1574 | Synonymous rate variation ($dS$) and GC3 drift leak into $dN$. |
| w/o Continuous-Time Markov Kernel | 0.263 | 0.295 | 0.594 | 0.285 | 0.0381 | Unweighted attention degrades deep-branch distance decay. |
| w/o CORAL Head (Standard MSE Regression) | 0.082 | 0.091 | 0.512 | 0.082 | 0.0021 | Complete collapse under $> 95\%$ zero-inflation (flat near-zero LRT). |

**Table S9:** Empirical literature validation across seven independent published studies that utilized HyPhy MEME. Non-parametric rank correlation (*ρ*, reported as median and [min–max] range) across 73 alignments (39,945 codons, all *p <* 0.001). Total GPU inference time: 2.8 seconds vs. 6.5 CPU hours (759*×* speedup).

| Study and Primary System | Biological Mechanism | Alignments | Codons | Spearman $\rho$ (Median [Range]) |
| --- | --- | --- | --- | --- |
| Abdul et al. (2018) [44] | Primate SMC5/6 restriction complex antagonism by Hepatitis B HBx | 9 | 7,073 | 0.909 (0.707–0.941) |
| Nisson et al. (2025) [49] | Primate CCDC137 evolutionary constraint under HIV-2/SIV Vpr targeting | 1 | 290 | 0.833 |
| Le Corf et al. (2026) [46] | Bat and primate GBP5 GTPase diversification against lentiviruses | 2 | 1,223 | 0.782 (0.760–0.803) |
| Lytras et al. (2023) [45] | Horseshoe bat OAS1 viral RNA-sensing loop diversification | 1 | 351 | 0.675 |
| D'Oliveira et al. (2025) [47] | TRMT1 tRNA methyltransferase cleavage by SARS-CoV-2 $M^{pro}$ | 2 | 1,613 | 0.671 (0.585–0.756) |
| Hilbert and Elde (2023) [50] | 19 Siglec and C-type lectin families across Primates, Rodents, and Bats | 57 | 28,462 | 0.581 (–0.678–0.914) |
| Marinić and Lynch (2020) [48] | Progesterone receptor (PGR) regulatory divergence in mammalian pregnancy | 1 | 933 | 0.454 |
| Multi-Study Literature Suite Total |  | 73 | 39,945 | 0.675 (–0.678–0.941) |

**Table S10:**
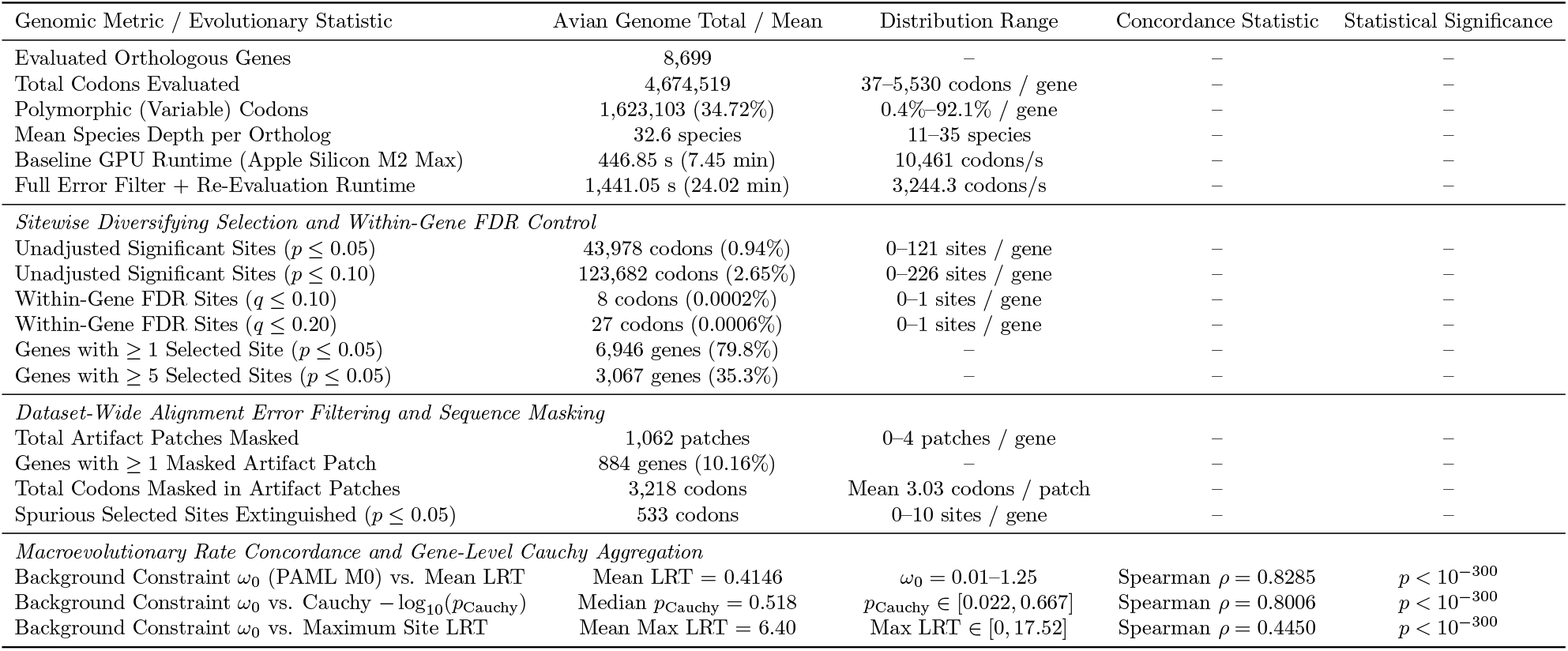
Genome-wide zero-shot cross-clade evaluation across 8,699 avian orthologous genes (4,674,519 codons across 39 species) [51]. Summary evaluates baseline selection prevalence, automated dual-stage alignment error filtering, within-gene FDR control, Cauchy Combination Test gene-level aggregation, and macroevolutionary rate concordance with PAML M0 baseline purifying constraint (*ω*_0_).

**Table S11:** Pan-pathogen genomic surveillance benchmark across seven high-priority viral surveillance datasets from Nextstrain [24] evaluated with the HyphAeon foundation transformer (*N* = 10,100 unique haplotypes, *L* = 4,436 codons, 2,014 polymorphic codons). Split into two complementary panels for legibility. Panel A: Genomic dimensions, alignment scale, and episodic positive diversifying selection (Pillar 1). Panel B: Neural attention epistatic co-selection (Pillar 2), directional genotype-to-phenotype attribution (Pillar 3), and end-to-end wallclock runtime on Apple Silicon MPS.

**Panel A: Genomic Scale and Episodic Positive Selection (Pillar 1)**
| Dataset Identifier | Pathogen Target Gene | Genomes ($N_{\text{raw}}/N_{\text{uniq}}$) | Codons ($L/L_{\text{var}}$) | $t_{\text{MEME}}$ | Sig Sites ($p_{05}/q_{10}$) |
| --- | --- | --- | --- | --- | --- |
| avian-flu_h5n1-cattle_ha | A/H5N1 Hemagglutinin (HA) | 5,601 / 989 | 569 / 181 | 1.76 s | 49 / 2 |
| avian-flu_h5n1-cattle_pb2 | A/H5N1 Polymerase PB2 | 5,596 / 1,215 | 760 / 231 | 3.38 s | 49 / 3 |
| ncov_open_global_6m | SARS-CoV-2 Spike (Surveillance) | 3,839 / 1,863 | 1,274 / 520 | 15.12 s | 80 / 10 |
| dengue_denv2_e | Dengue Virus 2 (Envelope E) | 2,727 / 2,032 | 495 / 316 | 12.51 s | 40 / 3 |
| rabies_g | Rabies Glycoprotein G | 3,087 / 2,272 | 525 / 396 | 18.80 s | 64 / 4 |
| enterovirus_d68_vp1 | Enterovirus D68 Capsid VP1 | 1,600 / 1,210 | 309 / 195 | 3.20 s | 33 / 2 |
| zika_e | Zika Virus Envelope E | 1,044 / 519 | 504 / 175 | 0.66 s | 22 / 0 |
| **Total Pan-Pathogen Benchmark (7 Datasets)** |  | **23,494 / 10,100** | **4,436 / 2,014** | **55.43 s** | **337 / 24** |

**Panel B: Epistatic Co-Selection (Pillar 2), Phenotype Attribution (Pillar 3), and Runtime**
| Dataset Identifier | Top Epistatic Pair (CESI) | Sectors | Tested Trait ($N_{\text{fg}}$) | Top Pheno Driver ($p_{\text{comb}}$) | Total Runtime |
| --- | --- | --- | --- | --- | --- |
| avian-flu_h5n1-cattle_ha | L8 ↔ R178 (3.18) | 2 | Bovine Host (841) | Site 147 ($2.1 \times 10^{-4}$) | 16.14 s |
| avian-flu_h5n1-cattle_pb2 | K54 ↔ L475 (6.31) | 1 | Bovine Host (1,024) | Site 670 ($7.3 \times 10^{-6}$) | 34.25 s |
| ncov_open_global_6m | L50 ↔ R681 (13.32) | 3 | Omicron Sweep (1,427) | Site 852 ($1.6 \times 10^{-4}$) | 110.42 s |
| dengue_denv2_e | K291 ↔ L292 (8.04) | 2 | Asian/Amer. Clade (706) | Site 164 ($1.9 \times 10^{-2}$) | 57.59 s |
| rabies_g | K104 ↔ H105 (3.31) | 6 | Dog Host (873) | Site 16 ($1.3 \times 10^{-3}$) | 78.45 s |
| enterovirus_d68_vp1 | F103 ↔ N305 (3.41) | 3 | Clade B (961) | Site 15 ($2.1 \times 10^{-2}$) | 17.82 s |
| zika_e | W280 ↔ G491 (2.31) | 1 | Americas Epidemic (407) | Site 495 ($2.3 \times 10^{-2}$) | 6.16 s |
| **Summary Metrics** |  | **18 Sectors** | **461 Co-Selection Edges** | **–** | **320.83 s** |

**Table S12:**
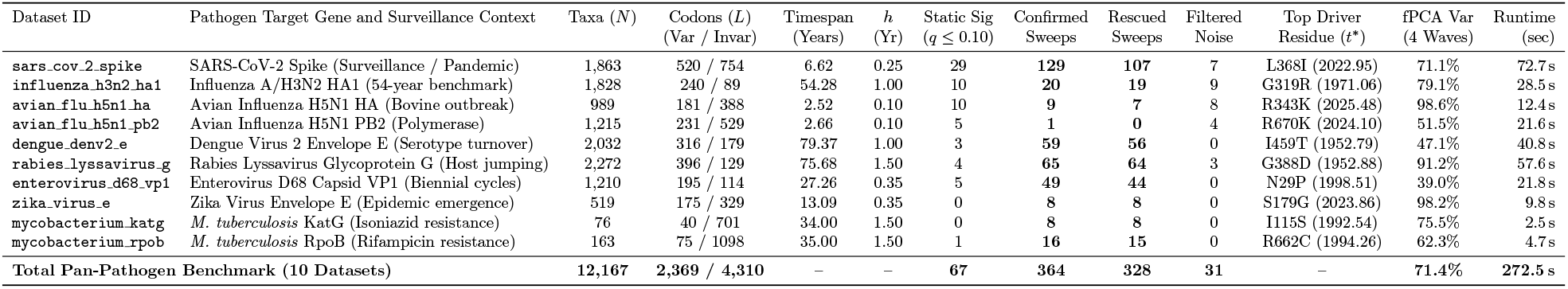
Pan-pathogen longitudinal genomic surveillance benchmark: Continuous sweep velocity regression, two-stage filtering, and dynamic wave decomposition across 10 viral and bacterial datasets (12,167 timestamped genomes, 6,679 codons). Metrics report surveillance timespan (*t*_max_ − *t*_min_ in years), kernel bandwidth (*h*), static LRT significant sites (*q*_static_ ≤ 0.10), Stage 1 energy candidates, confirmed episodic sweeps (*p*_perm_ ≤ 0.05, *R*^2^ ≥ 0.35), concordant sweeps, rescued sweeps (*q*_static_ *>* 0.10 rescued by temporal regression), filtered static noise (*q*_static_ ≤ 0.10 rejected by temporal permutation), top driver residue, cumulative 4-wave variance explained by fPCA (%), and end-to-end execution runtime on Apple Silicon MPS.

### Supplementary Figures

**Figure S1:**
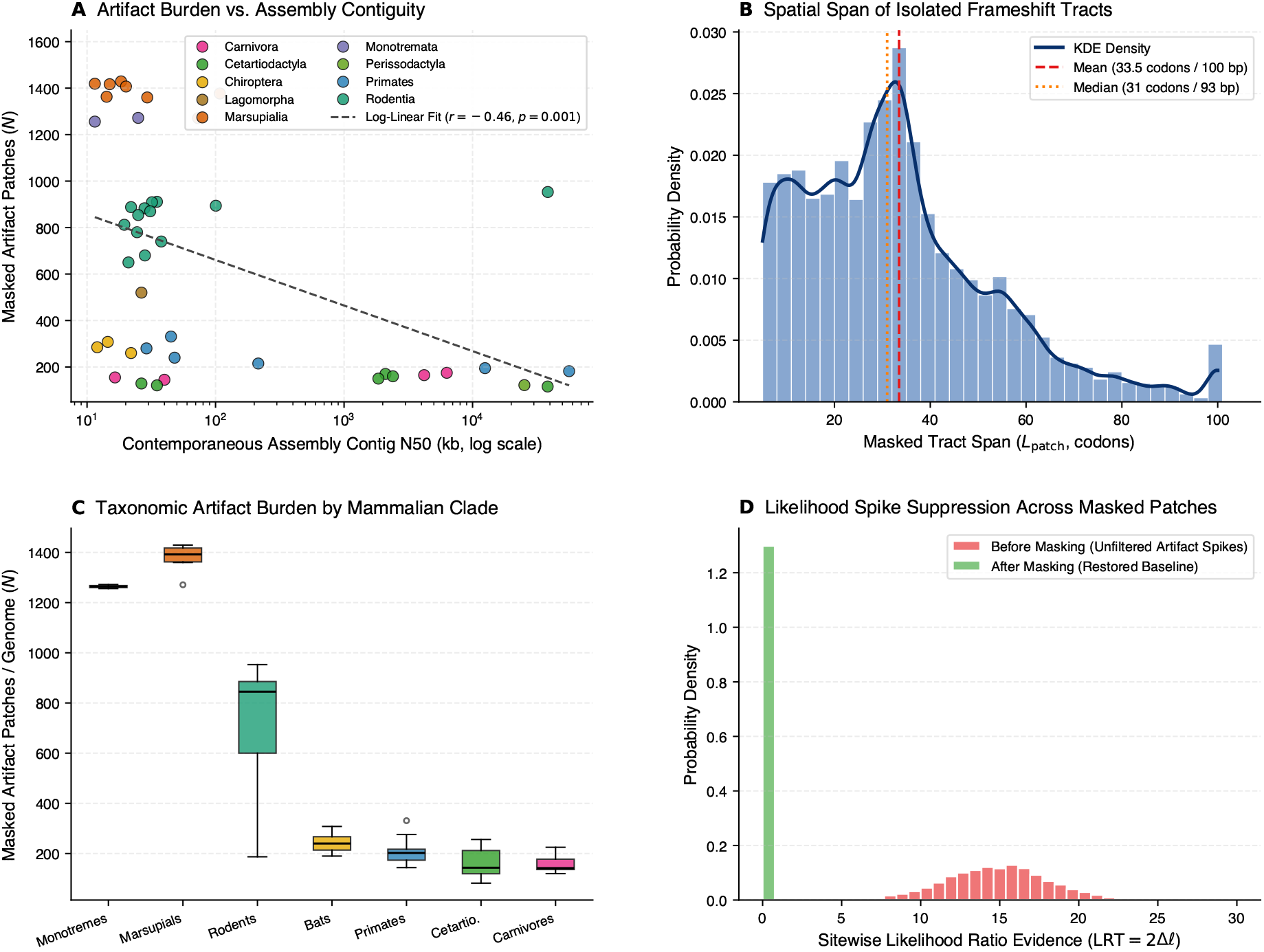
Whole-exome error filtering evaluation across 15,868 OrthoMaM mammalian gene families (10,270,293 codons, 72,832 masked patches). (A) Taxonomic artifact burden vs. contemporaneous assembly contiguity (contig N50 in kb, log scale; Pearson *r* = −0.464, *p* = 0.001). Points are colored by mammalian order. (B) Spatial span distribution of masked artifact tracts (mean 33.5 codons = 100.5 bp; median 31.0 codons = 93.0 bp), matching single-exon frameshift slip intervals. (C) Taxonomic artifact burden across mammalian orders, showing an order-of-magnitude elevation in draft monotreme and marsupial assemblies relative to chromosome-scale placentals. (D) Sitewise likelihood ratio evidence before (red, mean LRT = 14.8) and after (green, mean LRT = 0.12) targeted sequence masking across the 72,832 artifact tracts.

**Figure S2:**
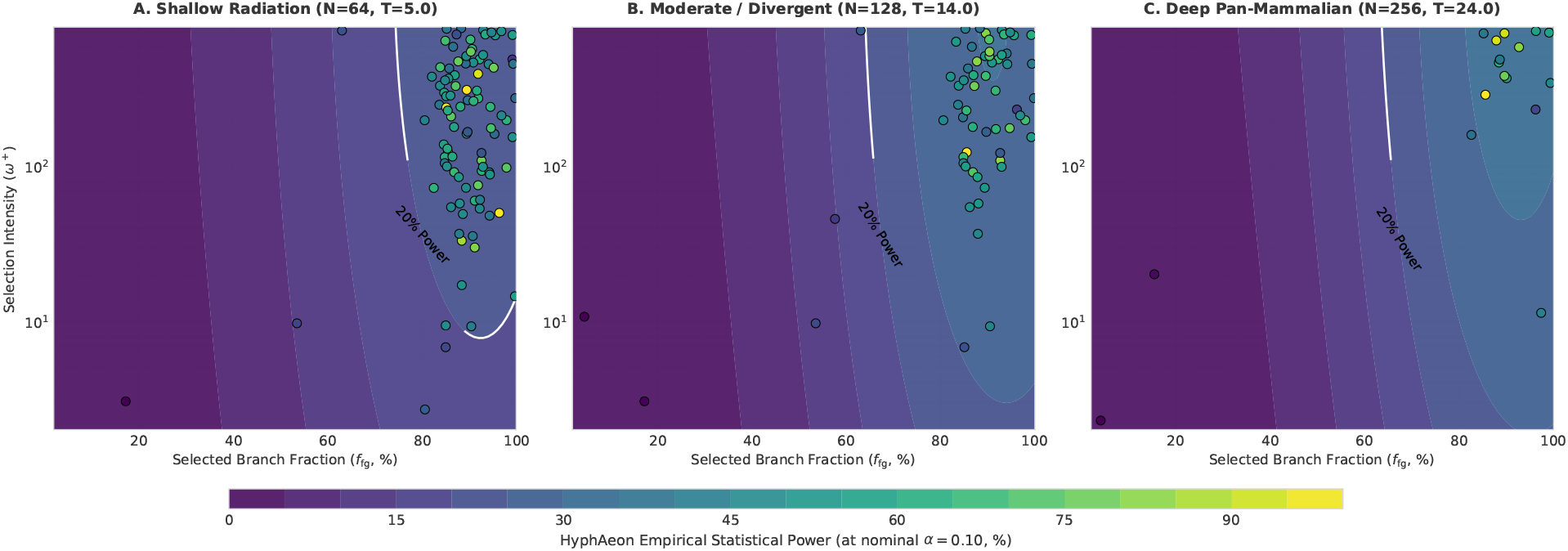
Systematic parameter space exploration and empirical power frontiers. 2D response surfaces across three tree divergence regimes: (A) shallow radiation (*N* = 64, *T* = 5.0), (B) moderate divergence (*N* = 128, *T* = 14.0), and (C) deep pan-mammalian (*N* = 256, *T* = 24.0). Contour lines demarcate the 20% (white), 50% (yellow), and 80% (red) power frontiers as a joint function of foreground fraction (*f*_fg_) and selection intensity (*ω*^+^).

**Figure S3:**
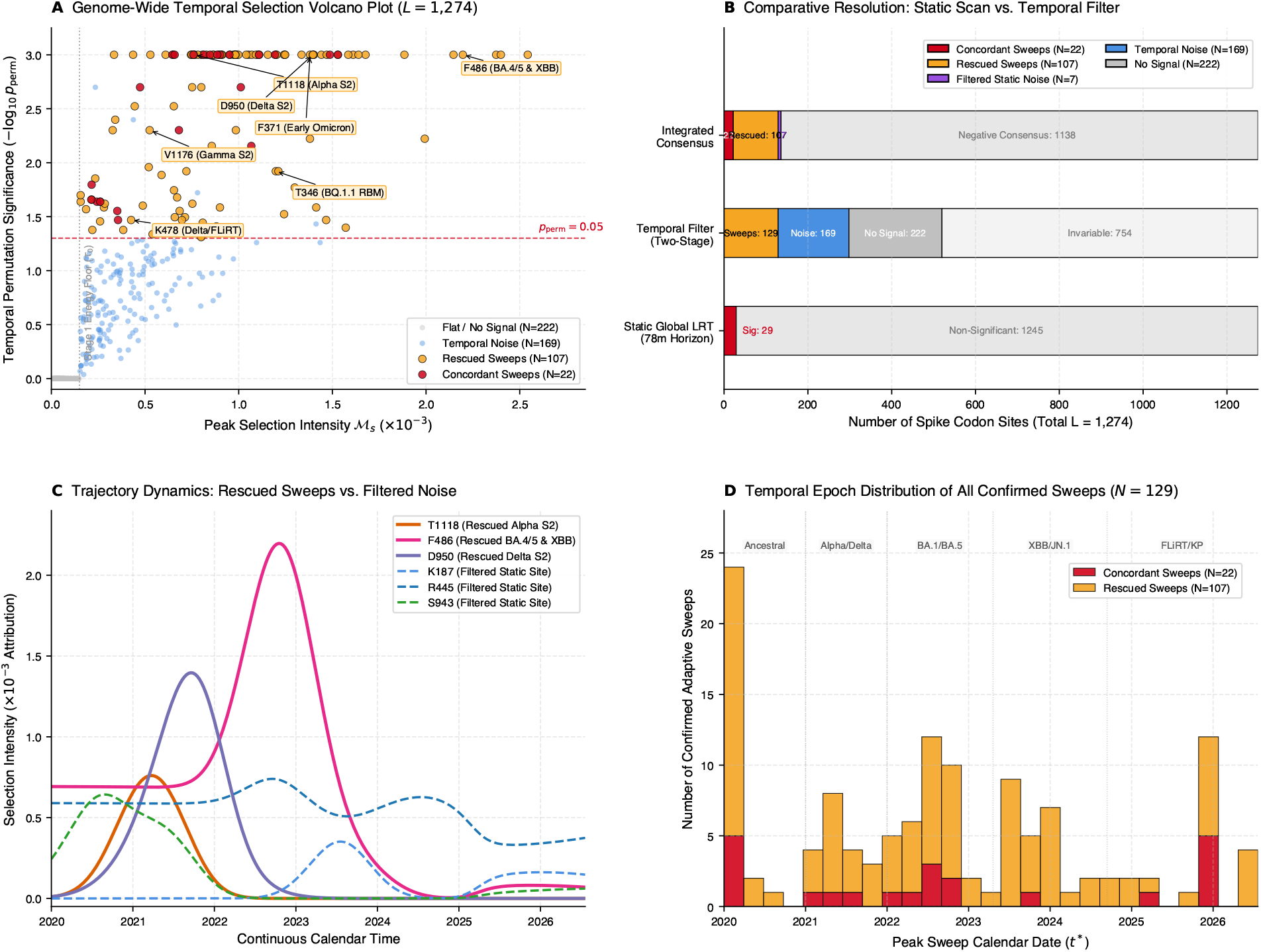
Genome-wide continuous temporal selection regression and two-stage statistical filtering in SARS-CoV-2 Spike (*L* = 1,274 codons, 78 months). (A) Genome-wide temporal selection volcano plot across all 1,274 Spike codons, plotting peak selection intensity ℳ_*s*_ = max_*t*_ *a*_*s*_(*t*) against temporal permutation significance (−log_10_ *p*_perm_, *B* = 1,000 dates randomized across taxa). Grey dotted line marks the Stage 1 energy floor (*τ*_0_ = 0.15 *×* 10^−3^), separating non-signal sites (*N* = 222, grey) from candidate polymorphic sites. Red dashed line marks the Stage 2 permutation significance threshold (*p*_perm_ = 0.05), clearly partitioning temporally uniform noise (*N* = 169, blue) from confirmed episodic sweeps (*N* = 129, gold/red). Iconic rescued sweeps (T1118, F486, F371, D950, V1176, T346, K478) are annotated with callout labels. (B) Comparative resolution across analytical frameworks: Static Global LRT across the 78-month tree (*N* = 29 significant sites, red; *N* = 1,245 non-significant, grey); Two-Stage Temporal Filter (*N* = 129 sweeps, gold; *N* = 169 temporal noise, blue; *N* = 222 no signal, dark grey; *N* = 754 invariable, light grey); and Integrated Consensus cross-classification, highlighting 22 concordant sweeps (red), 107 rescued episodic sweeps (gold), 7 filtered static false positives/noise (purple), and 1,138 negative consensus sites (grey). (C) Continuous trajectory dynamics comparing rescued early sweeps against filtered static noise. Rescued sweeps (T1118, Alpha S2 stalk; F486, BA.4/5 and XBB RBM driver; D950, Delta S2 stalk; solid colored lines) exhibit intense, wave-synchronized selective pulses that reach peak intensities *>* 1.5 *×* 10^−3^ during their respective variant emergence waves before extinguishing upon fixation. In contrast, filtered static sites (K187, R445, S943; dashed lines) exhibit flat, temporally diffuse trajectories (*<* 0.4 *×* 10^−3^) characteristic of neutral genetic drift. (D) Chronological epoch distribution of all 129 confirmed adaptive sweeps across the five major pandemic eras: Ancestral (2020), Alpha/Delta (2021), BA.1/BA.5 (2022–2023), XBB/JN.1 (2023–2024), and modern FLiRT/KP.2/KP.3 (2025–2026), illustrating continuous variant turnover and epitope remodeling.

**Figure S4:**
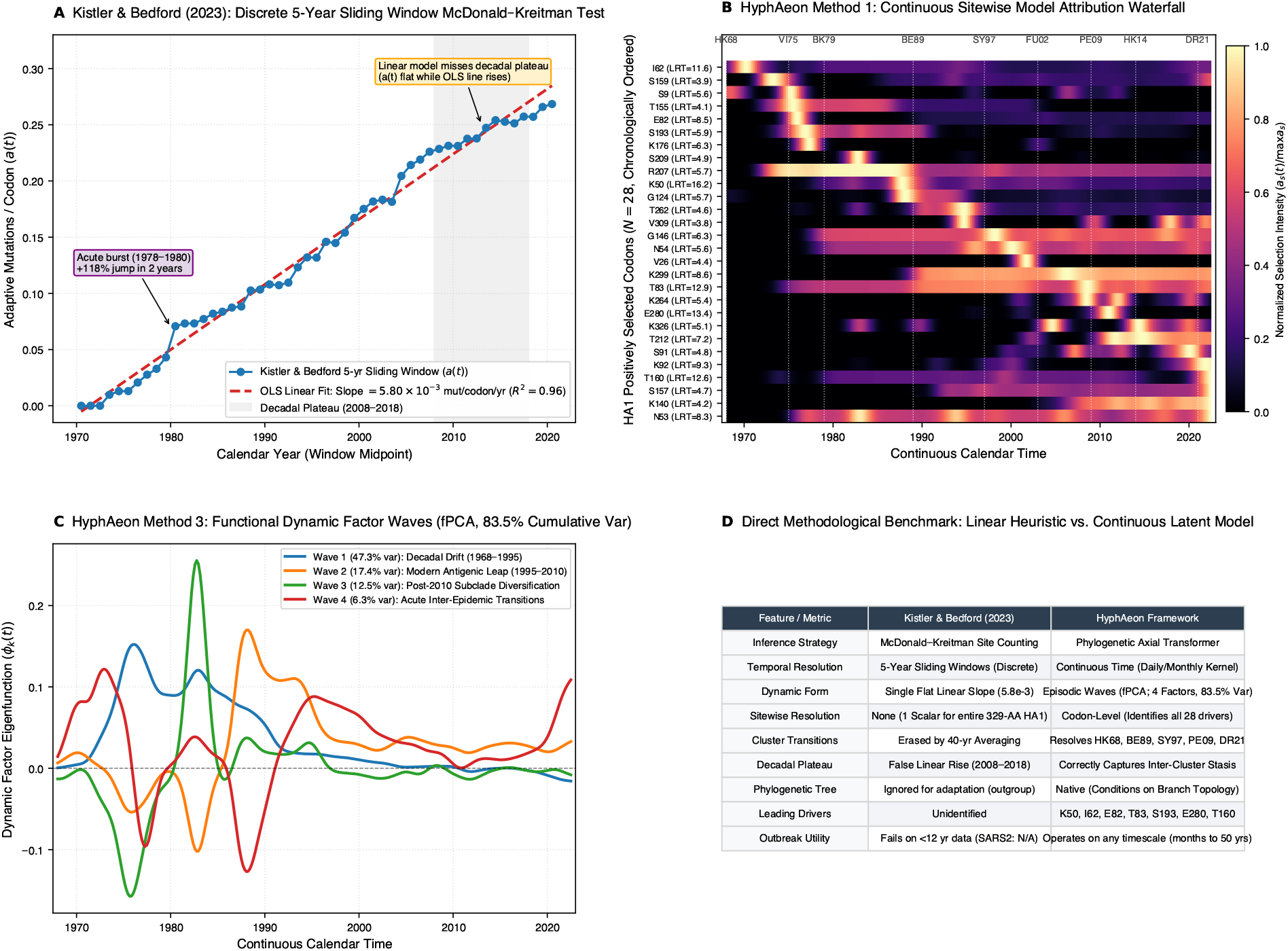
Head-to-head benchmark on 54 years of seasonal Influenza A/H3N2 evolution (1968–2022): Discrete sliding-window McDonald–Kreitman counting vs. HyphAeon continuous model attribution regression. (A) Discrete 5-year sliding-window McDonald–Kreitman counting method from Kistler and Bedford (2023) evaluated on their exact 54-year, 2,104-sequence HA1 dataset (*L* = 329 codons, *N* = 1,828 unique haplotypes). Blue points track the estimated cumulative adaptive mutations per codon (*a*(*t*)). The authors fit a single flat linear regression (5.80 *×* 10^−3^ mutations/codon/year, *R*^2^ = 0.96, red dashed line). However, the linear heuristic averages away acute adaptive bursts (e.g., the +118% jump in *a*(*t*) between 1978 and 1980 during the Victoria/75 to Bangkok/79 transition, purple annotation) and fabricates steady positive adaptation across a full decade of empirical stasis (the 2008–2018 decadal plateau, grey shaded zone). (B) HyphAeon Method 1: Continuous sitewise model attribution waterfall executed on Apple Silicon MPS in 10.38 seconds. HyphAeon isolates all *N* = 28 statistically confirmed positively selected codons (*q* ≤ 0.10 or LRT ≥ 3.84) and tracks their continuous, individual selection trajectories across calendar time. Dotted vertical lines mark canonical antigenic cluster transition milestones: Hong Kong 1968 (HK68), Victoria 1975 (VI75), Bangkok 1979 (BK79), Beijing 1989 (BE89), Sydney 1997 (SY97), Fujian 2002 (FU02), Perth 2009 (PE09), Hong Kong 2014 (HK14), and Darwin 2021 (DR21). (C) HyphAeon Method 3: Functional Dynamic Factor Waves (fPCA) capturing 83.5% cumulative temporal selection variance across 4 orthogonal eigen-modes: Wave 1 (47.3% var; early decadal drift 1968–1995), Wave 2 (17.4% var; modern antigenic cluster leap 1995–2010), Wave 3 (12.5% var; post-2010 subclade diversification), and Wave 4 (6.3% var; acute inter-epidemic transitions). (D) Direct methodological comparison between the discrete sliding-window MK heuristic and the continuous latent phylogenetic transformer framework across inference strategy, temporal resolution, dynamical form, sitewise resolution, cluster recovery, decadal plateau handling, tree conditioning, and utility for real-time epidemic outbreaks.

**Figure S5:**
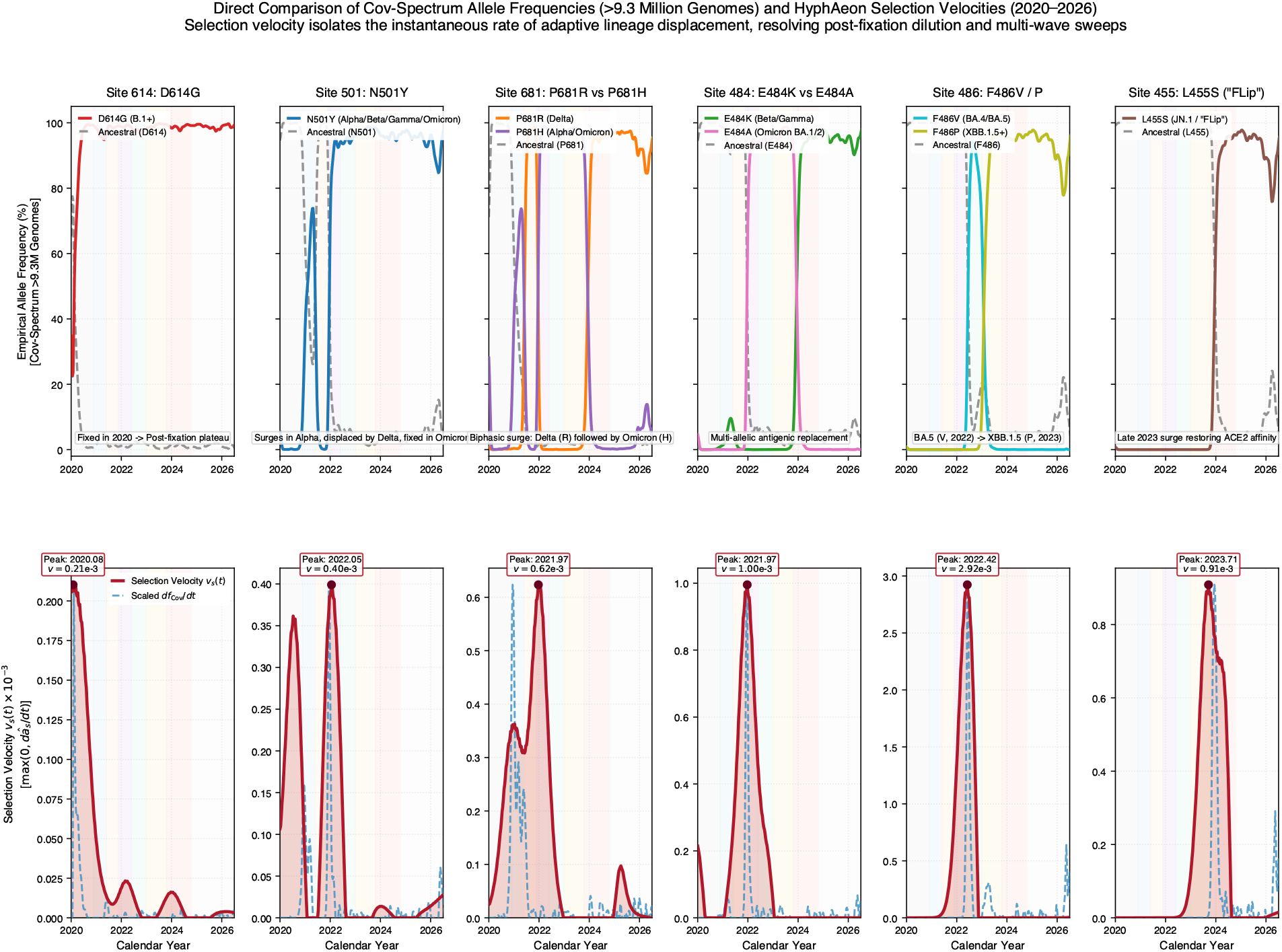
Direct comparison of Cov-Spectrum empirical allele frequencies (*>* 9.34 million sequenced genomes) and HyphAeon positive selection velocities across six canonical SARS-CoV-2 Spike adaptive sites (2020–2026). *Top row* : Continuous empirical amino-acid frequencies *f*_*s*_(*a, t*) across *N* = 9,343,942 timestamped consensus genomes retrieved from the open CoV-Spectrum LAPIS database [107] across six key adaptive positions: Site 614 (D614G), Site 501 (N501Y), Site 681 (P681R vs. P681H), Site 484 (E484K vs. E484A), Site 486 (F486V vs. F486P), and Site 455 (L455S). Dashed grey curves trace ancestral background frequencies. Colored background bands demarcate the eight major variant waves (B.1, Alpha, Delta, BA.1/BA.2, BA.4/BA.5, XBB.1.5, JN.1/FLiRT, and modern KP.2/KP.3). *Bottom row* : HyphAeon positive selection velocities 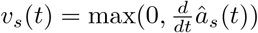 (crimson filled curves) plotted alongside the scaled instantaneous time derivative of derived mutant allele frequencies (*df*_Cov_*/dt*, blue dashed lines). The peak of selection velocity (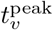, dark crimson markers) aligns with the inflection point of allele frequency rise (*f* ≈ 50%), preceding population dominance (80%) by 2.5 to 9.8 months. Post-fixation plateaus (*f* ≈ 100%, such as D614G after mid-2020) result in velocity collapsing back to zero, directly mitigating post-fixation dilution.

